# MnSOD Diversifies, while FeSOD Dominates Superoxide Defense in Cyanobacteria

**DOI:** 10.64898/2026.10.09.757883

**Authors:** Monis Athar Khan, Anne Durand, Anne-Soisig Steunou, Solène Leleux, Corinne Cassier-Chauvat, Franck Chauvat, Soufian Ouchane

**Affiliations:** Université Paris-Saclay, CEA, CNRS, Institute for Integrative Biology of the Cell (I2BC), Gif-sur-Yvette, France

**Keywords:** Cyanobacteria, Superoxide dismutases, Photooxitative stress, SOD evolution, Lipoprotein, thylakoid MnSOD

## Abstract

Cyanobacteria are unique among prokaryotes in possessing photosynthetic thylakoid membranes enclosing an internal lumen, where superoxide production is expected to be substantial. Thylakoid-associated superoxide dismutases (SODs) would seem the obvious first line of defense, yet many cyanobacteria rely instead on a single soluble cytoplasmic enzyme, raising the question of what role thylakoid-associated SOD isoforms actually play. Using *Cyanothece* sp. PCC 7425, which encodes both a cytoplasmic FeSOD and a thylakoid-associated MnSOD, we combined sequence analysis, structural, genetics, and biochemical approaches to characterise FeSOD and MnSOD cellular localisation and assess each enzyme’s contribution to superoxide defense. Cyanobacterial MnSODs display diverse N-terminal architectures, consistent with soluble, transmembrane, or lipid-anchored forms, yet contribute only modestly to superoxide defense in *Cyanothece* PCC 7425 and, very likely, across cyanobacteria broadly. Soluble FeSOD instead provides the major line of defense, showing that thylakoid association does not guarantee physiological importance. This predominance of a cytoplasmic enzyme highlights the central role of cytoplasmic superoxide detoxification, consistent with superoxide being generated mainly at the PSI acceptor side, and extends to the chloroplast stroma of plants. Our findings support a model in which SOD distribution reflects metal availability, enzyme properties, and subcellular organisation : FeSOD protects the compartment harbouring the main targets of PSI-associated superoxide, whereas MnSOD is retained where thylakoid association offers additional local protection, but becomes dispensable when the local ROS burden is limited. Spatial matching between superoxide production, antioxidant localisation, and target vulnerability may thus shape the diversification and maintenance of antioxidant systems across cyanobacteria and chloroplasts.

## Introduction

Superoxide (O_2_^.-^) anion is a ubiquitous byproduct of electron transfer reactions arising from the one electron reduction of molecular (di)oxygen [1]. In plants, these reactive oxygen species (ROS)-generating processes take place in two distinct organelles: mitochondria and chloroplast, whereas in cyanobacteria, they are associated with the cytoplasmic and thylakoid membranes. In the respiratory chain, electron leakage from complexes I (NADH dehydrogenase) and III (cytochrome *b_6_f*) leads to the formation of O_2_^.-^ [2]. In the photosynthetic chain, excess light energy generates O_2_^.-^ and other ROS, driving photooxidative damage. Superoxide is thought to originate at multiple sites [3] within the photosynthetic apparatus, including photosystem II (PSII), the cytochrome *b_6_f* complex, and photosystem I (PSI), with the primary production site being located at the acceptor side of PSI [4]. Under excess light generating excess reducing power, oxygen serves ineluctably as an alternative electron acceptor at PSI, resulting in the formation of O_2_^.-^ directly in the cytoplasm. While it was also reported that O_2_^.-^ anion radical could be formed by the one electron reduction of oxygen by the semiquinones at the acceptor side (lumen) of PSII and in the *b_6_f* complex [3]. In plant chloroplasts, the acceptor side of PSI is considered a major site of O_2_^.-^ generation through the Mehler reaction. Consequently, superoxide is produced predominantly on the stromal side of the thylakoid membrane [4].

Superoxide dismutases (SODs) are a primary component of the antioxidant defense system, catalyzing the conversion of O_2_^.-^ into molecular oxygen and hydrogen peroxide [1]. SODs are metalloenzymes classified according to their metal cofactors, with Fe and Mn containing SODs (FeSOD and MnSOD) being among the most widely distributed forms in bacteria. Other SOD families, including CuZnSOD and NiSOD, are also present across diverse bacterial lineages [5,6]. Despite their distinct metal centers, SODs share highly conserved catalytic architectures that enable the efficient dismutation of superoxide radicals. Beyond their metal cofactors, SODs also exhibit striking differences in their subcellular localization. In higher plants, FeSOD and CuZnSOD are predominantly localized in chloroplasts, whereas MnSOD is primarily targeted to mitochondria [7,8]. MnSOD is thought to have been lost from plastids during plastid evolution [9,10]. In contrast, the green alga *Chlamydomonas reinhardtii* retains MnSOD in both the chloroplast lumen and mitochondria [11], illustrating the diversity of SOD compartmentalization among photosynthetic eukaryotes. Cyanobacteria display an even broader range of SOD localization, including membrane-associated and lumenal MnSODs associated with the thylakoid system. Consistent with this diversity, genomic analyses reveal substantial variation in SOD complements among cyanobacteria, with some species encoding multiple SOD isoforms and others possessing only a single SOD [6,12]. In the well-studied cyanobacterial models *Synechocystis* sp. PCC 6803 and *Synechococcus elongatus* PCC 7942, *sodB* encodes a cytosolic FeSOD and represents the sole SOD gene identified in either organism [13,14]. Its expression responds to oxidative and light stress, consistent with an important role in protecting cells against O_2_^.-^ generated during photosynthesis. By contrast, *Anabaena* sp. PCC 7120, *Pseudanbaena* PCC 7367, and several other cyanobacteria possess both *sodB*, encoding a cytosolic FeSOD, and *sodA*, encoding a thylakoid-localized MnSOD [15,16]. The presence of a single *sodB* gene in *Synechocystis* sp. PCC 6803 and *Synechococcus elongatus* PCC 7942 thus reflects a relatively streamlined SOD system, in which FeSOD alone serves as a sufficient and efficient enzymatic defense against O_2_^.-^ radicals. Since its identification as a thylakoid-bound enzyme in *Anabaena* sp. PCC 7120, MnSOD has been implicated in heterocyst function and proposed to protect photosystem II (PSII), yet its broader physiological role remains unclear. Additional SOD types are sporadically distributed among cyanobacteria, including CuZnSod in a few species and NiSod primarily in marine lineages. This diversity raises fundamental questions about the functional importance and specialisation of SOD multiplicity in cyanobacteria. The unicellular, nitrogen-fixing cyanobacterium *Cyanothece* sp. PCC 7425 provides an informative system to address this question, as its genome encodes both a conserved soluble FeSOD and a putative membrane-associated MnSOD. Using comparative genetics, and biochemical approaches, we show that soluble FeSOD makes the dominant contribution to O_2_^.-^ defense, underscoring the importance of cytoplasmic O_2_^.-^ detoxification, consistent with high O_2_^.-^ production at the PSI acceptor side. These findings challenge the assumption that membrane association necessarily confers a major role in photosynthetic superoxide defense, and provide a framework for understanding the differential retention and loss of SOD isoforms across cyanobacteria and plants.

## Materials and methods

### Strains and culture conditions

All strains of *Cyanothece* PCC 7425 were cultured at 30°C with continuous agitation at 140 rpm (Infors rotary shaker). Cultures were maintained under cool-white light (Sylvania Luxline Plus) in standard conditions at an intensity of 2500 lux (32 μmol photons m^-2^s^-1^), low-light intensity of 600 lux (8 μmol photons m^-2^s^-1^), or high-light intensity of 8000 lux (100 μmol photons m^-2^s^-1^). Cells were grown in Mineral Medium (MM) containing 1.89 mM sodium carbonate (Na_2_CO_3_) as the carbon source. For growth on solid medium, MM was supplemented with 1% Bacto Agar (Difco). *Escherichia (E.) coli* strains NEB 10-beta (New England Biolabs) and AM5501, used for plasmid cloning and conjugative transfer, were cultured in liquid or solid LB medium supplemented with antibiotics when required. The antibiotics were used at the following concentrations : ampicillin (Amp), 100 μg mL^-1^; kanamycin (Km), 50 μg mL^-1^; streptomycin (Sm), 50 μg mL^-1^; and spectinomycin (Sp), 50 μg mL^-1^. For *Cyanothece* PCC 7425, antibiotics were used at lower concentrations : 5 μg mL^-1^ for Sm, 5 μg mL^-1^ for Sp, and 25 μg mL^-1^ for Km.

### Plasmid constructions

All plasmids used in this study are listed in **Table S1**. Cloning was performed in the *E. coli* strain NEB10-beta. PCR fragments were cloned into the NdeI/ EcoRI sites of pFCK using the NEB Gibson Assembly Cloning Kit. The primers used for cloning are listed in **Table S2**. All enzymes were purchased from NEB, except AarI, which was obtained from Thermo Fisher Scientific. All plasmids were verified by sequencing at Eurofins Genomics.

#### pSL2680-ΔsodA plasmid construction

For the construction of vector pSL-ΔsodA, the complementary phosphorylated oligos cr-sodA_Fw/Rv (Integrated DNA Technologies) corresponding to the designed crRNA were annealed by heating at 65 °C and ligated at the AarI-linearized pSL2680 vector, generating vector pSL-crRNA-sodA. After successful cloning of the crRNA, the homology platforms of ∼600 bp lying both upstream the start codon of *sodA* and downstream the stop codon of *sodA* gene were generated by overlapping PCR (Phusion polymerase, NEB) on genomic DNA of *Cyanothece* 7425 using oligos F1-sodA_Fw/Rv and F3-sodA_Fw/Rv, respectively. The Sm^R^Sp^R^ cassette was obtained by PCR using oligos F2-Ω_Fw/Rv (**Table S2**) on purified pC plasmid. The three fragments were cloned at the SalI site in pSL-crRNA-sodA using NEB Gibson assembly cloning kit, generating the final vector pSL-ΔsodA. The vector pSL-ΔsodB was previously described in [17].

#### pFCK derivative plasmids construction

All pFCK deriative plasmids (**Table S1**) were generated by cloning PCR amplified fragments at the NdeI/ EcoRI using the NEB Gibson assembly cloning kit.

### Transfer of plasmids via conjugation

The plasmids were transferred from *E. coli* to *Cyanothece* PCC 7425 and the other cyanobacteria by either triparental or biparental conjugation, using *E. coli* NEB 10-beta together with the helper strain CM404 for triparental conjugation, or strain AM5501 for biparental conjugation, as previously described in [17].

### Selection of ΔsodA mutant

The pSL-ΔsodA was transferred to *Cyanothece* 7425 via triparental conjugation as described. Colonies were selected on MM plates containing 5 μg mL^-1^ SpSm. Their genomes were analysed by PCR to confirm deletion and full segregation of the chromosomes.

### Spot viability assay

For spot viability assay, the wild-type and SOD mutant strains were grown in liquid MM until exponential phase (OD_750nm_ ∼0.8 - 1.2) at standard-light intensity. A drop of 10 μL from each dilution was spotted on MM agar plates and incubated for 8 days at 30°C with the indicated light intensities.

### SOD in-gel activity assay on non-denaturing gel

Cultures of WT and SOD deletion strains were grown to exponential phase (OD_750nm_ : 0.8 - 1.2). Protein extracts were prepared in PBS (pH 7.6) as previously described (Ivleva and Golden, 2007), and protein concentrations were determined using the Pierce BCA Protein Assay Kit (Thermo Scientific). To separate the membrane and soluble fractions, total protein extracts were centrifuged at 100.000 g for 60 min at 4°C. The membrane fraction was washed and resuspended in PBS, and proteins were solubilized using 1% (W/V) DDM (dodecyl maltoside) when required. For SOD activity analysis, 40 μg of protein extracts were resolved by non-denaturing polyacrylamide gel electrophoresis and stained for SOD activity as previously described (Weydert and Cullen, 2010). Briefly, gels were incubated for 15 min at room temperature (RT) in 20 mL of 50 mM Tris-HCl (pH 7.2) containing 0.85% TEMED (Sigma-Aldrich) and 56 μM riboflavin-5-phosphate (Sigma-Aldrich). Gels were subsequently incubated in 2 mg mL^-1^ nitroblue tetrazolium chloride (Sigma-Aldrich) for 15 min in the dark at RT. After two washes with water, gels were placed in a light box at RT until SOD activity was visualized.

### Western blotting and immunodetection of FeSOD

Equal amounts of total protein (20 μg) from WT and SOD mutant strains were separated by 14% SDS-PAGE and transferred onto Hybond ECL polyvinylidene difluoride (PVDF) membranes (GE Healthcare). Membranes were blocked in TBST buffer (50 mM Tris, 150 mM NaCl, and 0.1% Tween 20 (v/v), pH 7.4) supplemented with 5% BSA (w/v). Following three 5-min washes with TBST, membranes were incubated for 1 h with α-FeSOD antibodies (PhytoAB) diluted 1:1000 in TBST. After washing with TBST, membranes were incubated for 1 h at room temperature with an HRP-conjugated goat anti-rabbit secondary antibody (Bio-Rad) diluted 1:5000 in TBST. Reactive bands were detected using a chemiluminescent HRP substrate, and images were acquired using a ChemiDoc camera system (Bio-Rad).

### Disc diffusion assay

500 μL aliquot of WT or SOD deletion strain cultures (OD_750nm_ : 0.8) were mixed with 20 mL of 50% semi-solid MM agar and uniformly spread onto 20 mL MM agar plates. Sterile 6 mm discs were saturated with 5 µL of 10 mM menadione (Sigma-Aldrich) and placed in the center of each plate. Plates were incubated at 30 °C under the specified light conditions for 7 days prior to photography.

### Phylogenetic tree construction

Homologues of the query sequence (*Cyanothece* PCC 7425 MnSOD; WP_041237046.1) were identified by BLASTp search against the NCBI non-redundant protein database, restricted to cyanobacterial sequences. BLAST hits were filtered using an e-value cutoff of ≤1×10^-5^, minimum sequence identity of 30%, and minimum query coverage of 50%, yielding 164 non-redundant sequences for further analysis (**Table S4**). Metagenome-assembled genomes were also excluded. *E. coli* MnSOD was included as an outgroup. Signal peptide and lipoprotein signal prediction was performed using SignalP 6.0 (https://services.healthtech.dtu.dk/services/SignalP-6.0/) and SMART (Simple Modular Architecture Research Tool: http://SMART.embl-heidelberg.de) (**Table S4**). Sequence alignment was performed using the https://www.phylogeny.fr/ platform [18]. Sequences were classified into four topology classes based on SignalP 6.0 predictions and multiple sequence alignment: lipoprotein, soluble, membrane-associated, and other. Multiple sequence alignment was performed using MAFFT v7 with the auto strategy [19]. Poorly aligned and gap-rich columns were trimmed using trimAl v1.4 in automated1 mode [20]. A maximum likelihood phylogenetic tree was inferred using IQ-TREE2 with automatic substitution model selection and 1000 ultrafast bootstrap replicates [21]. The tree was visualized and annotated using the Interactive Tree of Life (iTOL) v6 [22]. All sequence processing, alignment, and tree inference steps were performed using the Galaxy platform (usegalaxy.eu).

## Results

### *Cyanothece* PCC 7425 has two SOD encoding genes

Despite the production of toxic ROS, including O_2_^.-^ during photosynthesis (**Fig. 1**), *Cyanothece* PCC 7425, like other cyanobacteria, thrives even under high-light stress. Along with singlet oxygen, O_2_^.-^ is among the earliest ROS generated by the photosystems, placing the SODs at the first line of defense by catalyzing its conversion to H₂O₂ and preventing photoinhibition. Genome analysis of *Cyanothece* PCC 7425 identified two genes, *cyan7425_3014* and *cyan7425_1660*, encoding putative MnSOD and FeSOD, respectively, both of which may contribute to O_2_^.-^ detoxification. Sequence comparisons revealed strong conservation between the two proteins, except at the N-terminus of MnSOD, which contains an additional signal peptide and a lipoprotein-binding motif (lipobox) (**Fig. 2A** and **Fig. S1**). This feature suggests that MnSOD is a membrane-anchored lipoprotein, potentially localized to the cytoplasmic and/or thylakoid membrane, where it may protect the periplasmic space and/or thylakoid lumen from superoxide. Such membrane anchoring of SODs trough a N-terminal lipid modification has been reported for the CuZnSOD of *Mycobacterium tuberculosis* [23] and suggested for the *Anabaena* sp. PCC 7120 MnSOD [16]. This unusual feature prompted us to examine cyanobacterial genomes more broadly to investigate the distribution and diversity of MnSODs in this phylum.

**Figure 1.**
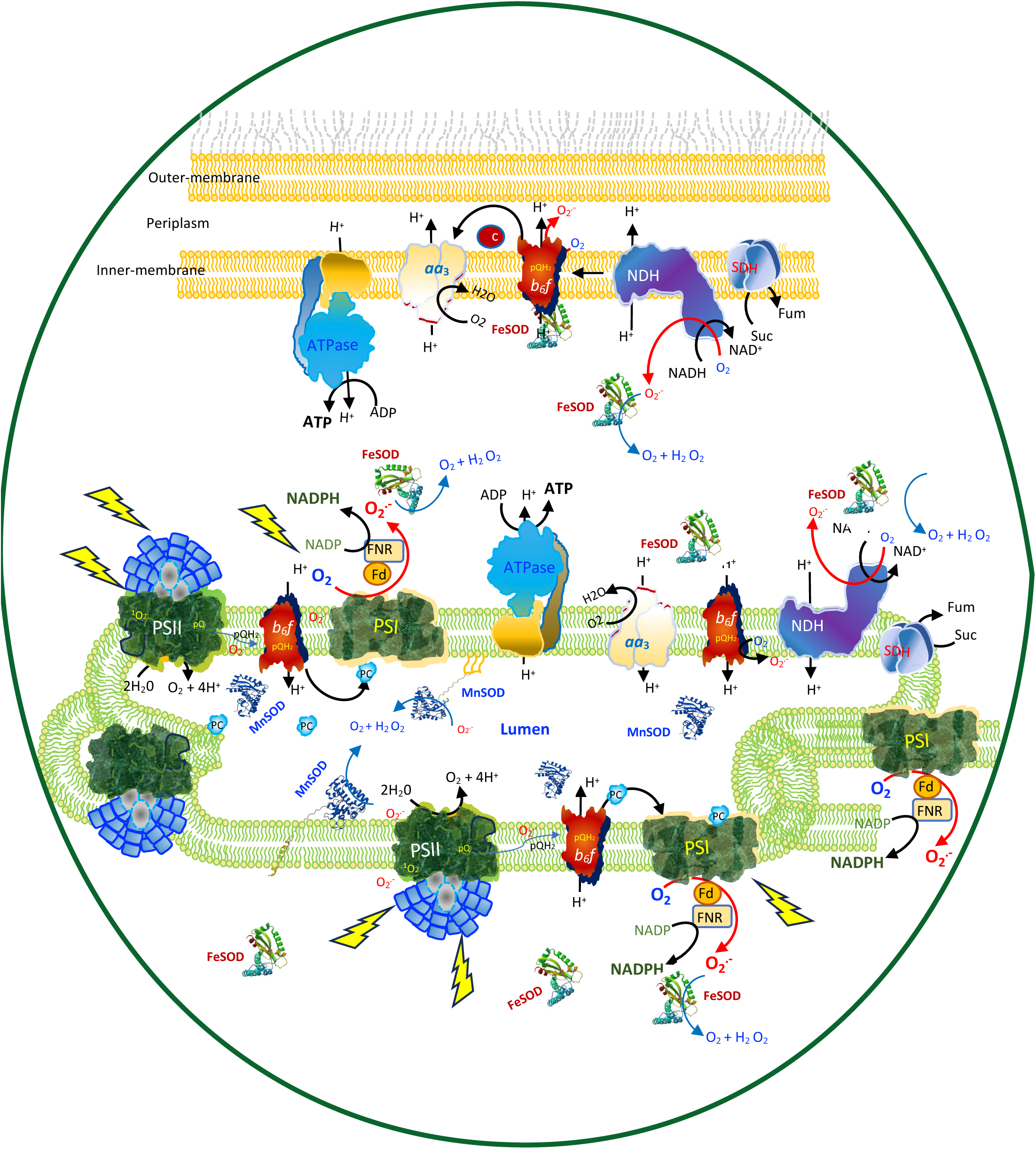
Subcellular sites of superoxide generation and SOD localisation in cyanobacteria. Schematic of the cyanobacterial cytoplasmic and thylakoid membranes, showing sites of O_2_^.-^ generation and the predicted subcellular localisation of SODs. The respiratory electron transfer chain occupies both membranes, whereas the photosynthetic chain is restricted to the thylakoid membrane. Increased light exposure promotes electron leakage from both chains, generating O_2_^.-^, which SODs dismutate into O_2_ and H_2_O_2_. FeSOD is cytoplasmic, whereas MnSOD is predicted to localise to the thylakoid membrane or lumen, as well as the cytoplasmic membrane and periplasm.

**Figure 2.**
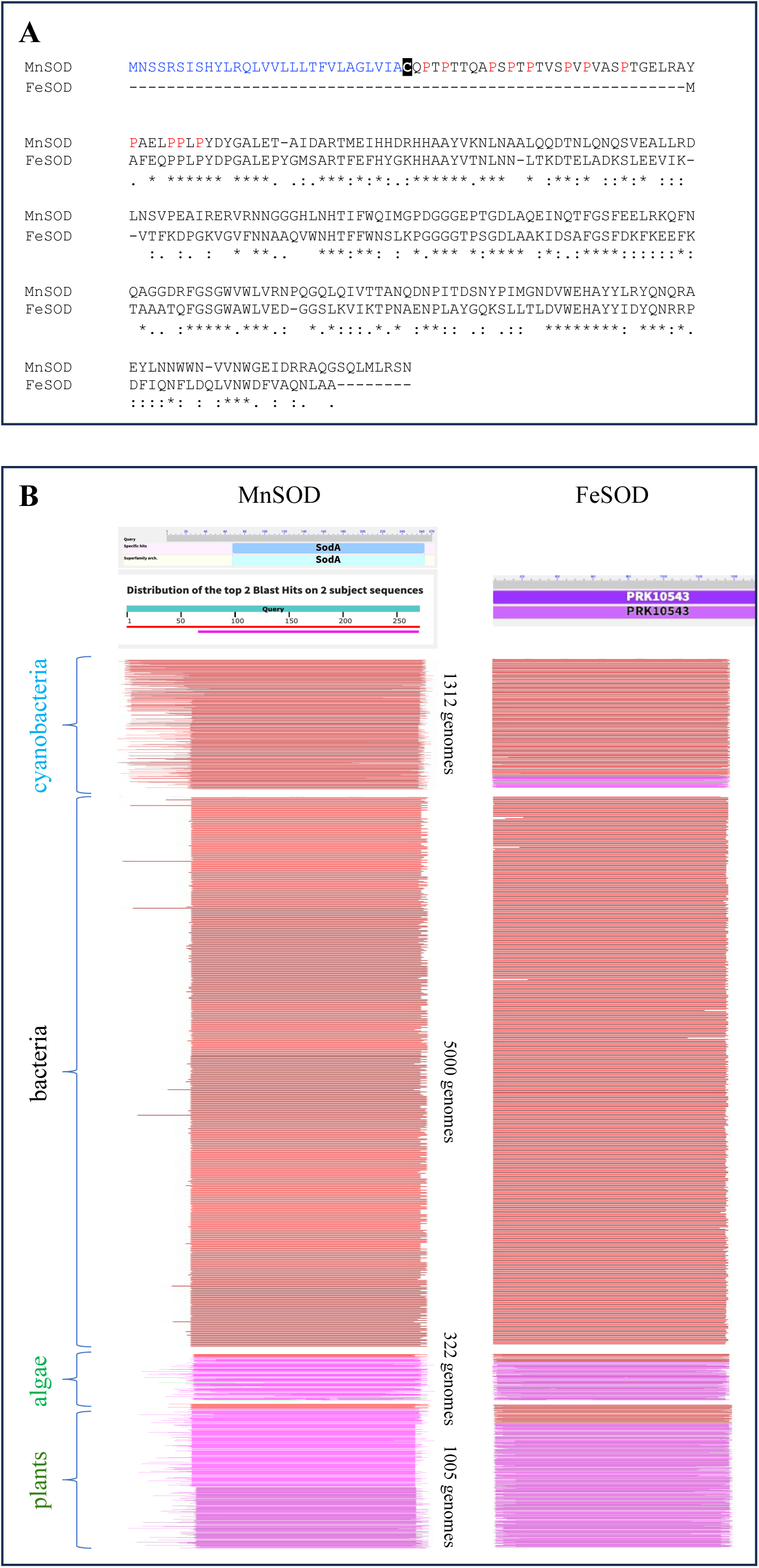
(**A**) **Sequence alignment of MnSOD and FeSOD from *Cyanothece* sp. PCC 7425**. MnSOD’s N-terminus carries a lipoprotein signal peptide (blue) and targeting motif, with a predicted cleavage site immediately preceding the conserved cysteine that undergoes lipid modification for membrane anchoring; proline-rich residues within the putative “hinge” domain are shown in red. FeSOD, a soluble protein, lacks both features. asterisks indicate residues conserved between the two. (**B**) **BLASTP analysis of MnSOD and FeSOD against cyanobacterial, non-cyanobacterial bacterial, algal, and plant genomes**. The MnSOD-type N-terminal extension was widespread among cyanobacteria but found in only four bacterial genomes (excluding cyanobacteria), and was generally absent from algal and plant MnSODs. FeSODs, by contrast, were highly conserved across all groups, consistent with a soluble localisation. Full BLASTP outputs are provided in the supplementary data.

### A conserved N-terminal protein extension distinguishes cyanobacterial MnSODs

As aerobic respiration and oxygenic photosynthesis, the main cellular sources of O_2_^.-^, are localized in the thylakoid membrane, cyanobacteria are expected to harbor thylakoid-associated SODs, either lumenal or membrane-bound, that would constitute the primary line of defense against O_2_^.-^. Although the phylogeny and evolutionary history of cyanobacterial SODs have been extensively examined [6,12], gaps remain in our understanding of their diversification, particularly with respect to thylakoid-localized SODs. Clarifying this diversification is essential for elucidating SOD function and evolution in the context of the emergence of oxygenic photosynthesis. To begin addressing this question, the MnSOD from *Cyanothece* PCC 7425 was used as a query in BLASTP searches against 1312 cyanobacterial, 5000 bacterial (excluding cyanobacteria), 322 algal, and 1005 plant genomes. As expected, MnSOD sequences were detected in all analyzed genomes. Strikingly, the majority of cyanobacterial MnSODs contained an N-terminal extension similar to that of *Cyanothece* PCC 7425 MnSOD (**Fig. 2B and Fig. S2**), whereas this feature was detected in only 4 of the 5000 bacterial (excluding cyanobacteria) genomes analyzed (**Fig. 2B**). In contrast, MnSODs identified in algal and plant genomes generally lacked this N-terminal extension and are therefore likely soluble in the chloroplast and mitochondria, as previously proposed. A parallel BLASTP analysis of FeSODs revealed a high level of conservation across all examined organisms, consistent with a soluble cytosolic localization (**Fig. 2B**). Together, these observations strongly suggest that the N-terminal extension represents a “cyanobacterial specific” evolutionary adaptation for targeting MnSOD to the thylakoid membranes.

### N-terminal targeting signals define distinct cyanobacterial MnSOD classes

As in bacteria, protein translocation in cyanobacteria across the plasma and thylakoid membranes occurs *via* the Sec (secretory) and Tat (twin-arginine translocation) pathways, whereas integral membrane proteins are inserted into membranes by the signal recognition particle (SRP) system [24]. Cyanobacteria also harbor lipoproteins along with the lipoprotein-processing enzymes Lgt, LspA, and Lnt [25]. These lipoproteins are synthesized as precursors containing N-terminal signal peptides followed by a lipid-modified cysteine residue. Consequently, analysis of the N-terminal sequences of cyanobacterial MnSODs may provide insights into their subcellular localization and allow further subclassification of these MnSODs. Therefore, 164 cyanobacterial MnSOD sequences were aligned using Clustal, and analyzed with both the SignalP 6.0 [26] and SMART [27] programs to search for possible signal peptides and translocation-associated domains (**Fig. S3 - S4**). In these analyses we included the ancestral *Pseudanabaena* sp. PCC 7367 and the thylakoids-less cyanobacterium *Gloeobacter (G.) violaceus* PCC 7421 given their early branching phylogenetic position; as well as the soluble MnSOD sequence from the evolutionary-distant bacterial model *E. coli*. SignalP 6.0 and Smart analyses discriminated three distinct classes of MnSODs (**Figure 3, Fig. S3 - S4**): (i) MnSODs harboring either a Tat signal peptide as observed in both *Synechococcus* sp. PCC 6312 and *Pseudanabaena* sp. PCC 7367, or Sec signal peptide as found in *G. violaceus* PCC 7421, suggesting that they are translocated across the cytoplasmic and/or thylakoid membranes, and subsequently targeted to the periplasm or the lumen. (ii) MnSODs containing a single transmembrane α-helix, suggesting that they are located in membranes. This simple architecture is observed in MnSODs from *Nostoc* sp. and MnSOD2 from *Nostoc (N.) punctiforme* PCC 73102 whose thylakoid membrane localization was previously inferred based on its N-terminal sequence [28,29]. (iii) MnSODs bearing either a lipoprotein-signal peptide as in *Cyanothece* PCC 7425 or a Tat-lipoprotein signal peptide as in MnSOD1 from both *N. punctiforme* PCC 73102 and *Anabaena* sp. PCC 7120, characterized by a conserved cysteine at the +1 position of the cleavage site, which undergoes lipid modification, thereby anchoring the protein to the membrane (**Fig. 3A** and **Fig. S1 - S4**). In addition, both putative membrane-associated MnSODs possess a « proline-rich hinge domain » which may function as a flexible spacer between the membrane-anchor and the SOD catalytic domains (**Fig. 3A** and **Fig. S1 - S4**), allowing the mobility and activity of MnSODs in the vicinity of the membrane and within the periplasmic or lumenal space. To examine the evolutionary relationships among these proteins, a maximum likelihood phylogenetic tree of the selected MnSODs and FeSODs sequences was constructed, using *E. coli* MnSOD and FeSOD sequences as outgroups for comparison (**Fig. 3B**). The phylogenetic reconstruction separated the two well supported major clades corresponding to MnSODs and FeSODs, confirming that these proteins have undergone substantial evolutionary divergence. Within the MnSOD clade, the *E. coli* MnSOD sequence emerged as a distant branch, indicating an early evolutionary divergence from the cyanobacterial MnSOD proteins. Among cyanobacteria, MnSOD from *Pseudanabaena* sp. PCC 7367, and *Synechococcus* PCC 6312, both soluble, branched off first indicating earliest evolution from the remaining strains. MnSOD of *G. violaceus* PCC 7421 clusters with a subgroup comprising MnSOD from the nitrogen-fixing *Anabaena* sp. PCC 7120, *N. punctiforme* PCC 73102, and *Cyanothece* PCC 7425, reflecting a closer evolutionary relationship among these predicted membrane localized MnSODs (**Fig. 3B**). The closest association, supported by a high bootstrap value is observed between MnSOD from *Cyanothece* PCC 7425 and *N. punctiforme* PCC 73102, which also clusters also with *Anabaena* sp. PCC 7120, and all three MnSODs were predicted to be lipoproteins, indicating they have a common ancestor. The FeSOD clade exhibited a notable difference in branching order. The “ancestral” branching cyanobacteria *Pseudanbaena* PCC 7367 and *G. violaceus* PCC 7421 diverge early from *E. coli* and the remaining cyanobacterial sequences. Furthermore, a maximum-likelihood phylogenetic tree was constructed from the 164 cyanobacterial MnSODs sequences (**Table S4**) to investigate the evolutionary distribution of predicted protein cellular localization **(Fig. 4A)**. Sequences were classified into four predicted cellular localization classes: lipoprotein (n=114), and membrane-associated (n=14), soluble (n=34), and other (undefined) (n=2). The lipoprotein form was the predominant topology, accounting for nearly 70% of the analyzed sequences, and was broadly distributed across the tree, encompassing representatives from diverse cyanobacterial orders including Nostocales, Oscillatoriales and Pleurocapsales. The MnSOD from *Cyanothece* PCC 7425 was also classified as a lipoprotein and clustered within a predominantly lipoprotein-containing clade, consistent with its predicted lipoprotein signal peptide. The widespread occurrence of the lipoprotein form across phylogenetically diverse cyanobacterial lineages indicates that this topology is deeply conserved across the cyanobacterial phylum. Interestingly, the soluble form was restricted to a well-supported, phylogenetically distinct clade (bootstrap ≥ 95), comprising predominantly thermophilic *Thermosynechococcus* strains. The number of soluble MnSODs within this clade, particularly among thermophilic lineages, raises the possibility that the soluble localization might be associated with adaptation to high-temperature environments or arose through lineage-specific loss of the lipoprotein signal peptide. In contrast, the non-lipoprotein membrane-associated form was exclusively restricted to *Nostoc* species, forming a well-supported monophyletic clade (bootstrap ≥ 95) within the broader Nostocales. All 14 membrane-associated MnSODs sequences were derived from *Nostoc punctiforme* PCC 73102 or uncharacterised *Nostoc* strains, indicating a lineage-specific transition to membrane association within this genus. The occurrence of membrane-associated forms (both lipoproteins and non-lipoproteins) within *Nostoc* further suggests that diversification of MnSODs cellular localisation has occurred within this lineage.

**Figure 3.**
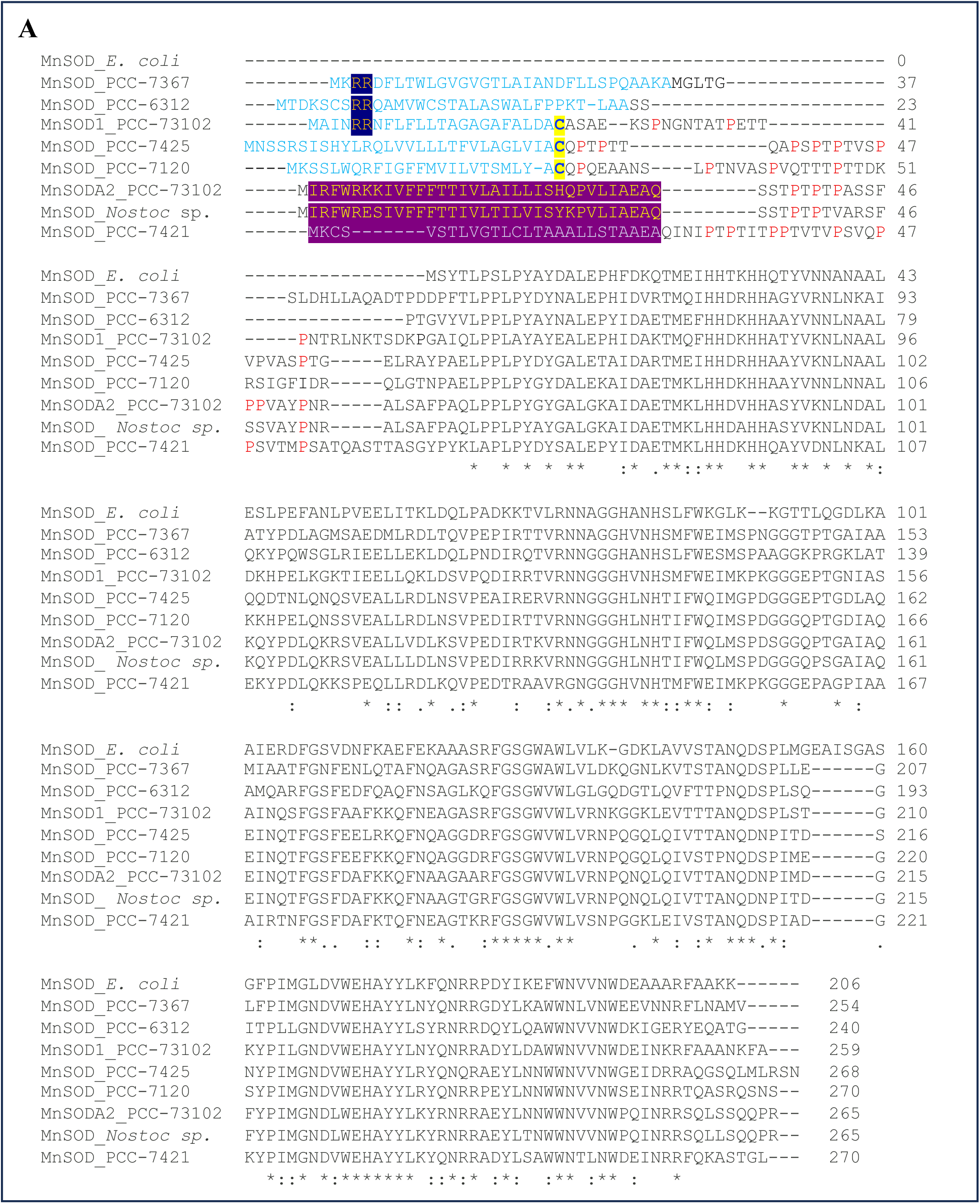
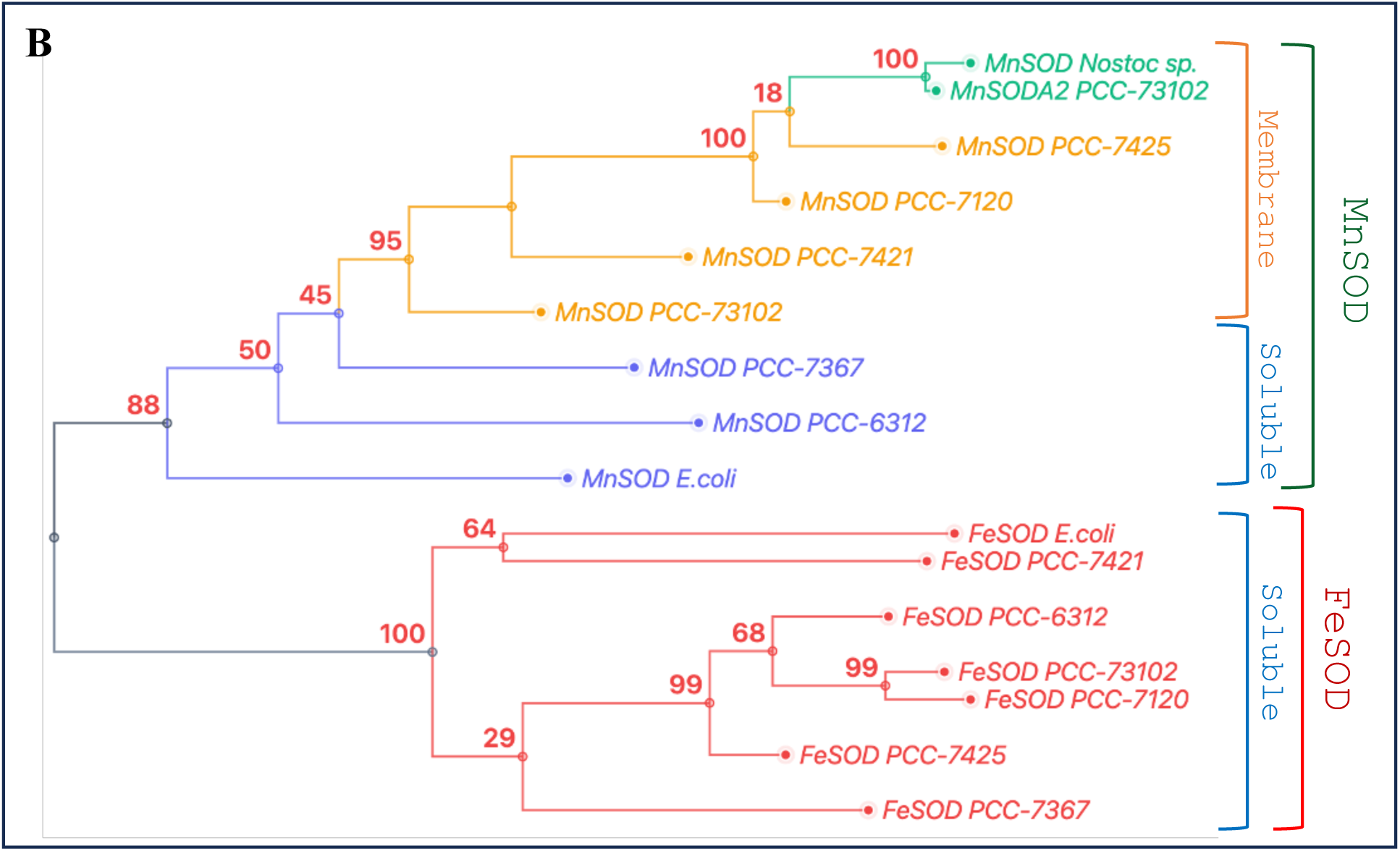
**(A) Sequence alignment of MnSODs from selected cyanobacteria, grouped by predicted N-terminal targeting signal**: (i) Tat or Sec signal peptides (cyano) in *Synechococcus* sp. PCC 6312, *Pseudanabaena* sp. PCC 7367, suggesting periplasmic or luminal translocation; (ii) a putative single transmembrane α-helix in *Nostoc* punctiforme PCC 73102 and *Nostoc* sp., suggesting membrane localization; (iii) a Sec-lipoprotein signal peptide in *Cyanothece* sp. PCC 7425, indicating membrane anchoring; and (iv) a Tat-lipoprotein signal peptide, as in MnSOD1 from both *N. punctiforme* PCC 73102 and *Anabaena* sp. PCC 7120. Prolines in the Pro-rich domain are shown in red. The N-terminus of *Gloeobacter* PCC-7421 MnSOD could either be processed or remain uncleaved, resulting in a membrane protein. Soluble *E. coli* MnSOD is included for comparison; asterisks mark conserved residues. (**B**) **Phylogenetic dendrogram of MnSOD and FeSOD sequences**, resolving two well-supported clades corresponding to the two SOD types; within the MnSOD clade, membrane-associated and soluble (periplasmic/luminal) variants segregate clearly.

**Figure 4.**
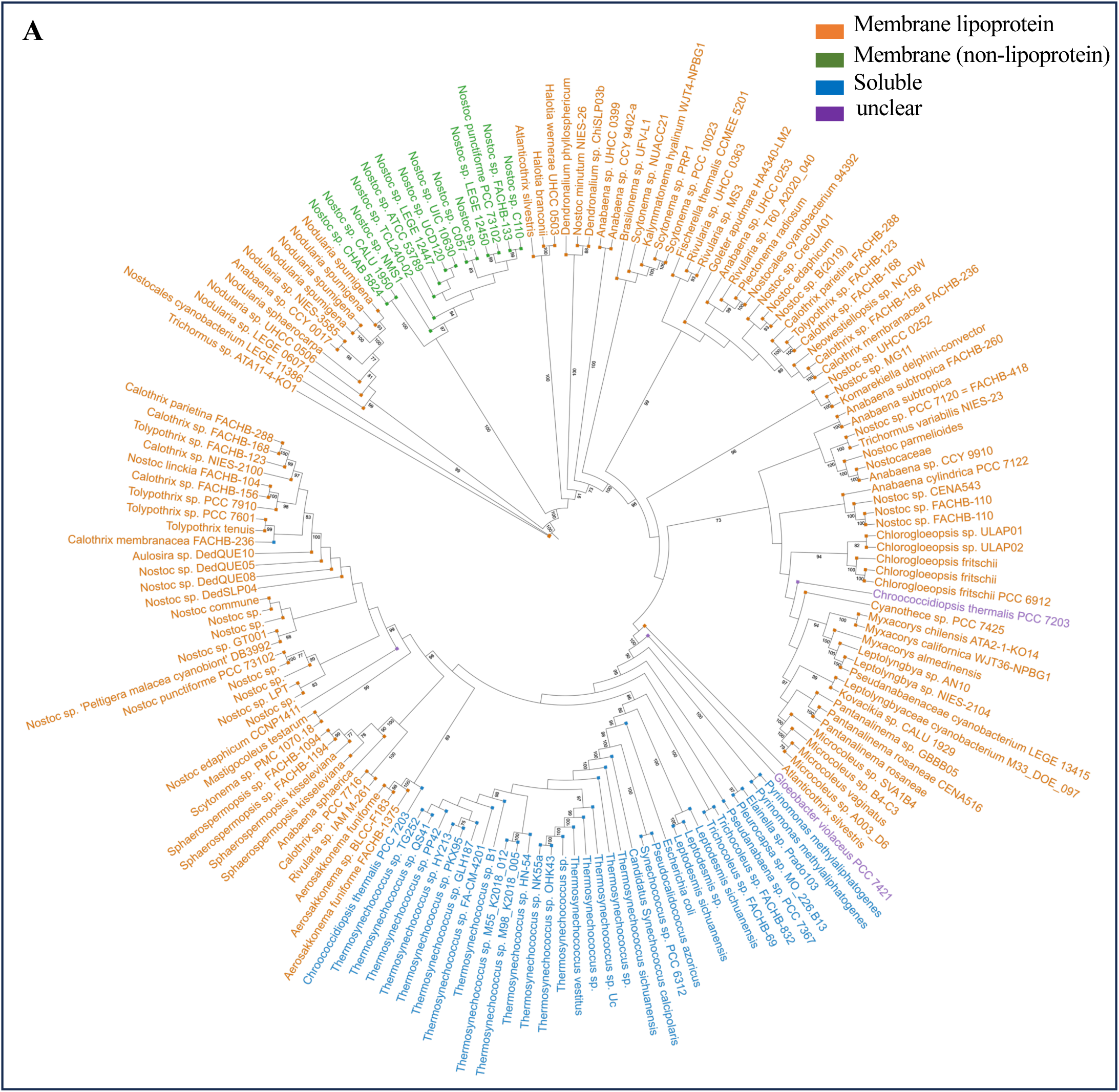
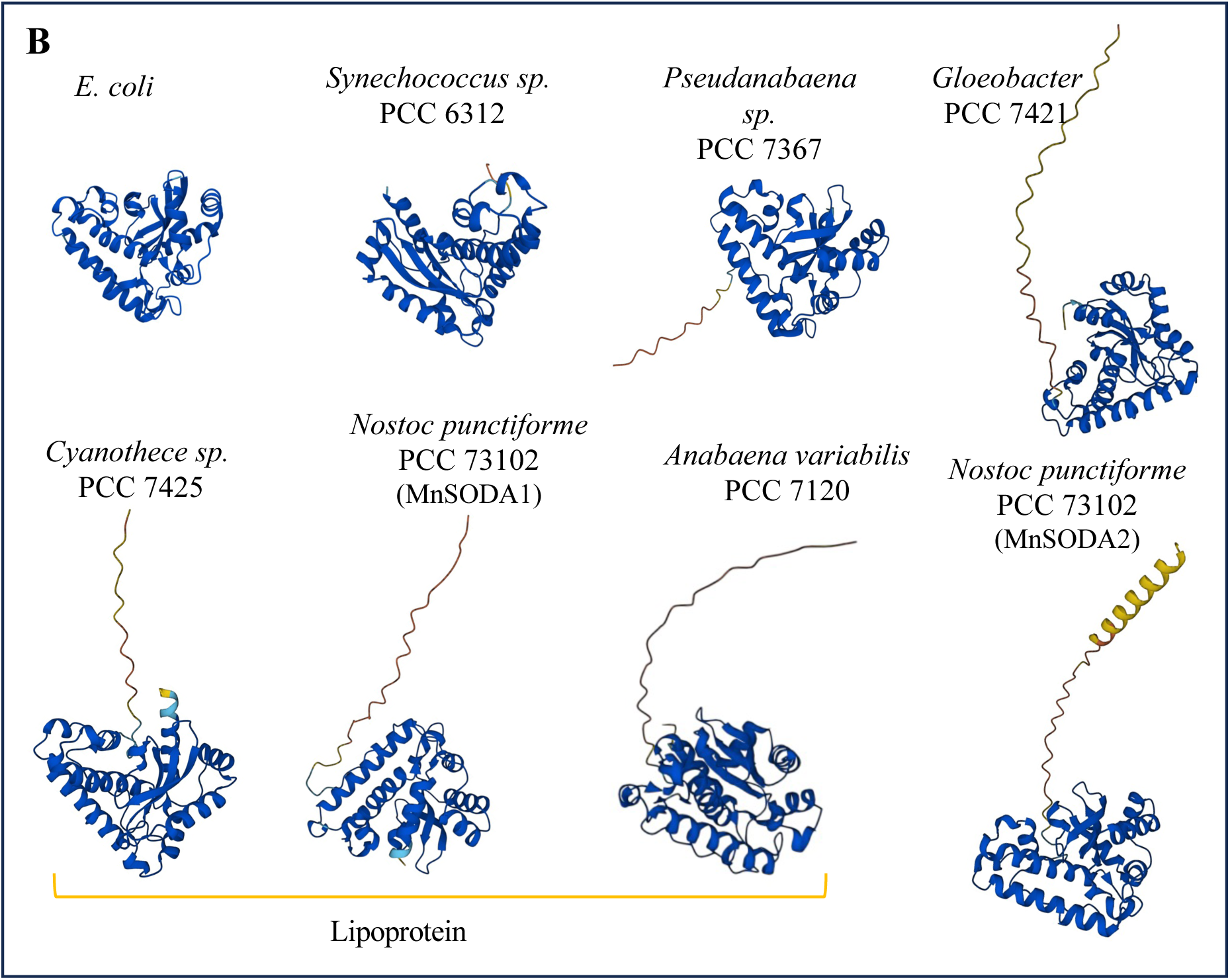
(**A**) Phylogenetic analysis of cyanobacterial MnSODs according to predicted protein topology. Maximum-likelihood phylogenetic tree constructed from 164 conserved cyanobacterial MnSOD sequences. Tip colours indicate the predicted protein topology: lipoprotein (orange), membrane-associated (green), soluble (blue), and unclear (membrane or soluble) (purple). Bootstrap support values are indicated at the corresponding nodes. (**B**) AlphaFold2-predicted structures of SODs from selected cyanobacteria and *E. coli*; MnSOD models were generated from the mature sequence, following removal of the signal peptide at its cleavage site.

### Structural predictions reveal distinct N-terminal architectures of cyanobacterial MnSODs

To investigate whether structural features of the MnSOD N-terminal sequence are associated with cellular localization, we compared AlphaFold-predicted structures of the selected cyanobacterial MnSODs with that of the soluble MnSOD from *E. coli* (**Fig. 4B**). As expected, MnSOD proteins from *Synechococcus* PCC 6312 and *Pseudanabaena* PCC 7367, both predicted to be soluble, exhibited structures highly similar to *E. coli* MnSOD. Although the *Pseudanabaena* PCC 7367 protein contains an additional 14 amino-acids unstructured N-terminal extension, it is still predicted to be targeted to the periplasm or thylakoid lumen, as previously suggested by Tamanna *et al* [15]. In contrast, MnSODs from *Cyanothece* PCC 7425, *N. punctiforme* PCC 73102 and *Anabaena* sp. PCC 7120 possess intrinsically disordered N-terminal extensions of 30, 24 and 25 amino acids, respectively, containing the conserved cysteine residue required for lipidation and subsequent proline rich domain. Similarly, the MnSOD from *G. violaceus* PCC 7421 exhibits a 39-amino-acid unstructured N-terminal extension. In contrast to the MnSOD of *Anabaena* sp. PCC 7120, isolated from thylakoid membranes, the extension in *G. violaceus* MnSOD lacks the lipidated cysteine, making membrane attachment through lipid modification unlikely. Consequently, mature *G. violaceus* MnSOD may either remain soluble in the periplasm or associate with the membrane through an alternative mechanism involving its unstructured N-terminal extension. Although no experimental evidence is currently available to distinguish between these possibilities, the presence of a putative proline-rich hinge domain is consistent with a membrane-associated form of MnSOD also in *G. violaceus* PCC 7421.

### FeSOD predominates over MnSOD in cyanobacterial SOD activity

Expression of the *sodA* and *sodB* genes in *Cyanothece* PCC. 7425 was first assessed by semi-quantitative RT-PCR, which showed that both genes are transcribed under all tested light conditions (**Fig. S5**). Proteomic analysis further identified the presence of FeSOD only in the soluble protein fraction, with 16 tryptic peptides identified by MS/MS. In contrast, only three MnSOD derived peptides were detected in the membrane fraction, indicating that MnSOD is expressed, but at a comparatively low abundance (**Fig. S6**). MnSOD was also detected in a previous proteomic study of the genus Cyanothece by Aryal *et al.* [30]. To further analyze the abundance of MnSOD and FeSOD, their superoxide dismutase activity was assessed using an *in-gel* NBT-SOD assay. *Synechocystis* PCC 6803 which contains only FeSOD, and *Synechococcus* PCC 6312, predicted to contain both soluble MnSOD and FeSOD, were used as reference strains. As expected, a single SOD active band was detected in total protein extracts from *Synechocystis* PCC 6803, whereas two distinct SOD active bands were observed in *Synechococcus* PCC 6312, consistent with the presence of both FeSOD and MnSOD (**Figure 5A**). Notably, only a single SOD active band, corresponding to FeSOD, was detected in total protein extracts from *Cyanothece* PCC 7425. Because *Cyanothece* PCC 7425 MnSOD is predicted to be membrane-associated, soluble and detergent-solubilized membrane fractions from *Cyanothece* PCC 7425 and *Synechococcus* PCC 6312 and were subsequently analyzed separately (**Fig. 5B - 5C**). In the soluble fraction of *Synechococcus* PCC 6312, two distinct SOD active bands were detected in addition to the soluble phycobiliproteins (PBS) (**Fig. 5B**). Based on their migration pattern, these SOD bands were assigned to FeSOD and MnSOD, respectively, confirming the presence of soluble MnSODs suggested by our above-mentioned (sequence-based) predictions, and supporting their presence in either the periplasm or thylakoid lumen. Remarkably, the SOD activity associated with FeSOD was markedly stronger than that associated with MnSOD. In *Cyanothece* PCC 7425, FeSOD was readily detected in the soluble fraction, together with a faint SOD active band indicated by an asterix, which may correspond to a truncated soluble form of MnSOD (**Fig. 5B**). However, as observed in *Synechococcus* PCC 6312, the SOD activity associated with FeSOD was markedly higher than that of the putative MnSOD. Detergent solubilized membrane fractions of *Synechococcus* PCC 6312 and *Cyanothece* PCC 7425 contained the expected solubilized photosystems and carotenoid-containing membrane proteins observed on the native electrophoresis gel (**Fig. 5C**). However, the NBT-SOD assay detected only the soluble FeSOD were detected in both strains (**Fig. 5C**), with no distinct SOD active band attributable to MnSOD in the detergent solubilized membrane fractions from repeated samples of *Cyanothece* PCC 7425. This absence of detectable membrane-associated MnSOD activity may be attributable to its low abundance and/or its loss or inactivation during membrane solubilization. Nevertheless, the detection of *sodA* transcripts, the detection of MnSOD-derived peptides in the membrane fraction, and the faint putative MnSOD-associated activity in the soluble fraction, collectively support MnSOD expression in *Cyanothece* PCC 7425. Taken together, these results demonstrate (i) the expression of both soluble MnSOD and FeSOD in *Synechococcus* PCC 6312 (ii) the expression of FeSOD and, most likely, MnSOD in *Cyanothece* PCC 7425, although no specific MnSOD activity was detected in the membrane fraction; and (iii) substantially higher SOD activity associated with FeSOD than with MnSOD. These findings suggest that FeSOD represents the predominant SOD contributing to superoxide detoxification under the conditions tested in *Synechococcus* PCC 6312 and *Cyanothece* PCC 7425.

**Figure 5.**
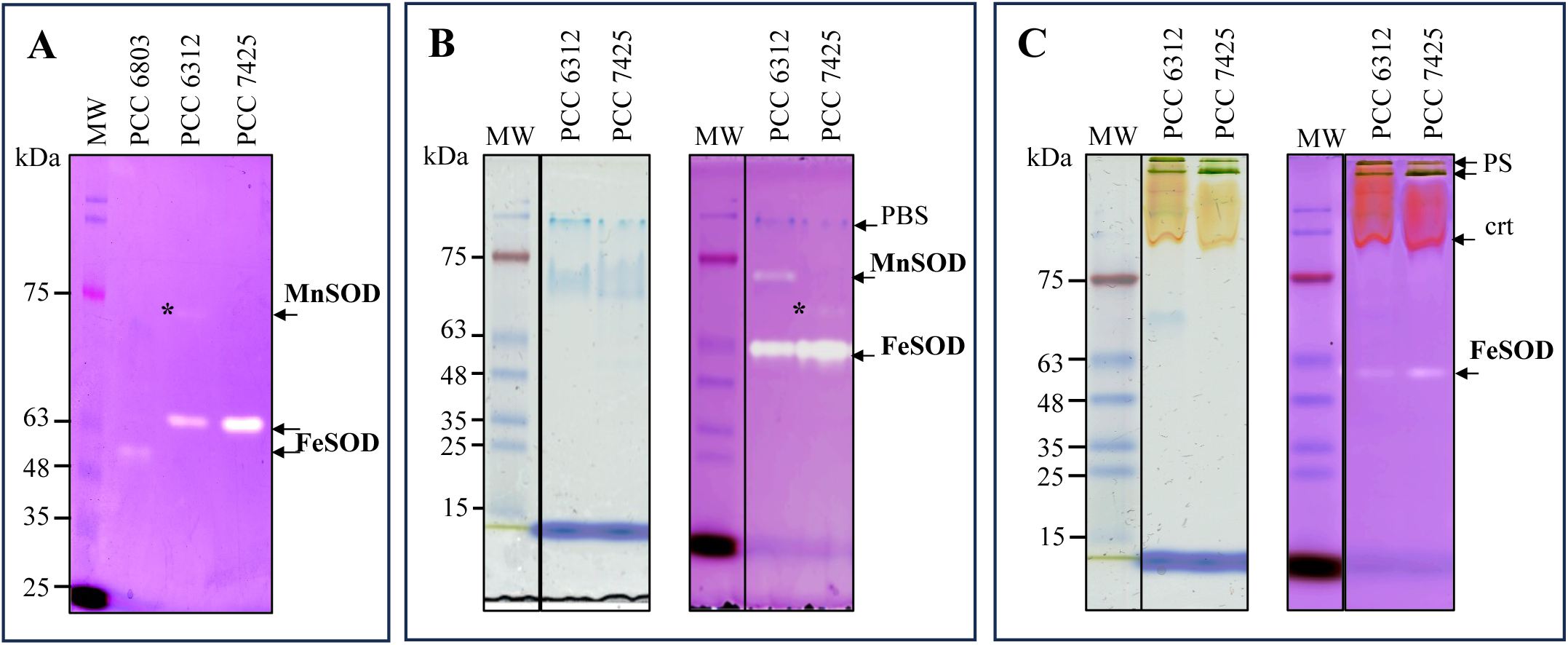
SOD *in-gel* activity assays. (**A**) Total protein fractions from *Synechocystis* sp. PCC 6803, *Synechococcus* sp. PCC 6312, and *Cyanothece* sp. PCC 7425; asterisks mark MnSOD. (**B**) Soluble protein fractions from *Synechococcus* sp. PCC 6312 and *Cyanothece* sp. PCC 7425; asterisks mark the putative truncated MnSOD. PBS, phycobilisomes. (**C**) Membrane protein fractions from the same two strains. PS, photosystems; crt, carotenoid-containing proteins.

### Genetic evidence supporting a dominant role for FeSOD in superoxide detoxification in Cyanothece

Targeted gene deletion was used to assess the respective contributions of *sodA* and *sodB* to superoxide detoxification in *Cyanothece* PCC 7425. A ΔsodB null mutant had previously been generated [17], and a ΔsodA mutant was constructed, using the recently established CRISPR-Cas12a genome-editing system. The crRNA and repair template were cloned together with the Ω (Sp/Sm) cassette into the pSL2680 vector, generating the pSL-ΔsodA plasmid, which was subsequently introduced into *Cyanothece* PCC 7425 by conjugation, as previously described [17]. Careful PCR analysis (45 amplification cycles) followed by NaeI restriction digestion was used to verify allele replacement and complete chromosome segregation. For the ΔsodA mutant, the 2538-bp PCR product was cleaved into 1499 and 1044 bp fragments, as expected from the presence of the NaeI site introduced withion the Ω (Sp/Sm) cassette, whereas the 2065-bp WT product remained undigested (**Fig. 6A**). Similarly, the 2372-bp ΔsodB product was cleaved into fragments of 1497 and 875 bp fragments, whereas the 1852-bp WT product was not cleaved (**Fig. 6B**). No WT size PCR products were detected in either mutant confirming complete chromosome segregation of the ΔsodA and ΔsodB alleles.

**Figure 6.**
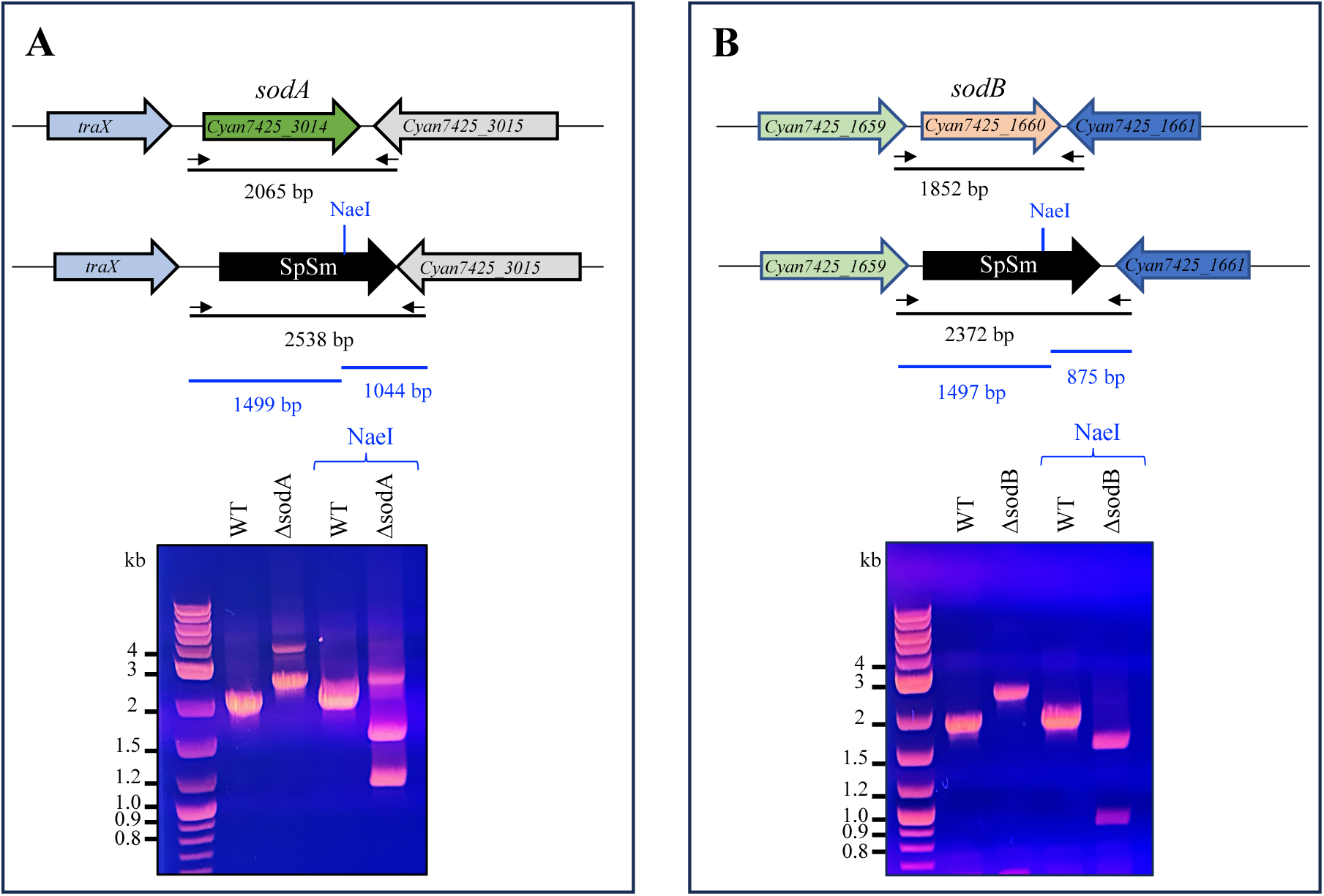
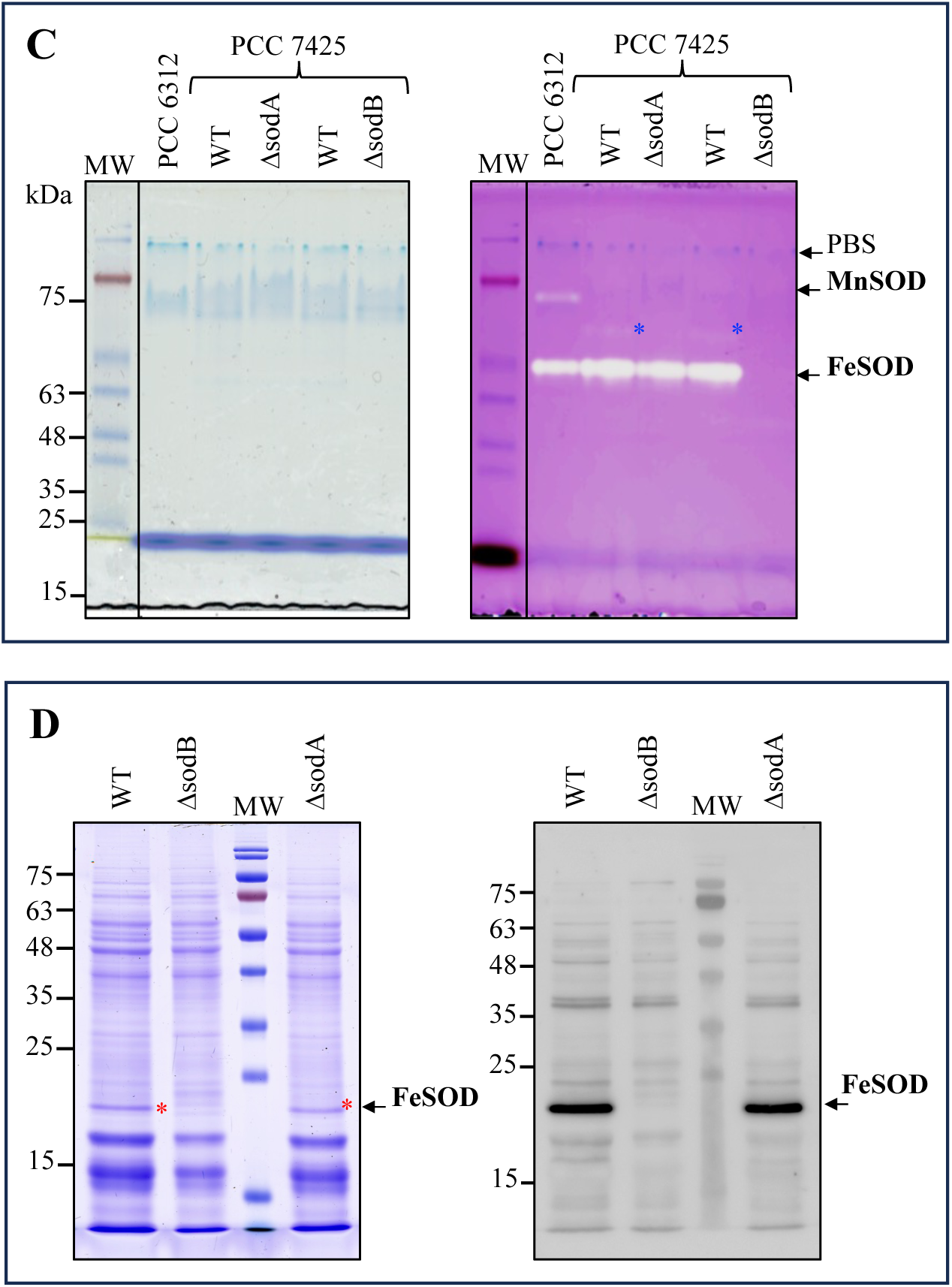
Generation and characterisation of Δsod mutants in *Cyanothece* sp. PCC 7425. (**A-B**) The *sodA* and *sodB* loci and flanking genes, showing replacement of each *sod* gene with the SpSm resistance cassette; agarose gels show the corresponding PCR products (primer positions indicated by arrows) and restriction digestion profiles. (**C**) Non-denaturing PAGE and SOD *in-gel* activity of soluble protein fractions from WT and Δsod mutants, with *Synechococcus* sp. PCC 6312 as a control; asterisks mark the putative truncated MnSOD. PBS, phycobilisomes. (**D**) Coomassie-stained SDS-PAGE and Western blot of total protein fractions from WT and Δsod mutants; asterisks mark FeSOD.

SOD assay was further used to confirm the deletion of the *sod* genes and also to check whether in absence of FeSOD, MnSOD could be induced and more active. As compared to the WT strain, the FeSOD-activity band was completely absent from the Δ*sodB* mutant, confirming *sodB* deletion **(Fig. 6C)**. In this mutant, no band corresponding to MnSOD could be detected. In the ΔsodA mutant, the FeSOD band was readily detected, whereas the above-mentioned band tentatively assigned to the truncated soluble form of MnSOD protein was not observed. However, this putative MnSOD band was also absent in the ΔsodB mutant, preventing its unambiguous assignment to MnSOD. Thus, while the SOD activity profile clearly confirms the loss of FeSOD activity in the ΔsodB strain, the identity of the putative MnSOD band remains uncertain. Coomassie staining, and Western blot analyses of total cellular extract of *Cyanothece* PCC 7425 with commercial antibodies raised against the *Synechocystis* PCC 6803 FeSOD enzyme, were further performed to verify that the FeSOD protein is abundant in *Cyanothece* PCC 7425 WT and its ΔsodA mutant, whereas it is totally absent in the ΔsodB mutant **(Fig. 6D)**. The antibodies did not show any detectable cross reactivity with MnSOD, most likely due to their specificity for FeSOD. Collectively, the genetic, enzymatic and immunochemical analyses, establish FeSOD as the major detectable SOD in *Cyanothece* PCC 7425, whereas the contribution and physiological role of MnSOD remain to be determined.

### FeSOD, but not MnSOD, is crucial for photoautotrophic growth under high-light conditions

To assess the contribution of MnSod and FeSOD to the growth of *Cyanothece* PCC 7425 under different conditions, WT, ΔsodA and ΔsodB cultures from mid-log phase growth were spotted onto solid mineral medium and incubated under either standard (2500 lux) or high-light (8000 lux) conditions. The results (**Fig. 7A**) showed that growth of the ΔsodA mutant was unaffected under both light conditions, whereas growth of the ΔsodB mutant was impaired under standard-light and completely inhibited by high-light. These results indicate that FeSOD, but not MnSOD, is required for growth under high-light conditions that triggers photooxidative stress. To further assess the sensitivity of the sod mutants to oxidative stress, strains were subjected to a menadione (superoxide generator) using a disc diffusion assay. Under standard-light conditions, the WT strain exhibited a clear inhibition zone of 7 mm in diameter in the presence of 5 mM menadione. The ΔsodA mutant displayed a slightly larger inhibition zone of 8.5 mm, indicating a modest increase in menadione sensitivity. In contrast, the ΔsodB mutant displayed a markedly larger sensitivity, with an inhibition zone of 14 mm in diameter (**Fig. 7B**). Under high-light conditions, the WT and ΔsodA strains exhibited comparable inhibition zones, though a small additional inhibition zone was observed for the ΔsodA mutant, again indicating its slightly higher stress sensitivity. In contrast, growth of the ΔsodB mutant was completely inhibited under this condition. Together, these results show that loss of FeSOD markedly increases sensitivity to high-light and superoxides, whereas loss of MnSOD has comparatively minor effects.

**Figure 7.**
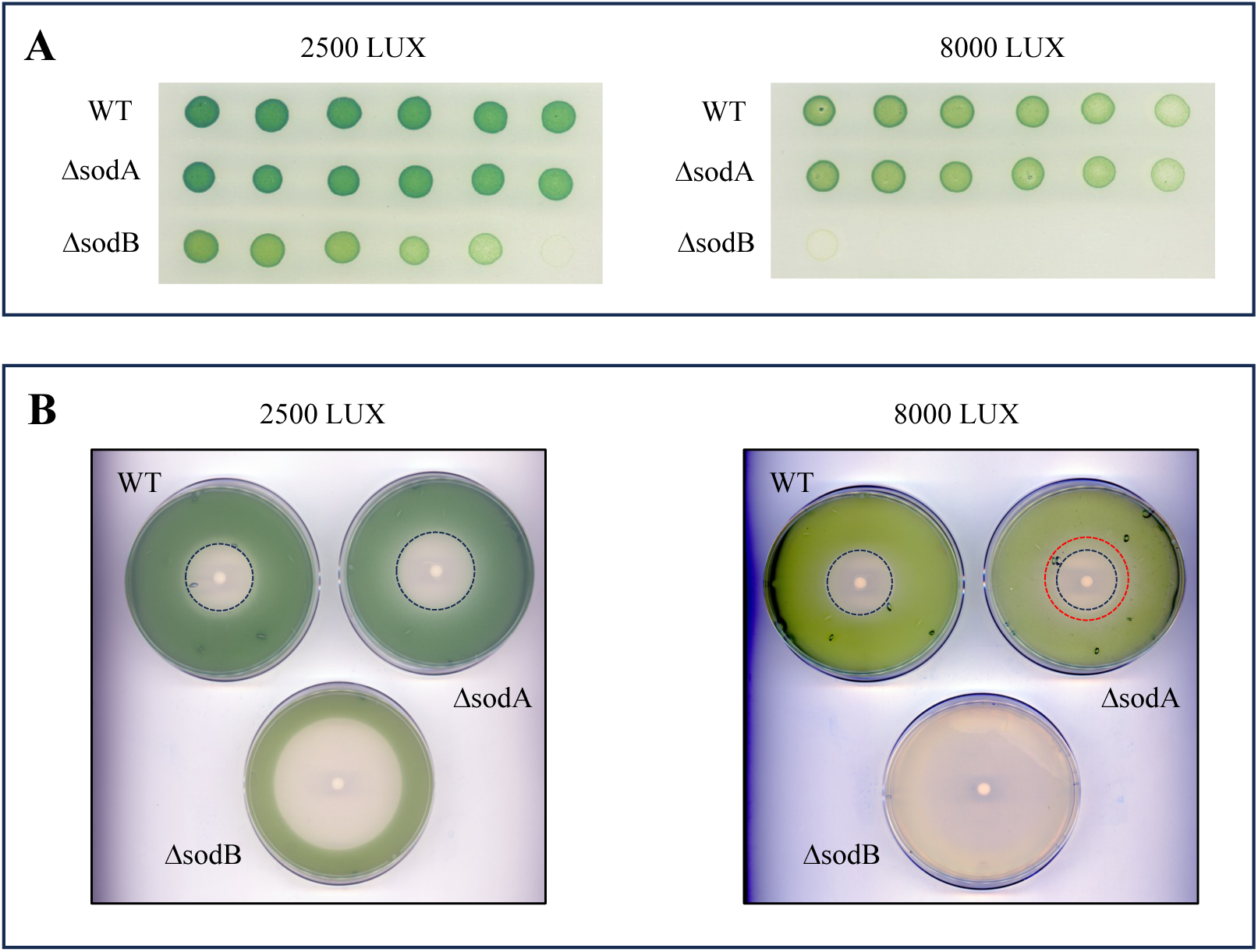
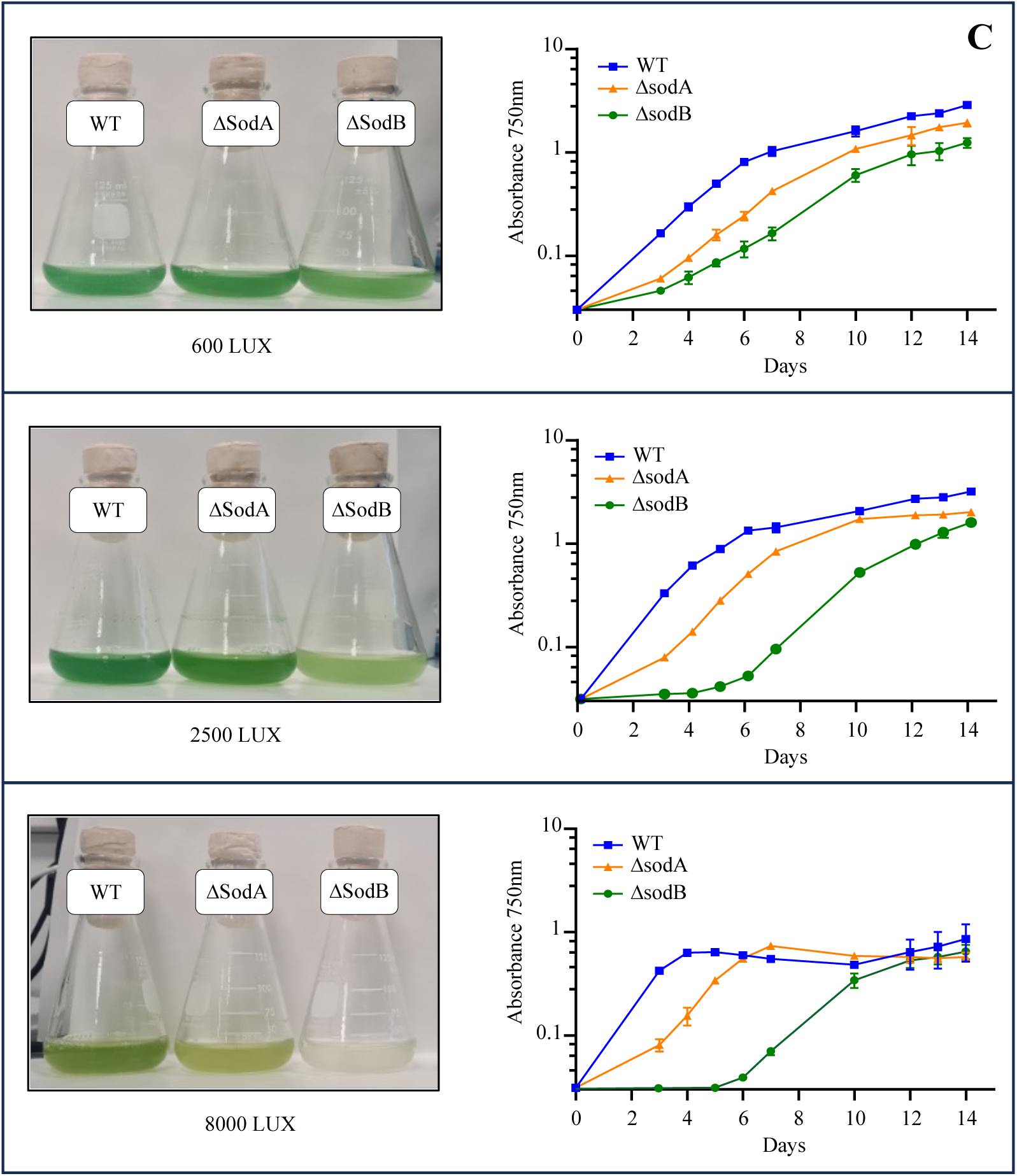
Light intensity and superoxide stress define the growth of sod mutants. (**A**) Serial dilutions of WT and Δsod mutants spotted on solid mineral medium under standard (2500 lux) or high-light (8000 lux) conditions. (**B**) Disc diffusion assay comparing sensitivity of WT and Δsod mutants to menadione under standard and high-light conditions. (**C**) Growth curves of WT and Δsod mutants in liquid mineral medium under low (600 lux), standard, or high-light conditions, showing the increasing light sensitivity of Δsod mutants.

### Light sensitivity of the ΔsodB strain is consistent with superoxide production at the PSI acceptor side (cytoplasm)

It is thought that the acceptor side of PSI, which faces the cytoplasm in cyanobacteria, can generate O_2_^.-^ through the transfer of electrons to O₂ (**Fig. 1**), particularly when downstream electron acceptors become over-reduced, as described by the Mehler reaction [4,31]. Superoxide may also be generated at the lumenal side of the thylakoid membrane, where it has been proposed to arise from PSII activity [32]. Given the distinct subcellular localization of MnSOD and FeSOD, comparison of the corresponding mutant phenotypes can provide insights into the cellular sites of superoxide production. Under low-light intensity, the WT strain exhibited the fastest growth, with a doubling time of approximately 32 h, followed by ΔsodA (34 h) and ΔsodB (42 h) (**Table S4**), with all three strains displaying comparable pigmentation (**Fig. 7C**). However, both mutants exhibited a similar lag phase of two to three days before the onset of growth. Under standard-light condition, the growth of the WT and the ΔsodA strains was faster (doubling times of 26h and 28 h, respectively), than under low-light, whereas the ΔsodB mutant displayed a more pronounced growth defect. Its growth resumed only after approximately five-six days, and its culture exhibited a yellowish color typical of light-sensitive cyanobacterial cells. Once growth was initiated, the ΔsodB mutant exhibited a growth rate comparable to those of the WT and ΔsodA strains, with doubling times of approximately 29h. Under high-light intensity, all strains were negatively affected but the effect was moderate in the WT strain (no growth delay, but the culture turned yellowish with an OD max ≤ 1 unit indicating a lower production of biomass) the doubling time was around 21h. The ΔsodA (small increase in growth delay, changes in color and biomass production respectively higher or similar to those of WT cells) showed a doubling time of 25h. The ΔsodB mutant showed a more pronounced growth delay, a doubling time of 31h and cultures frequently remained whitish throughout the experiment. In some cases, ΔsodB cultures resumed growth, suggesting the possible emergence of suppressors capable of growing in absence of FeSOD under high-light stress. Overall, the growth phenotype of the ΔsodA and ΔsodB mutants became progressively more pronounced with increasing light intensity. Whereas, ΔsodA exhibited a detectable phenotype primarily under high-light stress, ΔsodB displayed a pronounced growth defect under standard-light and an even stronger defect under high-light. These results demonstrate that FeSOD makes a major contribution to growth under light-induced oxidative stress, whereas the contribution of MnSOD appears to be lower. Given the cytoplasmic localization of FeSOD, the pronounced growth impairment of the ΔsodB mutant is consistent with a light enhanced O_2_^.-^ production at the acceptor side of PSI.

The relatively minor contribution of MnSOD to the high-light stress response in *Cyanothece* PCC 7425 is consistent with the loss of MnSOD in many cyanobacteria, including the well-studied models *Synechocystis* PCC 6803 and *Synechococcus* sp. PCC 7942 [13,14]. The differential equipment in FeSOD and MnSOD proteins observed in cyanobacteria is discussed below.

### The N-Ter lipoprotein-binding motif of MnSOD mediates toxicity in heterologous hosts

Heterologous expression provides a useful approach for the functional characterization of genes and proteins. Therefore, we attempted to clone the full-length *sodA* gene of *Cyanothece* PCC 7425 into the RSF1010-derived pCK replicative plasmid which allows strong constitutive expression in *E. coli* and cyanobacteria [33]. All attempts were unsuccessful, suggesting that MnSOD may impair *E. coli* viability. To test this hypothesis, *sodA* was cloned into pFC1K, a pCK related plasmid for tight temperature-controlled expression of studied genes [34]. At the permissive temperature (28°C), repression of the promoter allowed the recovery of colonies harboring the pFC1K*sodA* plasmid. In contrast, no colonies were recovered at the non-permissive temperature (37°C), at which promoter activation drives *sodA* expression, indicating that *sodA* expression prevents colony formation under these conditions (**Fig. S7**). As controls, the genes encoding the *Cyanothece* PCC 7425 FeSOD enzyme and the *Synechococcus* PCC 6312 MnSOD protein were also cloned in pFC1K and transformed to *E. coli*, yielding colonies that grew equally well at both 28°C and 37°C. The absence of toxicity in *E. coli* of these two proteins is correlated to their lack of a membrane anchoring domain, suggesting that the N-terminal membrane anchoring domain of the *Cyanothece* PCC 7425 MnSOD protein may be responsible for its toxicity. To test this hypothesis, this N-terminal membrane anchoring domain was either deleted from the WT MnSOD protein to generate a soluble form of MnSOD, or fused to the soluble FeSOD protein (**Fig. S8**). Deletion of the N-terminal domain abolished MnSOD toxicity, whereas its fusion to FeSOD conferred toxicity, confirming that this N-terminal membrane anchoring domain of MnSOD mediates toxicity in *E. coli*. We next tested whether this toxicity was specific to *E. coli* or also extendable to cyanobacteria. Plasmids expressing either FeSOD or the truncated, soluble form of *Cyanothece* PCC 7425 MnSOD did not impair growth of *Synechocystis* PCC 6803, *Synechococcus* PCC 7942 or *Cyanothece* PCC 7425. In contrast, WT *Cyanothece* PCC 7425 MnSOD or its N-terminal membrane anchoring domain fused to FeSOD inhibited growth at 37°C of *Synechocystis* PCC 6803 and *Synechococcus* PCC 7942, but not in *Cyanothece* 7425, whereas all strains grew well at 28°C. These results indicate that the N-terminal membrane anchoring domain of the *Cyanothece* PCC 7425 MnSOD protein, which contains a predicted lipoprotein binding motif, mediates toxicity in heterologous hosts (*E. coli* and the model cyanobacteria *Synechocystis* PCC 6803 and *Synechococcus* PCC 7942). The host-dependent toxicity of the *Cyanothece* PCC 7425 MnSOD protein may reflect differences in the lipoprotein processing or trafficking machineries of these organisms, potentially leading to interferences with the maturation of essential host lipoproteins.

### The ΔsodB mutant is a suitable host to investigate adaptation to high-light-induced oxidative stress

It is well established that oxidative stress promotes damage and mutagenesis to DNA, particularly in mutants deficient in reactive oxygen species scavenging enzymes [35]. In the course of this study, we observed that prolonged exposure of ΔsodB liquid cultures to high-light stress resulted in the accumulation of cells capable of growing under these otherwise non-permissive conditions. This observation suggested that high-light induced oxidative stress may promote the emergence of adaptive suppressors within the ΔsodB population. To test this hypothesis, ΔsodB cells were grown in liquid cultures under low or high-light conditions for four days, 10 μL were subsequently spotted onto plates maintained under the corresponding light conditions. Cells originating from the low-light culture grew uniformly on the plate, forming a homogeneous green spot (**Fig. 8A**). In contrast, the high-light culture gave rise to isolated colonies displaying different colors, which were able to thrive under this harsh condition, unlike the parental ΔsodB mutant. Colonies exhibiting either a greener than WT phenotype or a yellow coloration were selected (**Fig. 8B**), and their ability to grow in liquid culture under high-light conditions was confirmed (**Fig. 8B**). As compared to the WT strain, these mutant suppressors exhibited improved growth under high-light, but reduced growth under standard-light condition. PCR analysis confirmed that the *sodB* locus remained fully deleted in all selected clones, indicating that they had likely acquired compensatory mutations at secondary genetic loci. Characterization of these mutants is ongoing to identify the causative mutations and elucidate the underlying mechanisms that mitigate the high-light lethality associated with the absence of cytoplasmic superoxide dismutase in cyanobacteria.

**Figure 8.**
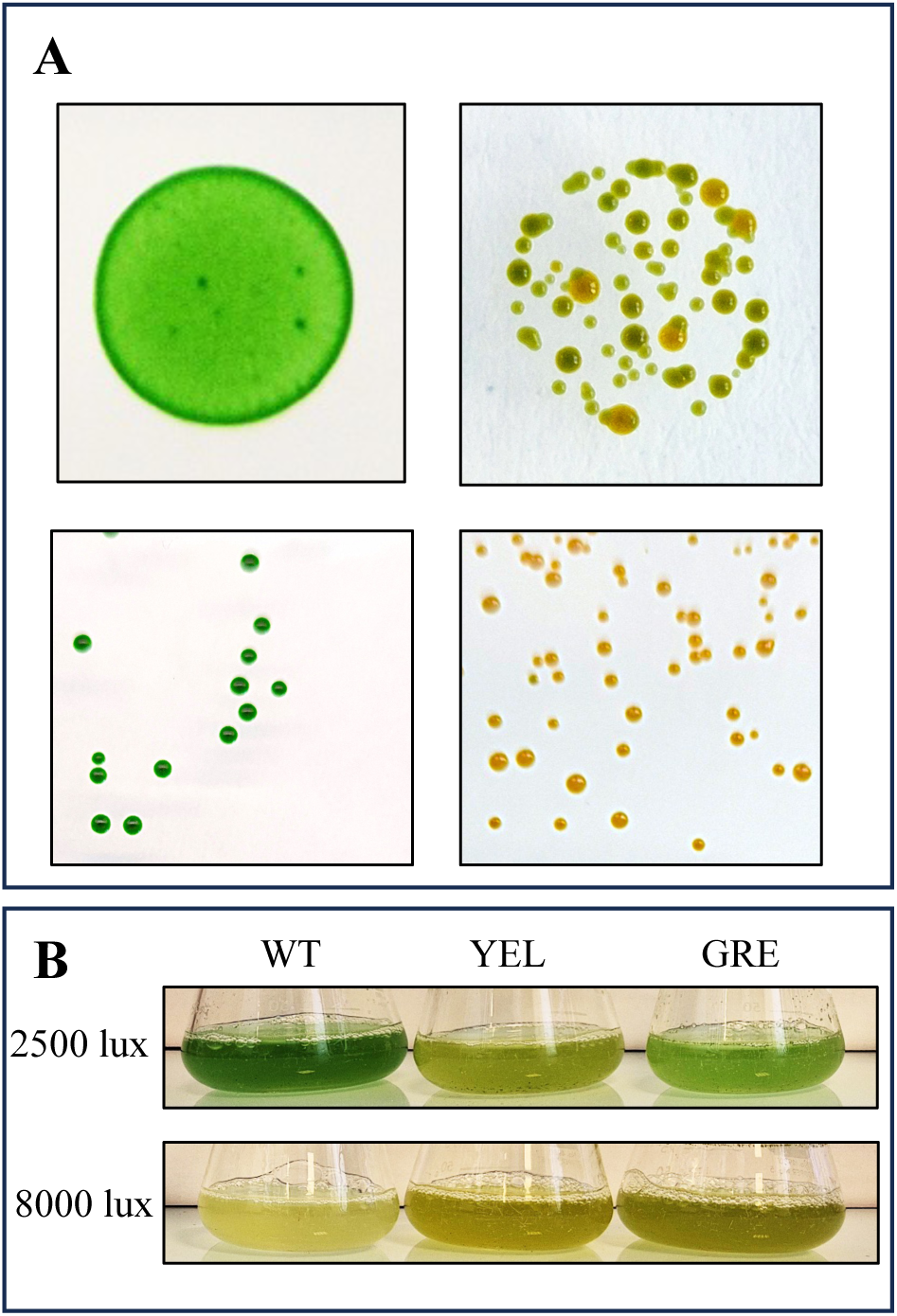
Suppressor selection restores high-light growth in ΔsodB. (**A**) ΔsodB cells spotted on solid mineral medium under standard or high-light conditions; colonies able to grow under high-light were selected as putative suppressors, and representative green and yellow suppressors isolated and propagated under high-light. (**B**) Growth of WT and the green/yellow *sodB* suppressors in liquid mineral medium under standard and high-light conditions.

## Discussion

In oxygenic photosynthetic organisms, high-light can drive photosynthetic electron transport beyond the capacity of downstream electron acceptors, resulting in over-reduction of the photosynthetic electron transport chain and increased production of ROS, including O_2_^.-^, thereby necessitating robust antioxidant defenses [36–39]. Phototrophs consequently possess diverse SOD repertoires that differ in number, subcellular localization, and metal cofactor. Cyanobacteria harbor Fe, Mn, CuZn, and/or Ni-dependent SODs that can be localized to the cytoplasm, periplasm, thylakoid membrane, or lumen [6,12]. This diversity is reminiscent of plants, which possess three major SOD classes with distinct subcellular distributions. FeSOD and CuZnSOD occur in the chloroplast stroma and cytosol, whereas MnSOD is localized to mitochondria and is notably absent from chloroplasts [7,8]. In this study, we investigated the sequence diversity and putative subcellular localization of cyanobacterial MnSOD and compared its functional importance with that of FeSOD in *Cyanothece* PCC 7425. The results raise two related questions: why has MnSOD been recurrently lost in some cyanobacterial lineages, and why is MnSOD absent from plant chloroplasts despite its presence in some cyanobacteria?

One possible explanation involves the relative availability and cellular allocation of Fe and Mn. Cyanobacteria maintain substantial and tightly regulated intracellular Fe pools, which may favor the maintenance of Fe dependent SOD while limiting competition between Fe and Mn. This may be particularly relevant in the thylakoid membrane, where Mn must be preferentially allocated to the oxygen evolving complex of photosystem II. Maintenance of a Mn-dependent antioxidant enzyme could therefore impose an additional demand on a metal with an essential role in photosynthetic water oxidation. However, metal availability alone is unlikely to account for the differential retention of FeSOD and MnSOD. The spatial distribution of ROS production within the photosynthetic electron transport chain provides an additional, and potentially more important, explanation. The acceptor side of PSI is considered a major site of O_2_^.-^ production through the reduction of molecular oxygen via the Mehler reaction. Consistent with an important role for FeSOD in this context, the *Cyanothece* PCC 7425 ΔsodB mutant was highly sensitive to high-light and menadione, while complete deletion of *sodB* could not be achieved in the well-established cyanobacterial model *Synechocystis* PCC 6803. In plants, loss of either cytosolic CuZnSod or FeSOD1 has comparatively very limited effects on growth, because O_2_^.-^ is unable to cross the chloroplast membranes ; whereas loss of the chloroplast FeSOD isoforms FeSOD2 and FeSOD3 causes severe defects in chloroplast development and plant growth. Together, these observations indicate that SOD function depends strongly on subcellular localization and suggest that proximity to sites of ROS generation is an important determinant of physiological relevance. A compelling rationale for the conservation of FeSOD in cyanobacteria is provided by the structure of PSI and the arrangement of the cofactors, in particular the 4Fe-4S clusters. The acceptor side of PSI faces the cytoplasmic side of the thylakoid membrane, placing FeSOD in a favorable position to intercept O_2_^.-^ generated in association with PSI. This proximity may be particularly important for protecting the vulnerable 4Fe-4S cofactors of PSI [40]. Superoxide can damage susceptible 4Fe-4S clusters [40], resulting in the release of redox-active ferrous iron. The liberated Fe can subsequently participate in Fenton chemistry and promote the formation of highly reactive hydroxyl radicals, thereby amplifying oxidative damage to nearby macromolecules, including DNA. FeSOD may therefore provide an important first line of defense against superoxide in the cytoplasmic compartment, limiting both direct damage to PSI 4Fe-4S cofactors and secondary oxidative reactions. By contrast, the selective pressure to retain SOD activity within the thylakoid lumen may be weaker. If O_2_^.-^ production is relatively limited in this compartment and major vulnerable targets, such as exposed 4Fe-4S clusters and DNA, are largely absent, the benefit of local superoxide detoxification would be correspondingly reduced. This situation differs from that in chloroplasts of algae and higher plants, where the diversification and compartmentalization of SOD isoforms may be necessary to control ROS across multiple subcompartments and protect vulnerable components, including the chloroplast genome. Thus, differences in SOD conservation may reflect, at least in part, differences in the spatial relationship between ROS sources and their most susceptible molecular targets.

This framework provides a possible explanation for the contrasting retention of FeSOD and MnSOD in cyanobacteria. FeSOD is well positioned to detoxify O_2_^.-^ associated with PSI, whereas MnSOD may provide a more localized function when associated with the thylakoid membrane or lumen. Consistent with this distinction, ΔsodA mutant of *Cyanothece* PCC 7425 and *Anabaena* sp. PCC 7120 exhibit reduced growth but lack the pronounced high-light hypersensitivity observed in the corresponding ΔsodB mutant. These phenotypes suggest that MnSOD contributes to cellular fitness but is less critical than FeSOD under conditions that strongly promote photosynthetic O_2_^.-^ production. The preferential retention of FeSOD may also help explain the absence of MnSOD from plant chloroplasts. It has been proposed that the MnSOD of cyanobacterial origin was lost during plastid evolution [9,10]. Our findings raise two possibilities: the cyanobacterial lineage that gave rise to primary plastids may already have lacked MnSOD, or MnSOD may have been lost during plastid evolution. In either scenario, the retention of FeSOD would be consistent with an antioxidant system in which protection against O_2_^.-^ generated near the PSI acceptor side was prioritized. This interpretation does not exclude other factors, including metal homeostasis and changes in cellular compartmentalization, from contributing to MnSOD loss.

A related question is why some cyanobacteria, including *Cyanothece* sp. PCC 7425 and *Anabaena* sp. PCC 7120, have retained a thylakoid-associated MnSOD. Its localization may provide a functional explanation. MnSOD associated with the thylakoid membrane and/or lumen could contribute to the detoxification of O_2_^.-^ generated near PSII or the cytochrome *b_6_f* complex. We therefore propose that electron leakage to O_2_ at or near these components may represent a secondary, but physiologically relevant, source of O_2_^.-^ that is locally controlled by MnSOD in MnSOD containing cyanobacteria. In lineages lacking MnSOD, such O_2_^.-^ production may be sufficiently low, or tolerated by intrinsically more robust lumenal components, to permit loss of this enzyme without impairing photosynthetic fitness. The structure of PSI further supports this interpretation. The FA and FB 4Fe-4S clusters are located on the stromal/cytoplasmic side of the thylakoid membrane and are therefore spatially separated from O_2_^.-^ generated within the lumen [41]. Consequently, lumenal MnSOD may provide limited direct protection to these vulnerable PSI cofactors. This spatial separation could contribute to the weaker selective pressure for maintaining lumenal SOD activity, while alternative antioxidant mechanisms may provide additional protection against ROS generated on the donor side of PSII. For example, APE1, a conserved thylakoid-bound Cu-binding protein, has recently been reported to contribute to O_2_^.-^ detoxification in oxygenic photosynthetic organisms [42].

Overall, our findings support a model in which the evolutionary distribution of SODs in photosynthetic organisms reflects the combined effects of metal availability, enzyme properties, and, importantly, subcellular organization. FeSOD appears to occupy a particularly important position because it is localized to the compartment containing major vulnerable targets of PSI-associated superoxide production. MnSOD, in contrast, may be retained in some cyanobacteria when its thylakoid association provides an additional local detoxification capacity, but may be dispensable when the corresponding ROS burden is limited or can be controlled by other mechanisms. Spatial matching between O_2_^.-^ production, antioxidant localization, and the distribution of susceptible molecular targets may therefore represent an important factor shaping the diversification and evolutionary maintenance of antioxidant systems in cyanobacteria and chloroplasts.

## Acknowledgment

We are grateful to Laila Sago and David Cornu from SICaPS platform. This work has benefited from the facilities and expertise of the SICaPS platform of I2BC (Institut de Biologie Intégrative de la Cellule).

## Author contributions

Conceptualization, methodology, data analysis and curation: Kahn M-A, Durand A, Steunou A-S, Leleux S., Cassier-Chauvat C., Chauvat F., and Ouchane S. : Investigations and data analysis: Kahn M-A, Durand A, Steunou A-S. Leleux S. and Ouchane Writing original draft: Ouchane S. Review & editing: Kahn M-A, Durand A., Steunou A-S, Leleux S., Cassier-Chauvat C., Chauvat F., and Ouchane S.

## Supplementary Material

Tables and Supplementray figures are available.

## Data availability statement

Data supporting the findings of this work are available within the paper and its Supplementary Files. The datasets and materials generated and analyzed in the study are available from the corresponding authors upon request.

## Declaration of generative AI-assisted technologies

The authors used AI for language editing purposes only, then reviewed and edited the content as needed and take full responsibility for the content of the publication.

## References

1. Imlay JA. Cellular Defenses against Superoxide and Hydrogen Peroxide. Annu Rev Biochem. 2008;77: 755–776. doi:10.1146/annurev.biochem.77.061606.161055

2. Forquer I, Covian R, Bowman MK, Trumpower BL, Kramer DM. Similar Transition States Mediate the Q-cycle and Superoxide Production by the Cytochrome *bc*1 Complex*. Journal of Biological Chemistry. 2006;281: 38459–38465. doi:10.1074/jbc.M605119200

3. Krieger-Liszkay A, Shimakawa G. Regulation of the generation of reactive oxygen species during photosynthetic electron transport. Biochemical Society Transactions. 2022;50: 1025–1034. doi:10.1042/BST20211246

4. Mubarakshina MM, Ivanov BN. The production and scavenging of reactive oxygen species in the plastoquinone pool of chloroplast thylakoid membranes. Physiologia Plantarum. 2010;140: 103–110. doi:10.1111/j.1399-3054.2010.01391.x

5. Frye KA, Sendra KM, Waldron KJ, Kehl-Fie TE. Old dogs, new tricks: New insights into the iron/manganese superoxide dismutase family. Journal of Inorganic Biochemistry. 2022;230: 111748. doi:10.1016/j.jinorgbio.2022.111748

6. Boden JS, Konhauser KO, Robbins LJ, Sánchez-Baracaldo P. Timing the evolution of antioxidant enzymes in cyanobacteria. Nat Commun. 2021;12: 4742. doi:10.1038/s41467-021-24396-y

7. Gallie DR, Chen Z. Chloroplast-localized iron superoxide dismutases FSD2 and FSD3 are functionally distinct in Arabidopsis. Fukai T, editor. PLoS ONE. 2019;14: e0220078. doi:10.1371/journal.pone.0220078

8. Pilon M, Ravet K, Tapken W. The biogenesis and physiological function of chloroplast superoxide dismutases. Biochimica et Biophysica Acta (BBA) - Bioenergetics. 2011;1807: 989–998. doi:10.1016/j.bbabio.2010.11.002

9. Bowler C, Montagu MV, Inze D. Superoxide Dismutase and Stress Tolerance. Annual Review of Plant Biology. 1992;43: 83–116. doi:10.1146/annurev.pp.43.060192.000503

10. Wolfe-Simon F, Grzebyk D, Schofield O, Falkowski PG. The Role and Evolution of Superoxide Dismutases in Algae. Journal of Phycology. 2005;41: 453–465. doi:10.1111/j.1529-8817.2005.00086.x

11. Allen MD, Kropat J, Tottey S, Del Campo JA, Merchant SS. Manganese Deficiency in Chlamydomonas Results in Loss of Photosystem II and MnSOD Function, Sensitivity to Peroxides, and Secondary Phosphorus and Iron Deficiency. Plant Physiology. 2007;143: 263–277. doi:10.1104/pp.106.088609

12. Harada M, Akiyama A, Furukawa R, Yokobori S, Tajika E, Yamagishi A. Evolution of Superoxide Dismutases and Catalases in Cyanobacteria: Occurrence of the Antioxidant Enzyme Genes before the Rise of Atmospheric Oxygen. J Mol Evol. 2021;89: 527–543. doi:10.1007/s00239-021-10021-5

13. Thomas DJ, Avenson TJ, Thomas JB, Herbert SK. A Cyanobacterium Lacking Iron Superoxide Dismutase Is Sensitized to Oxidative Stress Induced with Methyl Viologen but Is Not Sensitized to Oxidative Stress Induced with Norflurazon. Plant Physiol. 1998;116: 1593–1602. doi:10.1104/pp.116.4.1593

14. Ke W-T, Dai G-Z, Jiang H-B, Zhang R, Qiu B-S. Essential roles of iron superoxide dismutase in photoautotrophic growth of Synechocystis sp. PCC 6803 and heterogeneous expression of marine Synechococcus sp. CC9311 copper/zinc superoxide dismutase within its sodB knockdown mutant. Microbiology. 2014;160: 228–241. doi:10.1099/mic.0.073080-0

15. Tamanna SS, Boden JS, Kaiser KM, Wannicke N, Höring J, Sánchez-Baracaldo P, et al. Early-Branching Cyanobacteria Grow Faster and Upregulate Superoxide Dismutase Activity Under a Simulated Early Earth Anoxic Atmosphere. Geobiology. 2024;22: e70005. doi:10.1111/gbi.70005

16. Li T, Huang X, Zhou R, Liu Y, Li B, Nomura C, et al. Differential Expression and Localization of Mn and Fe Superoxide Dismutases in the Heterocystous Cyanobacterium *Anabaena* sp. Strain PCC 7120. J Bacteriol. 2002;184: 5096–5103. doi:10.1128/JB.184.18.5096-5103.2002

17. Khan MA, Durand A, Skouri-Panet F, Benzerara K, Cassier-Chauvat C, Chauvat F, et al. Targeted genome editing of the non-model cyanobacterium Cyanothece PCC 7425 via CRISPR/Cas12a. Appl Microbiol Biotechnol. 2026 [cited 27 Sept 2026]. doi:10.1007/s00253-026-13959-y

18. Dereeper A, Guignon V, Blanc G, Audic S, Buffet S, Chevenet F, et al. Phylogeny.fr: robust phylogenetic analysis for the non-specialist. Nucleic Acids Res. 2008;36: W465–469. doi:10.1093/nar/gkn180

19. Katoh K, Standley DM. MAFFT Multiple Sequence Alignment Software Version 7: Improvements in Performance and Usability. Mol Biol Evol. 2013;30: 772–780. doi:10.1093/molbev/mst010

20. Capella-Gutiérrez S, Silla-Martínez JM, Gabaldón T. trimAl: a tool for automated alignment trimming in large-scale phylogenetic analyses. Bioinformatics. 2009;25: 1972– 1973. doi:10.1093/bioinformatics/btp348

21. Minh BQ, Schmidt HA, Chernomor O, Schrempf D, Woodhams MD, von Haeseler A, et al. IQ-TREE 2: New Models and Efficient Methods for Phylogenetic Inference in the Genomic Era. Mol Biol Evol. 2020;37: 1530–1534. doi:10.1093/molbev/msaa015

22. Letunic I, Bork P. Interactive Tree Of Life (iTOL) v5: an online tool for phylogenetic tree display and annotation. Nucleic Acids Res. 2021;49: W293–W296. doi:10.1093/nar/gkab301

23. Piddington DL, Fang FC, Laessig T, Cooper AM, Orme IM, Buchmeier NA. ,Zn Superoxide Dismutase of *Mycobacterium tuberculosis* Contributes to Survival in Activated Macrophages That Are Generating an Oxidative Burst. Tuomanen EI, editor. Infect Immun. 2001;69: 4980–4987. doi:10.1128/IAI.69.8.4980-4987.2001

24. Frain KM, Gangl D, Jones A, Zedler JAZ, Robinson C. Protein translocation and thylakoid biogenesis in cyanobacteria. Biochimica et Biophysica Acta (BBA) - Bioenergetics. 2016;1857: 266–273. doi:10.1016/j.bbabio.2015.08.010

25. Okuda S, Tokuda H. Lipoprotein sorting in bacteria. Annu Rev Microbiol. 2011;65: 239–259. doi:10.1146/annurev-micro-090110-102859

26. Teufel F, Almagro Armenteros JJ, Johansen AR, Gíslason MH, Pihl SI, Tsirigos KD, et al. SignalP 6.0 predicts all five types of signal peptides using protein language models. Nat Biotechnol. 2022;40: 1023–1025. doi:10.1038/s41587-021-01156-3

27. Letunic I, Khedkar S, Bork P. SMART: recent updates, new developments and status in 2020. Nucleic Acids Res. 2021;49: D458–D460. doi:10.1093/nar/gkaa937

28. Regelsberger G, Laaha U, Dietmann D, Rüker F, Canini A, Grilli-Caiola M, et al. The Iron Superoxide Dismutase from the Filamentous Cyanobacterium *Nostoc* PCC 7120. Journal of Biological Chemistry. 2004;279: 44384–44393. doi:10.1074/jbc.M406254200

29. Regelsberger G, Atzenhofer W, Rüker F, Peschek GA, Jakopitsch C, Paumann M, et al. Biochemical Characterization of a Membrane-bound Manganese-containing Superoxide Dismutase from the CyanobacteriumAnabaena PCC 7120. Journal of Biological Chemistry. 2002;277: 43615–43622. doi:10.1074/jbc.M207691200

30. Aryal UK, Callister SJ, McMahon BH, McCue L-A, Brown J, Stöckel J, et al. Proteomic profiles of five strains of oxygenic photosynthetic cyanobacteria of the genus Cyanothece. J Proteome Res. 2014;13: 3262–3276. doi:10.1021/pr5000889

31. Kozuleva M, Petrova A, Milrad Y, Semenov A, Ivanov B, Redding KE, et al. Phylloquinone is the principal Mehler reaction site within photosystem I in high-light. Plant Physiol. 2021;186: 1848–1858. doi:10.1093/plphys/kiab221

32. Pospíšil P. Molecular mechanisms of production and scavenging of reactive oxygen species by photosystem II. Biochimica et Biophysica Acta (BBA) - Bioenergetics. 2012;1817: 218–231. doi:10.1016/j.bbabio.2011.05.017

33. Marceau F, Lamothe-Sibold M, Farci S, Ouchane S, Cassier-Chauvat C, Chauvat F. First Characterization of a Cyanobacterial Xi-Class Glutathione S-Transferase in Synechocystis PCC 6803. Antioxidants. 2024;13: 1577. doi:10.3390/antiox13121577

34. Ortega-Ramos M, Jittawuttipoka T, Saenkham P, Czarnecka-Kwasiborski A, Bottin H, Cassier-Chauvat C, et al. Engineering Synechocystis PCC6803 for hydrogen production: influence on the tolerance to oxidative and sugar stresses. PLoS One. 2014;9: e89372. doi:10.1371/journal.pone.0089372

35. Imlay JA. The molecular mechanisms and physiological consequences of oxidative stress: lessons from a model bacterium. Nat Rev Microbiol. 2013;11: 443–454. doi:10.1038/nrmicro3032

36. Krieger-Liszkay A, Shimakawa G. Regulation of the generation of reactive oxygen species during photosynthetic electron transport. Biochem Soc Trans. 2022;50: 1025–1034. doi:10.1042/BST20211246

37. Foyer CH. Reactive oxygen species, oxidative signaling and the regulation of photosynthesis. Environmental and Experimental Botany. 2018;154: 134–142. doi:10.1016/j.envexpbot.2018.05.003

38. Mullineaux PM, Exposito-Rodriguez M, Laissue PP, Smirnoff N. ROS-dependent signalling pathways in plants and algae exposed to high-light: Comparisons with other eukaryotes. Free Radic Biol Med. 2018;122: 52–64. doi:10.1016/j.freeradbiomed.2018.01.033

39. Latifi A, Ruiz M, Zhang C-C. Oxidative stress in cyanobacteria. FEMS Microbiol Rev. 2009;33: 258–278. doi:10.1111/j.1574-6976.2008.00134.x

40. Photodamage of iron–sulphur clusters in photosystem I induces non-photochemical energy dissipation | Nature Plants. [cited 1 Oct 2026]. Available: https://www-nature-com.insb.bib.cnrs.fr/articles/nplants201635

41. Fromme P, Jordan P, Krauß N. Structure of photosystem I. Biochimica et Biophysica Acta (BBA) - Bioenergetics. 2001;1507: 5–31. doi:10.1016/S0005-2728(01)00195-5

42. Malesinski S, Vidal-Meireles A, Giovannetti E, Chazaux M, Krieger-Liszkay A, Boderiou C, et al. Acclimation of Photosynthesis to the Environment 1 acts as a copper-binding superoxide detoxification enzyme. Nat Commun. 2026;17: 5192. doi:10.1038/s41467-026-71893-z

