## Supplemental Figures and Tables for "MnSOD Diversifies, while FeSOD Dominates Superoxide Defense in Cyanobacteria"

### Supplementary material
#### (Figures)

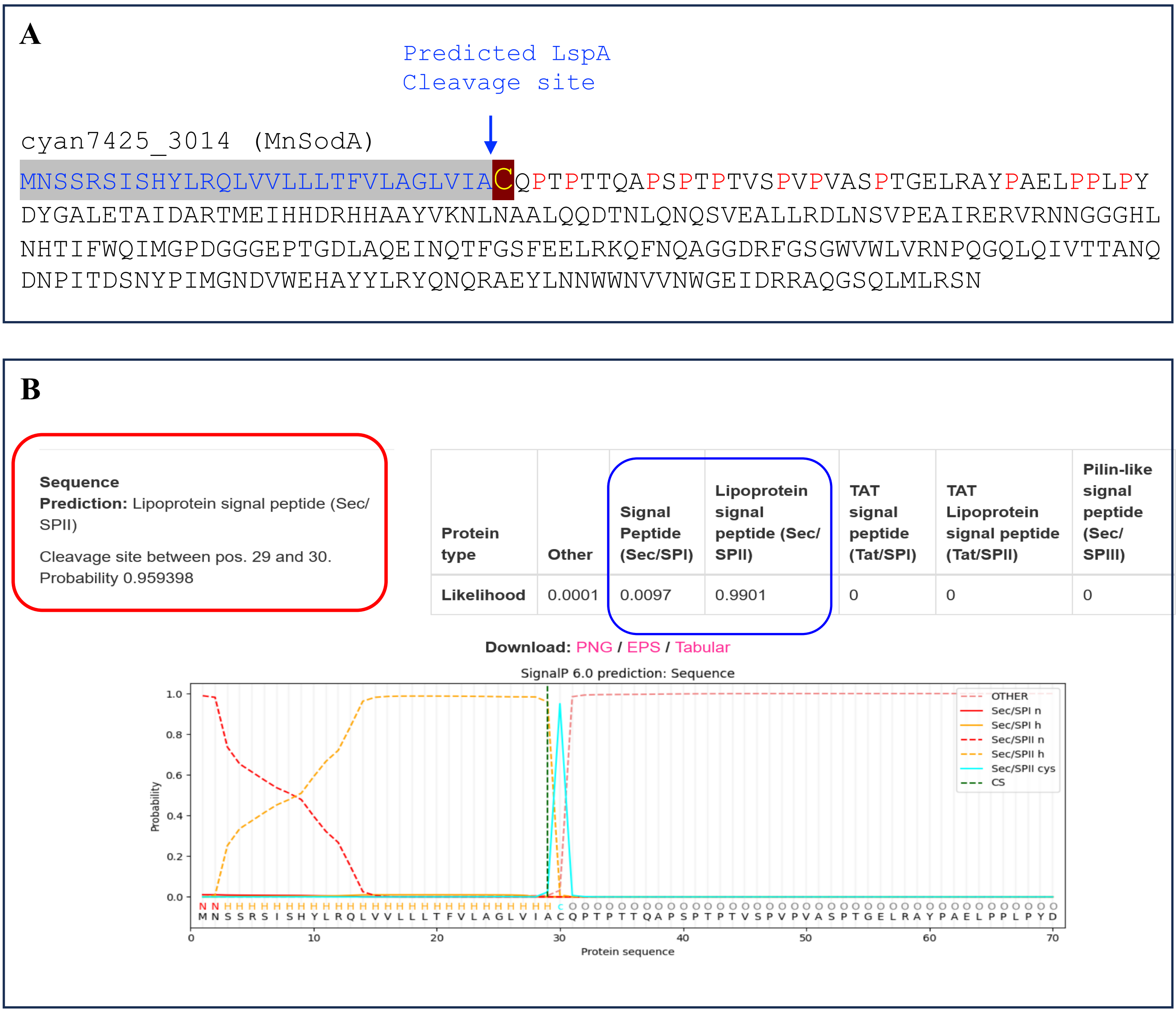

| Protein type | Other | Signal Peptide (Sec/SPI) | Lipoprotein signal peptide (Sec/SPII) | TAT signal peptide (Tat/SPI) | TAT Lipoprotein signal peptide (Tat/SPII) | Pilin-like signal peptide (Sec/SPIII) |
|---|---|---|---|---|---|---|
| Likelihood | 0.0001 | 0.0097 | 0.9901 | 0 | 0 | 0 |

**Fig. S1**: (**A**) Sequence of MnSodA showing the lipo-signal peptide with the concerved cysteine. (**B**) SignalP 6.0 predicted profile of MnSodA from *Cyanothece* PCC 7425.

Blast SodA against 1000 genomes of Cyanobacteria

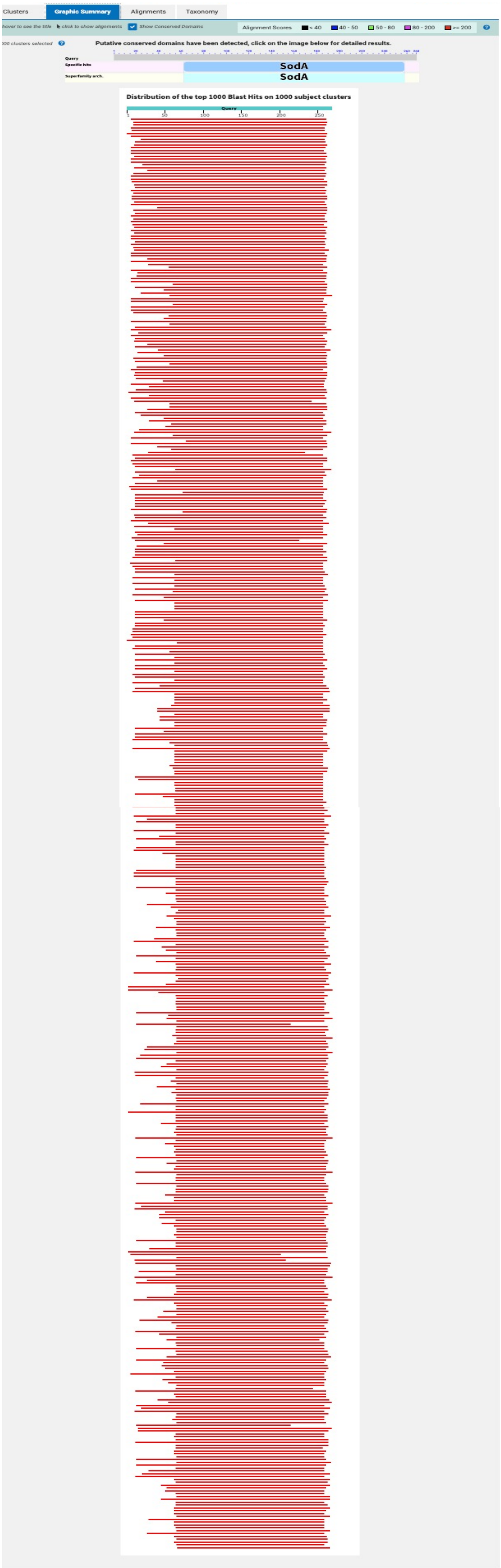

**Fig. S2:** BLASTP result profile showing the N-terminal extension in cyanobacterial MnSODs similar to that of *Cyanothece* PCC 7425 MnSodA.

Blast SodA against 5000 procaryote genomes (excluding cyanos)

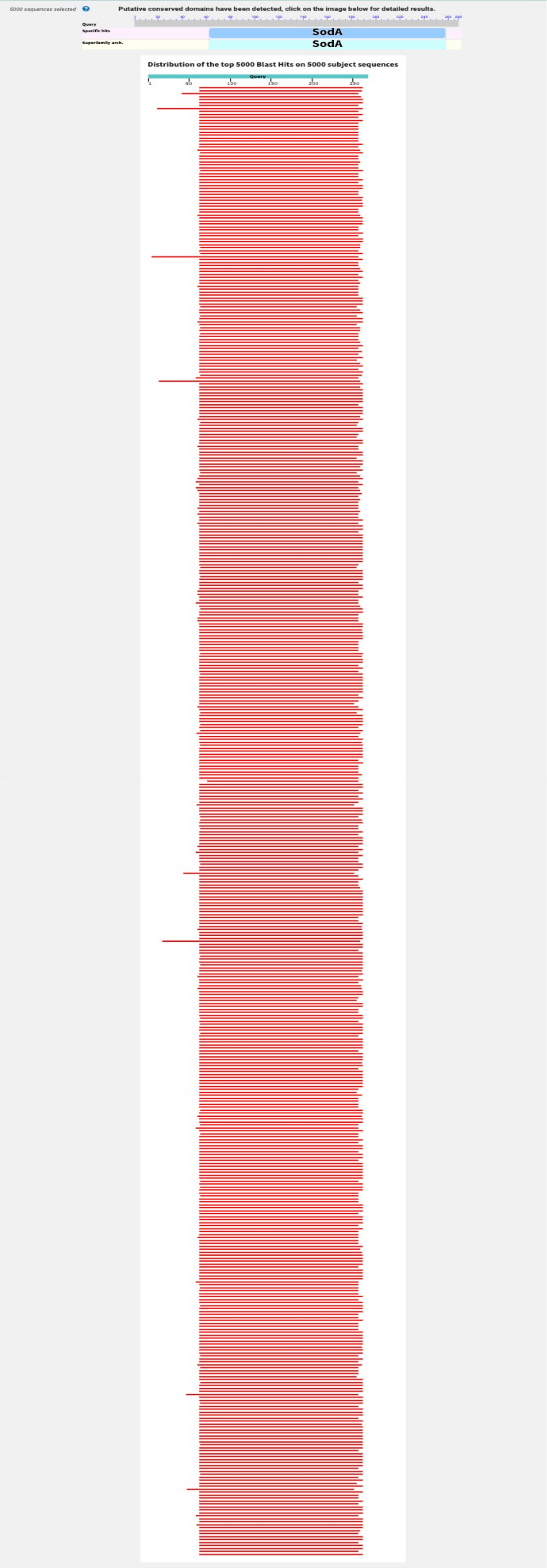

**Fig. S2:** BLASTP of *Cyanothece* PCC 7425 MnSodA against procaryote genomes.

Blast SodA against 322 genomes of Algae

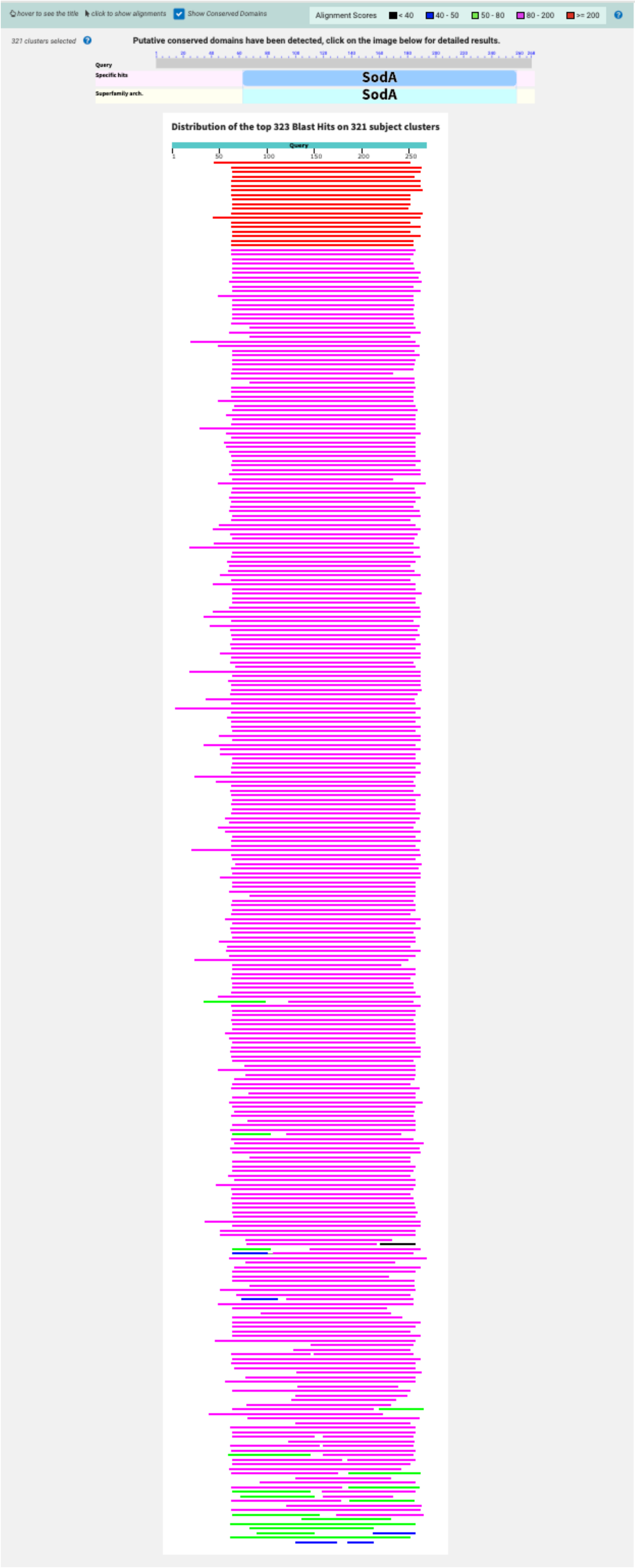

**Fig. S2:** BLASTP of *Cyanothece* PCC 7425 MnSodA against algal genomes.

Blast SodA against 1005 Plants genomes

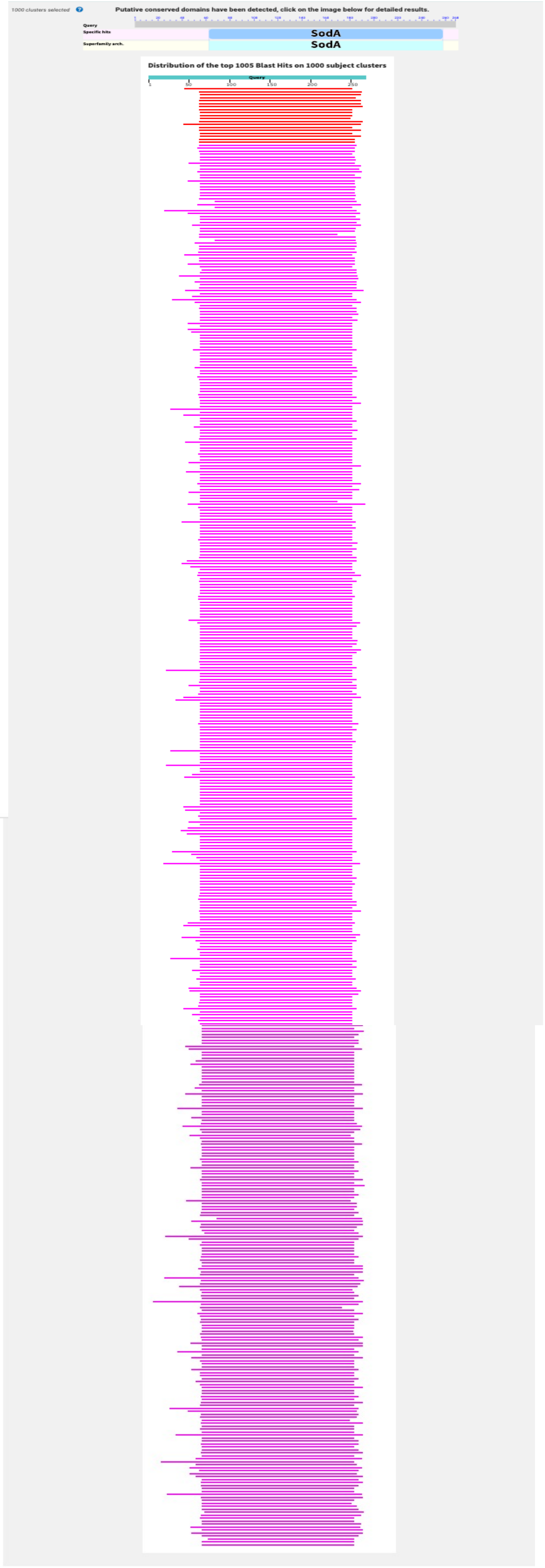

**Fig. S2:** BLASTP of *Cyanothece* PCC 7425 MnSodA against higher plants genomes

### Blast SodA against 5000 Eucaryotic genomes

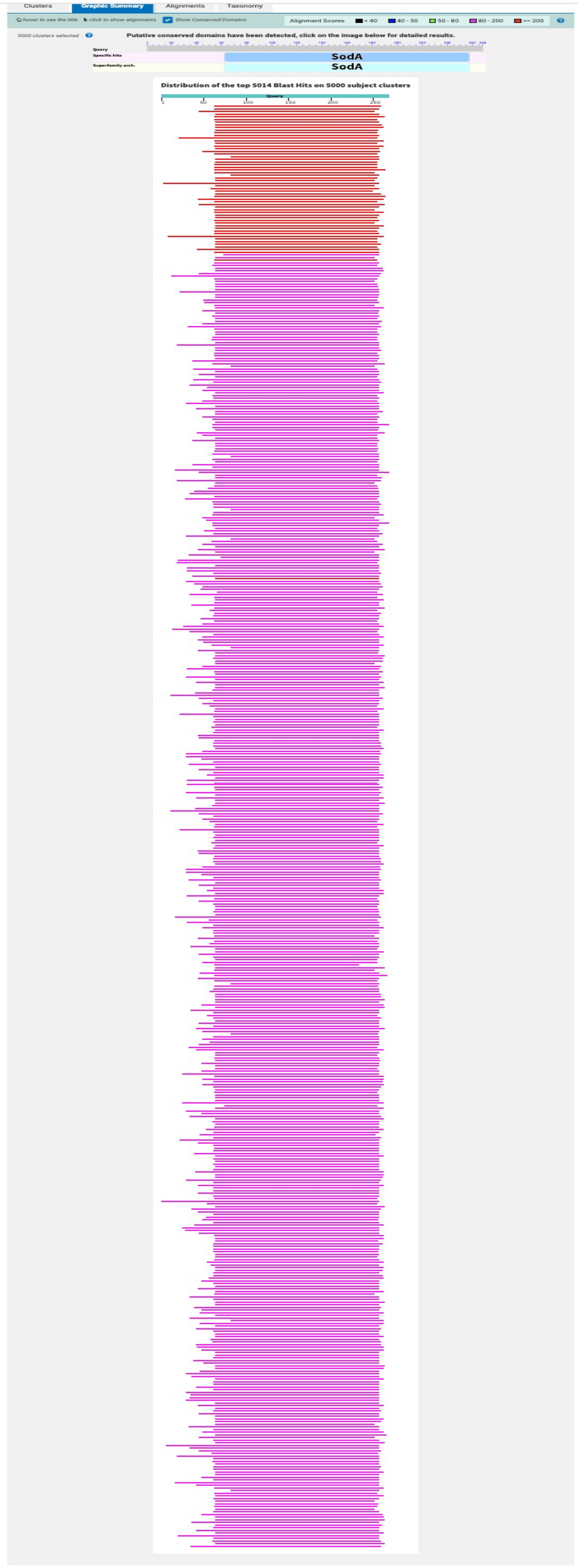

SodA blast against purple photosynthetic bacteria

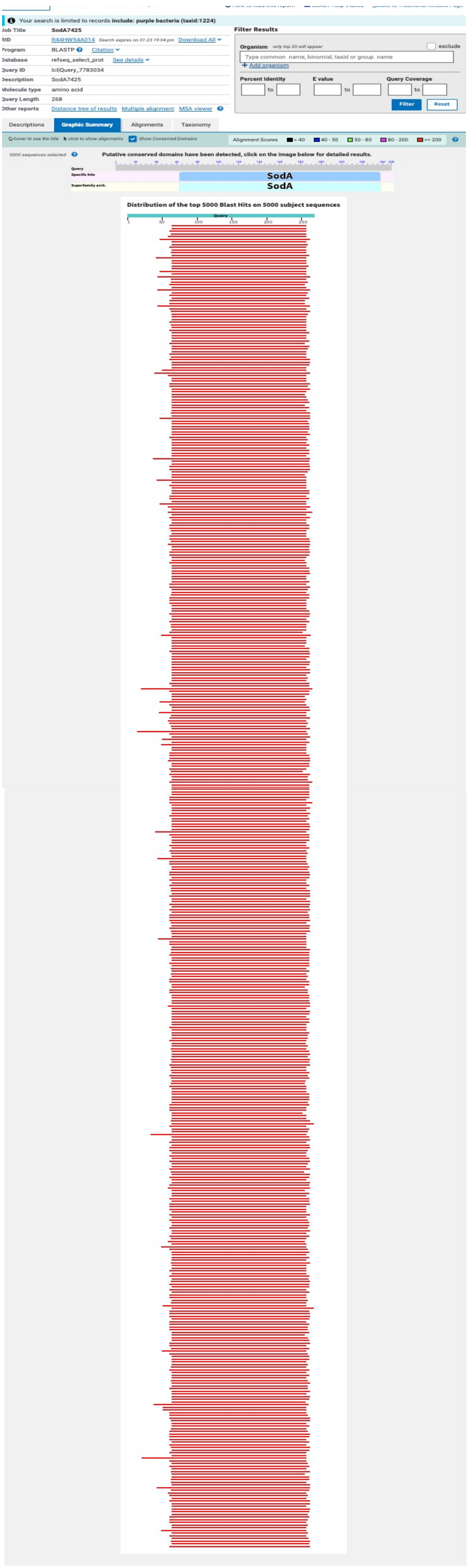

**Fig. S2:** BLASTP of *Cyanothece* PCC 7425 MnSodA against photosynthetic purple bacteria genomes

SodA blast against Green sulphur and non-sulphur photosynthetic bacteria

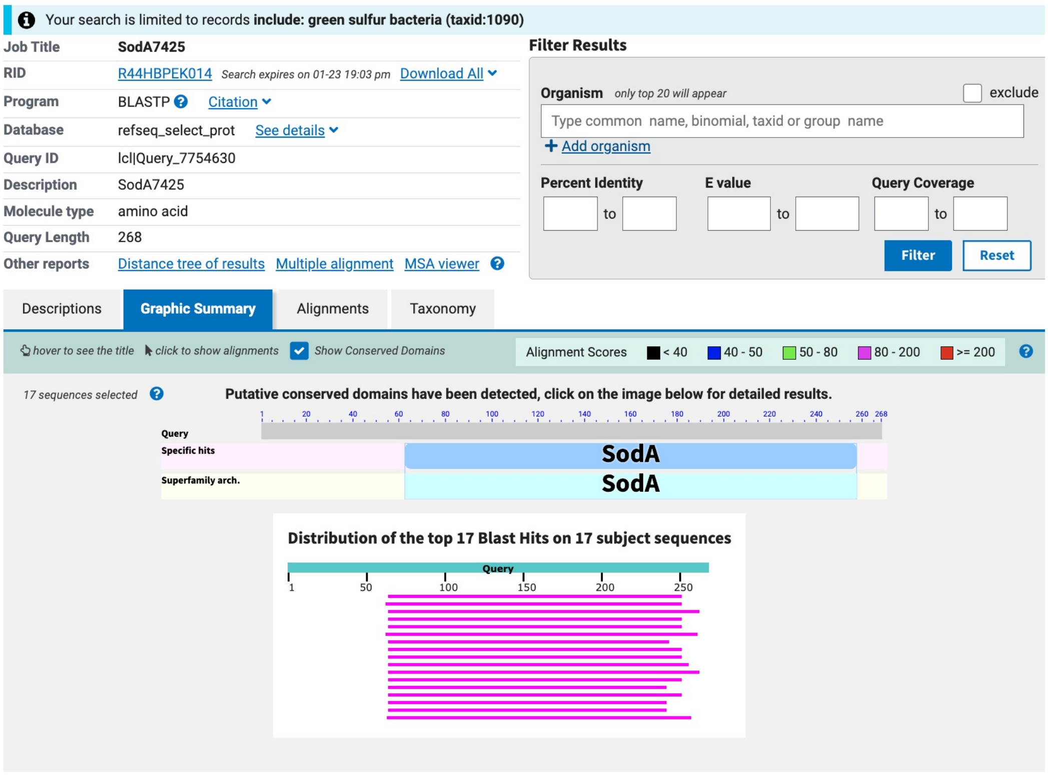

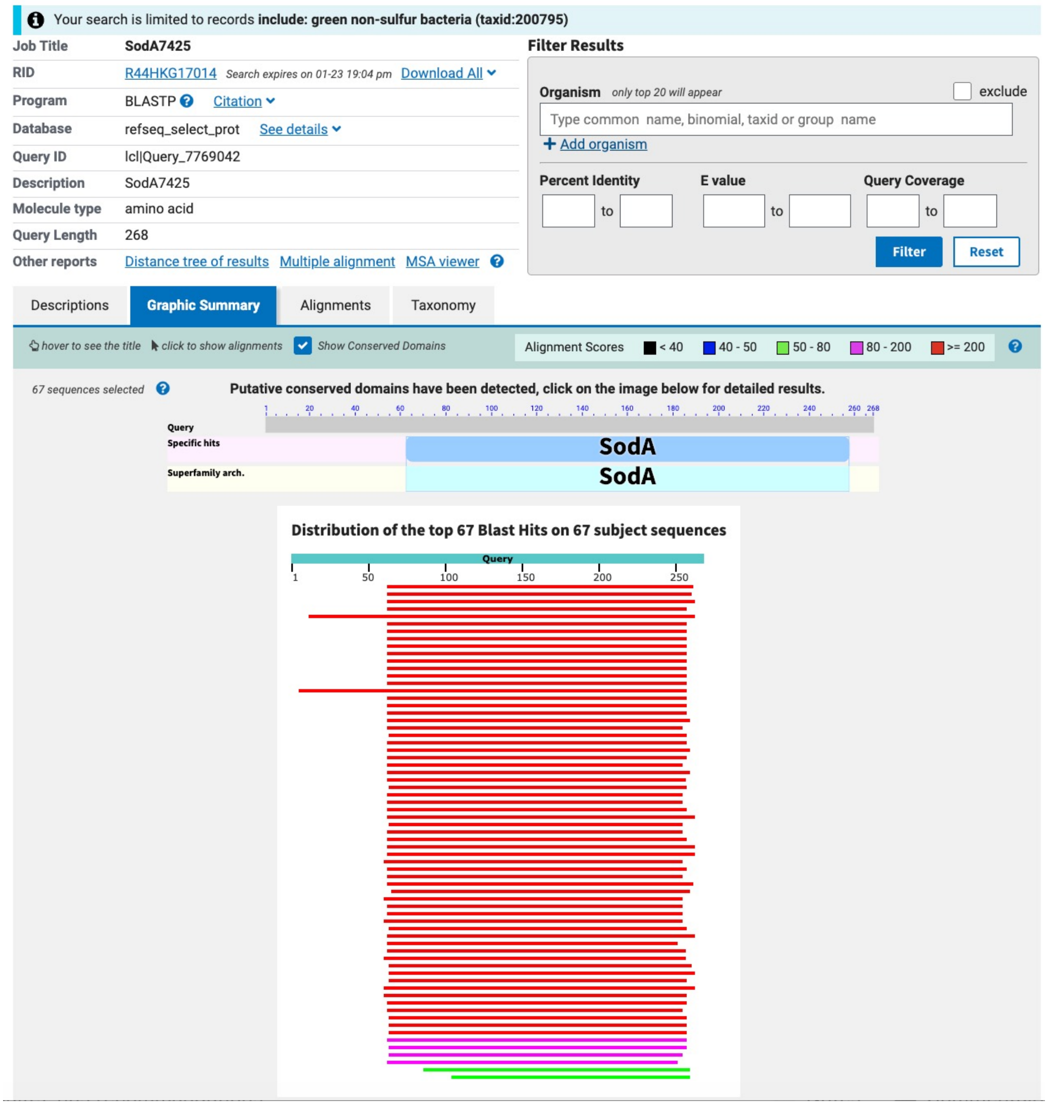

**Fig. S2:** BLASTP of *Cyanothece* PCC 7425 MnSodA against photosynthetic green sulfur and non-sulfur bacteria genomes

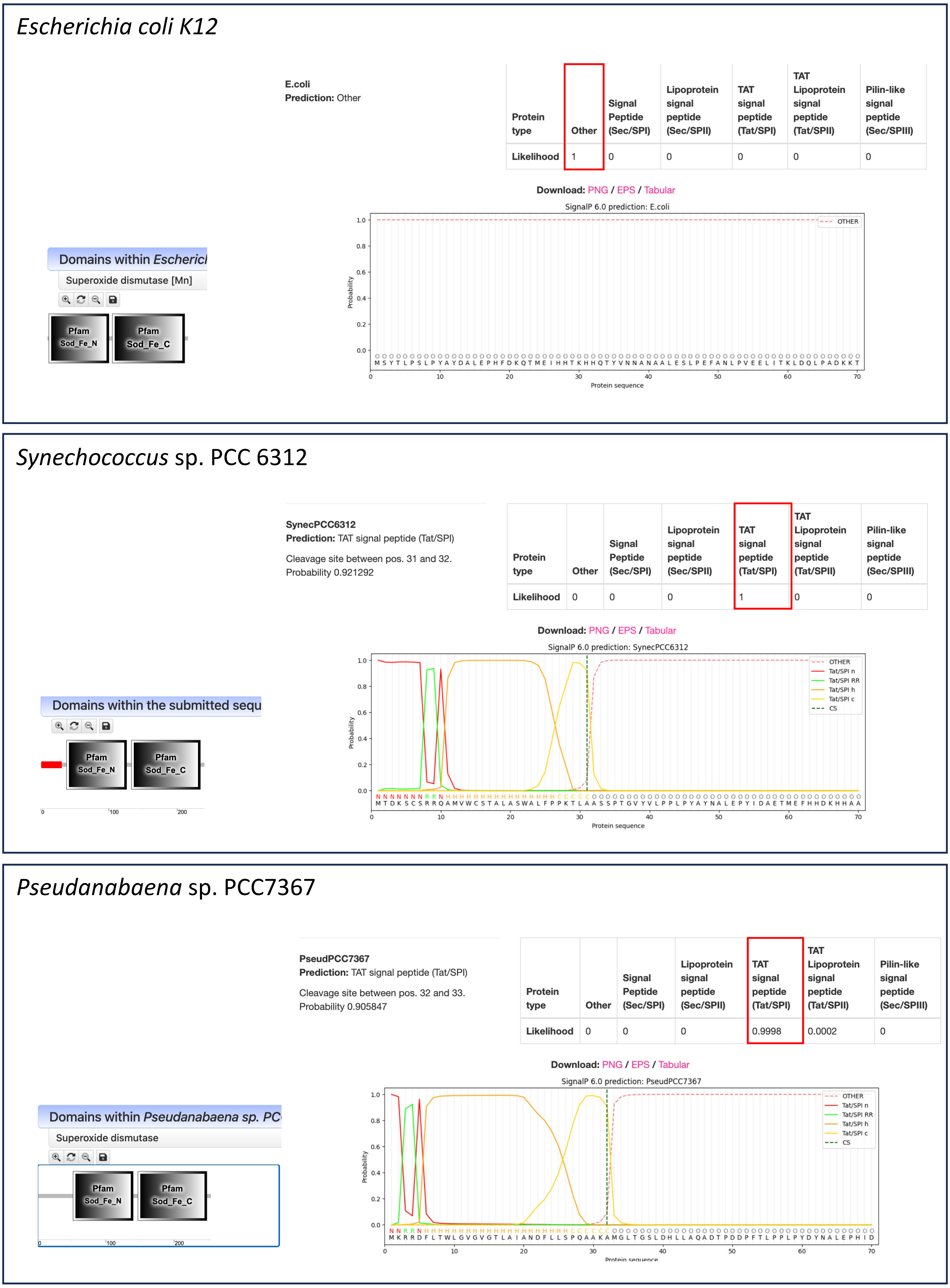

**Fig. S3**: SignalP 6.0 and Smart predicted profile of soluble MnSodA from *E. coli*, *Synechococcus* sp. PCC 6312 and *Pseudanabaena* sp. PCC7367.

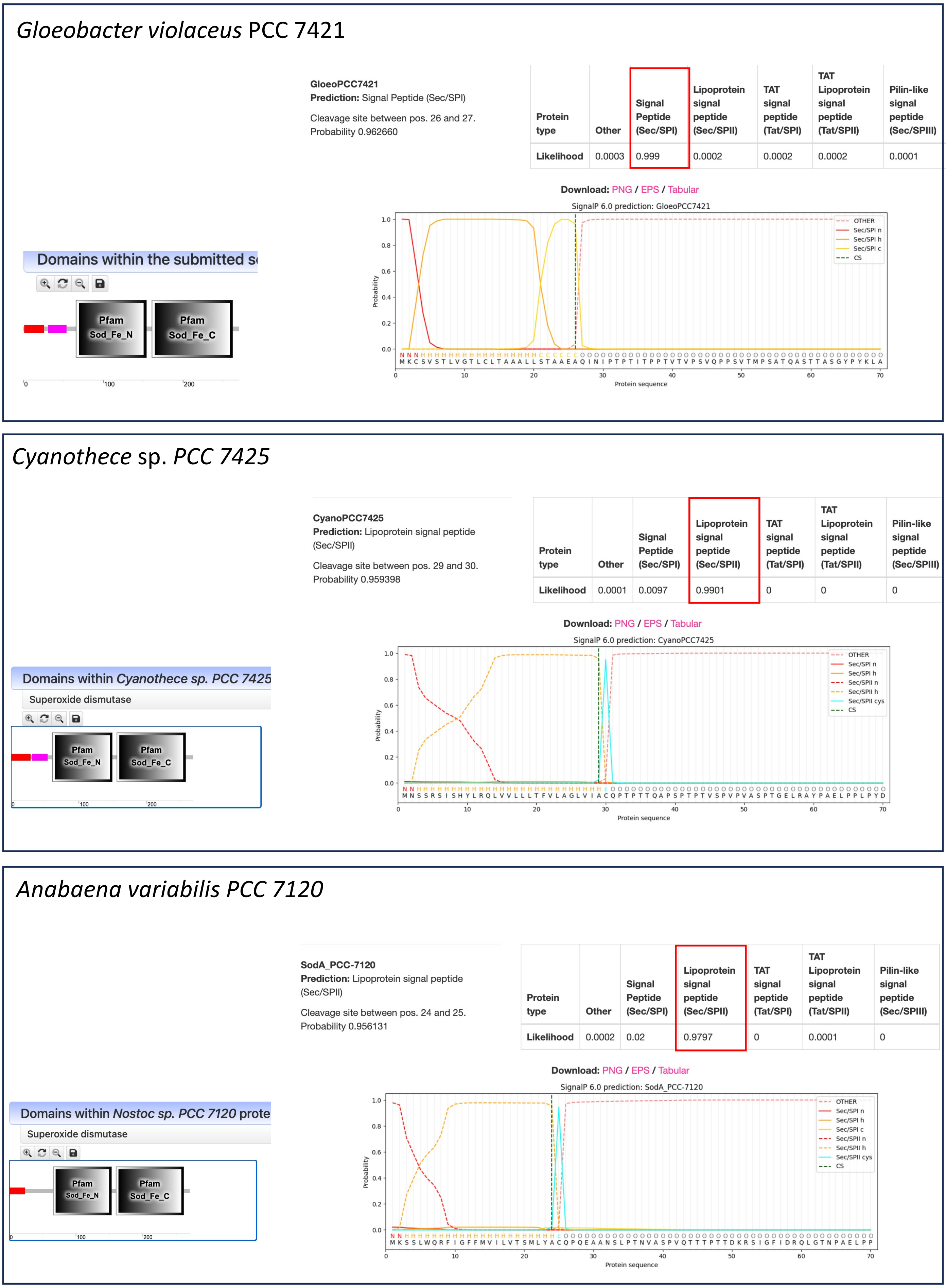

**Fig. S3**: SignalP 6.0 and Smart predicted profile of MnSodA from *Gloeobacter violaceus*, and Lipo-MnSodA from *Cyanothece* sp. *PCC 7425* and *Anabaena variabilis PCC 7120*

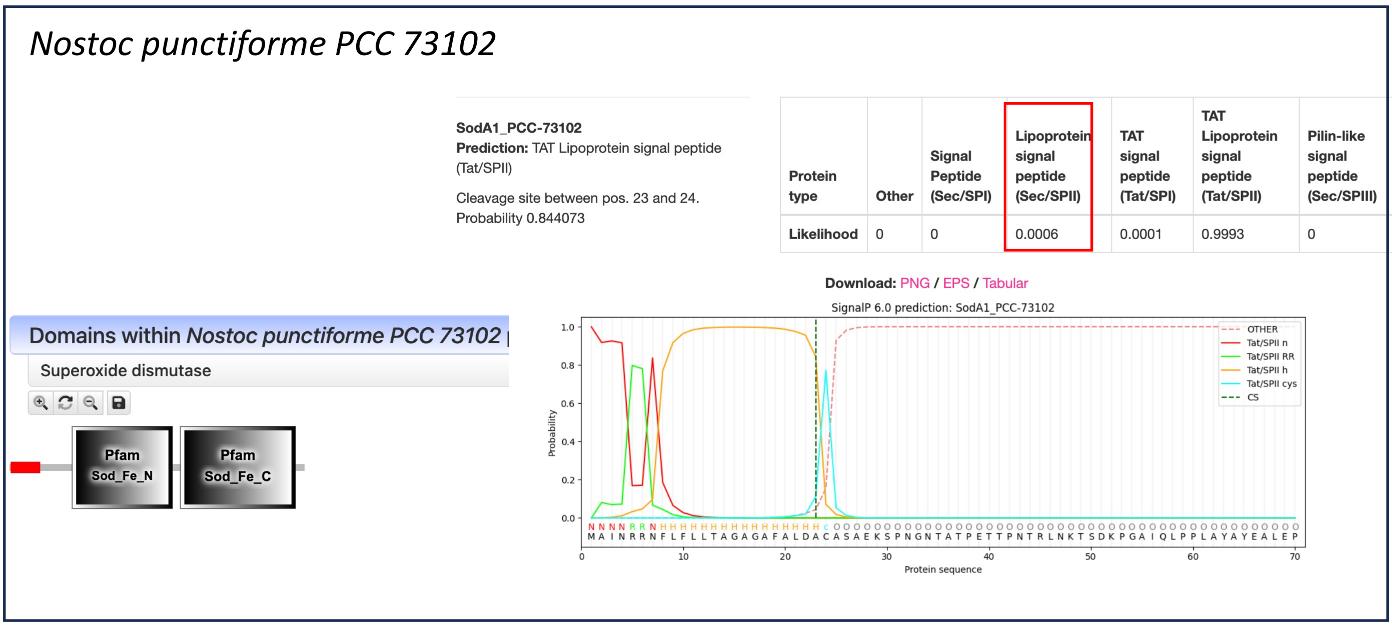

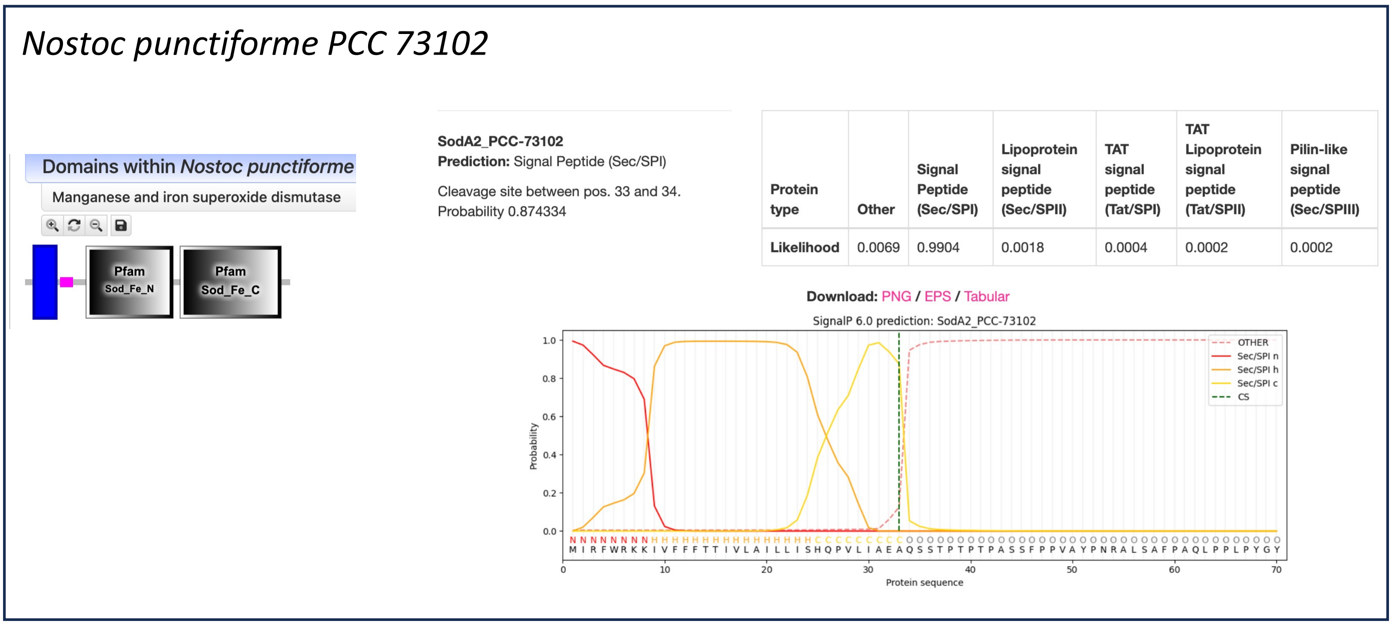

**Fig. S3**: SignalP 6.0 and Smart predicted profile of the two MnSodA from *Nostoc punctiforme* PCC 73102. MnSodA1 is predicted to be Lipo-anchored and MnSodA2 has a transmembrane helix.

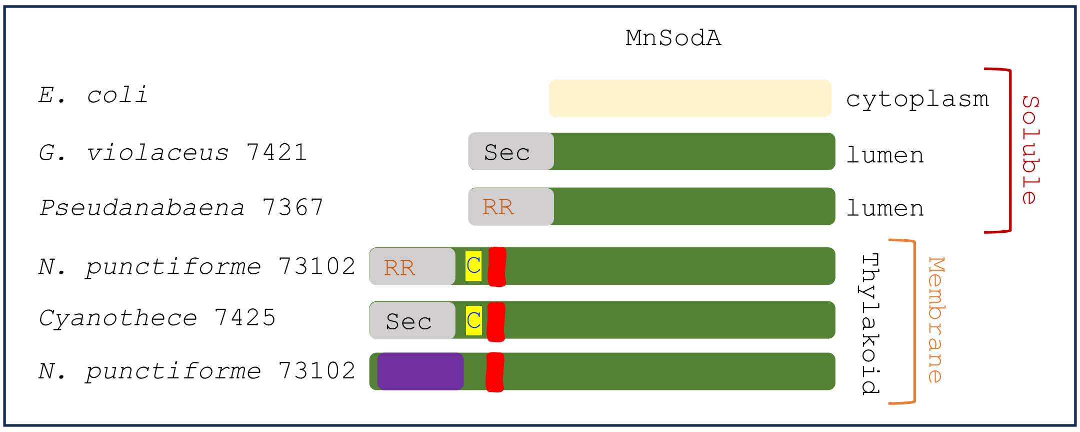

**Fig. S4:** Cartoon showing the different MnSodA forms and features. Grey: signal peptides, violet : transmebrane helix and red: Proline rich domain.

| | | |
|---|---|---|
| ***E. coli*** | ------------------------------------------------------------ | 0 |
| ***G. violaceus* 7421** | -------MKCS-------VSTLVGTLCLTAAALLSTAAEAQIN--IPTPTITPPTVTVPS | 44 |
| WP_015156211.1 | ----MRIRSLLK-VNKWFVVGFLLTIVLAIGQPL-LLA--QAQTSTA--ISQ---VSPVT | 47 |
| ***Synechococcus* 6312** | ----MTDKSCSRRQAMVWCSTALASWALFPPKT--------------------------- | 29 |
| WP_322876523.1 | ----MTDKPCSRRQALVWCGTALFSWALFPSKT--------------------------- | 29 |
| WP_277866136.1 | ------MVLLTRRRTLALLGITTLSWGLGSRPV--------------------------- | 27 |
| WP_011055891.1 | -------MRLTRRQTLGLLGVSAATLWLRSRPA--------------------------- | 26 |
| WP_297761391.1 | -------MRLTRRQTLSLLVGSAATLWLRSRPA--------------------------- | 26 |
| WP_024125507.1 | -------MRLTRRQTLSLLVGSAAALWLRSRPA--------------------------- | 26 |
| WP_365970883.1 | -------MRLTRRQTLSLLVGSAAALWLRSRPA--------------------------- | 26 |
| WP_409873764.1 | -------MLLTRRQALGLLVGSTTTLWLGSRPA--------------------------- | 26 |
| WP_300872640.1 | -------MPLTRRQTLGILVGSAATFWLRSRPA--------------------------- | 26 |
| WP_250682089.1 | -------MRLTRRQTLSLLVAGTATLWLRSRPA--------------------------- | 26 |
| WP_409869042.1 | -------MPLTRRQTLGLLVGSAATLWVRSRPA--------------------------- | 26 |
| WP_297053013.1 | -------MPLTRRQTLGLLVGSAATLWLRSRPA--------------------------- | 26 |
| WP_297051732.1 | -------MPLTRRQTLGLLVGSAATLWLRSRPA--------------------------- | 26 |
| WP_181494764.1 | -------MPLTRRQTLGLLVGSAATLWLRSRPA--------------------------- | 26 |
| WP_460193008.1 | -------MPLTRRQTLGLLVGSAATLWLRSRSA--------------------------- | 26 |
| WP_286627080.1 | -------MPLTRRQTLGLLVGSAATLWLRSRPA--------------------------- | 26 |
| WP_310743007.1 | -------MPLTRRQTLGLLVGSAATLWLRSRPA--------------------------- | 26 |
| WP_206200084.1 | -------MPLTRRQTLGLLVGSAATLWLRSRPA--------------------------- | 26 |
| WP_310821562.1 | -------MPLTRRQTLGLLVGSAATLWLRSRPA--------------------------- | 26 |
| WP_310800931.1 | -------MPLTRRQTLGLLVGSAATLWLRSRPA--------------------------- | 26 |
| WP_310454098.1 | -------MPLTRRQTLGLLVGSAATLWLRSRPA--------------------------- | 26 |
| WP_302677659.1 | -------MPLTRRQTLGLLVGSAATLWLRSRPA--------------------------- | 26 |
| ***Pseudanabaena* 7367** | ---------MKRRDFLTWLGVGVGTLAIANDFLLSPQAAKAMGL---------------- | 35 |
| MCU0526905.1 | -------MTFSRRHFLTLLGATTGTAAFSLVGLPETSPIAATQPA--------------- | 38 |
| MDJ0592545.1 | -------MSFNRRNFLIVLGAGVLATGLKTFPQL-------------------------- | 27 |
| MBX5479872.1 | ---------MNKRDFLRYALYAPAAVALWTRTSY-SSP---------------------- | 28 |
| WP_041973980.1 | ---------MNKREFLRYALYAPAAVALWTRTSY-SSP---------------------- | 28 |
| WP_428360810.1 | -------MTLNRRTFLYLLGASAGAAGLGLAAE----P---------------------- | 27 |
| WP_233744687.1 | -------MTLNRRTFLYLLGASAGAAGLGLAAE----P---------------------- | 27 |
| WP_445926464.1 | -------MTLNRRTFLYLLGASAGAAGLGLATE----P---------------------- | 27 |
| WP_015152615.1 | -------MSIKRRNFLFLVGGTAGAYAIETYTSHNLA----------------------- | 30 |
| WP_190455393.1 | -------MALNRRKFLFLLGATTGAVAIGAFPSACSAQSTQESNATQAP----------- | 42 |
| WP_190424133.1 | -------MALNRRKFLFLLGATTGAVAIGAFPSACSAQSTQESNATQAP----------- | 42 |
| WP_027843320.1 | -------MNLNRRHFLILFGTCAASYTLNACAFAENSPKTEAEEVT-------------- | 39 |
| WP_374791410.1 | -------MTVTRRHFLILLGASVSSITVGACLASGDNISRADNKST-------------- | 39 |
| WP_374825979.1 | -------MTVTRRHFLILLGASVSSITVGACLASGDNISRADNKST-------------- | 39 |
| WP_190465982.1 | -------MTVTRRHFLILLGASVSSITVGACLASGDNISTADNKST-------------- | 39 |
| BDA72795.1 | -------MILNRRHFLFLLGASVGSLTLEGCALDRNNPSPENTLTA-------------- | 39 |
| GJD16125.1 | -------MILNRRHFLFLLGASVGSLTLEGCALDRNNPSPENTLTA-------------- | 39 |
| MEC4881204.1 | -------MIISRRHFLYLLGASAGALVLDSCALAQNAPETTQAS---------------- | 37 |
| WP_190560307.1 | -------MSLNRRHFLVLLGASAGAFLLDGCASAETSP---------------------- | 31 |
| MFM5980434.1 | -------MSLNRRHFLVLLGASAGAFLLDSCASAETSP---------------------- | 31 |
| WP_096571931.1 | -------MSLNRRHFLVLLGASAGAFLLDSCASAETST---------------------- | 31 |
| WP_190648403.1 | -------MSLNRRHFLVLLGASAGAFLLDSCASAETST---------------------- | 31 |
| WP_190347369.1 | -------MSLNRRHFLVLLGASAGAFLLDSCASAETSP---------------------- | 31 |
| ***N. punctiforme* 73102** | -------MAINRRNFLFLLTAGAGAFALDACASAEKSPNGNTATPETTPNTR-------- | 45 |
| WP_334931430.1 | -------MTINRRNFLFLLTAGAGAFVFDACALAEKSPDGNAATLESTPNTK-------- | 45 |
| WP_335056299.1 | -------MTINRRNFLFLFTAGVGTFALDACASAETSPERNTAIPETTP----------- | 42 |
| WP_322673311.1 | -------MTINRRHFLFLLTSGAGAFALDACALAEKFPERNTAIPETTP----------- | 42 |
| WP_322689216.1 | -------MTINRRHFLFLLTSGAGAFALDACALAEKSPERNTAIPETTP----------- | 42 |
| WP_322701324.1 | -------MTINRRHFLFLLTSGAGAFALDACALAEKSPERNTAIPETTP----------- | 42 |

**Fig. S4:** Alignement of MnSodA from cynobacteria showing the different N-ter domains and conserved features. The twin arginine (RR), the signal peptide (in grey), the lipid attachement cysteine, the helix and proline domain are highlited.

```
WP_335012934.1        -------MTINRRHFLFLLTAGAGAFALDACALAEKSPNGNTTTPEAT-------------   41
WP_292868329.1        -------MTINRRHFLVLLTSGAGAFALDACASAEKSPQGNTATSNTTPNIT---------   45
WP_334926809.1        -------MTINRRHFLFLLTSGAGAFALDACASAEKSPQGNTATSETTPNIT---------   45
WP_109007648.1        -------MTINRRYFLFLLTSGAGAFALDACALAEKSPDGKTATPNTIANT----------   44
WP_289795776.1        -------MTINRRHFLFLLTAGAGAFALDACALAEKSPNENTATPN----T----------   40
WP_099100092.1        -------MTINRRHFLFLLTTGAGAFALDACALAENSPNGNTATPN----T----------   40
WP_335201774.1        -------MTINRRHFLFLLTSGAGAFALDACALAEKSPTGNTATPNTAANT----------   44
MDZ7959988.1          -------MAINRRHFLFLLGAGAGTFALEACALAEKSPSENPTSPEPKSD-----------   43
MBD2208974.1          -------MSMNRRHFLFLVGVGVGSFALEACALAENSPSENPTSPEPKSD-----------   43
MBD2339284.1          -------MSMNRRHFLFLVGVGVGSFALEACALAENSPSENPTSPEPKSD-----------   43
MBD2166265.1          -------MSMNRRHFLFLVGAGVGTFALEACALAENSPSENPTSPEPKSD-----------   43
WP_167724335.1        -------MSMNRRHFLFLVGAGVGTFALEACALAENSPSENPTSPEPKSD-----------   43
WP_045869977.1        -------MSIDRRHFLFLVGAGVGTFALEACALAEKSPSENPSSPEPKSD-----------   43
WP_096582728.1        -------MSINRRHFLFLVGAGVGTFALEACALAENSPSENPTSPEPKSD-----------   43
WP_096599871.1        -------MTINRRHFLFLLVAGTGTLALEGCASAEQSPTGNPTSPEPKPD-----------   43
WP_190743315.1        -------MTINRRHFLFLLAVGTGTLALEGCASAEQSPSGSQTSPEPKPD-----------   43
WP_190544005.1        -------MTINRRHFLFLLATGTGTLALQACASAEQSPSGSQTSPEPKPD-----------   43
WP_096643960.1        -------MTINRRHFLFLLATGTGTLALQACASAEQSPSGSQTSPEPKPD-----------   43
Cyanothece 7425       MNSSRSISHYLRQ-LVVLLLTFVLAGLVIACQP-TPTT--Q------APSPT------PT   44
WP_414578860.1        ------MFQFLQKTFAL----LLSAILLVACGT-MPIA--ESQSVNTT--TP------TI   39
WP_323267495.1        ------MFQFLQKTFAL----LLSAILLVACGT-MPIA--ESQSVNTT--EP------TI   39
WP_214431594.1        -------MKFVQKISVFLLAVVMVGMLLIACESSLPQA--DSQSPTANPNPT------PI   45
WP_322662701.1        ------MIRFVHKISAFLLTVIMVGMLLIACESSLPQA--DSQSPTANPNPT------PI   46
RCJ28434.1            ------MIRFVHKISAFLLTVIMVGMLLIACESSLPQA--DSQSPTANPNPT------PI   46
MBE9051467.1          ------MTKFLRDKIAFLLTGVMLAVLLISCQP-VPVA--ESQSPKAT--PT------TA   43
MBW4558831.1          ----------------------MLAVLLISCQP-VPVA--ESQSPKAT--PT------AA   27
WP_063871086.1        ------MSKFIREKIAFVLTGMLLAMMLISCQP-VPVA--ESQSPQGT--PI------AA   43
WP_006196478.1        ------MSKFIREKIAFVLTGMLLAMMLISCQP-VPVA--ESQSPQGT--PI------AA   43
WP_271918904.1        ------MSKFIREKIAFVLTGMLLAMMLISCQP-VPVA--ESQSPQGT--PI------AA   43
WP_271942563.1        ------MSKFIREKIAFVLTGMLLAMMLISCQP-VPVA--ESQSPQGT--PI------AA   43
WP_272068383.1        ------MSKFIREKIALVLTGVLVAVMLISCQV-IPIA--ESQSPQAT--PT------AV   43
WP_193998321.1        ------MSKLIREKIAFVLTGVLLAIVLISCQA-VPIA--ESQSPA-T--PT------AT   42
WP_089093562.1        ------MSKFIRKKIAFILTGVLLAIVLISCQT-VPVA--ESQSPQAT--PI------VI   43
WP_323215458.1        ------MSKFIREKIAFLFTGVLLAILLISCQA-VPVA--ESQSPQAT--PT------AT   43
WP_414550770.1        ------MSKFIREKIAFVLTGVLLAIVLISCQA-VPIA--ESQSPQPT--PT------AT   43
WP_214442418.1        -------MKRTFKKITAFFLTAVLGIVLISCQP-LPIA--GSQSPTAT--PV------PV   42
MEA5507287.1          -------MTNILQKITAFLFTSVLGILLISCQP-LPIA--GSQSPAAT--PT------SV   42
WP_281484005.1        -----MLITKIRQKVTAFLFTVVLGILLVSCQP-VPLA--GSQSPTVT--PT------PT   44
WP_162421941.1        ------MKTFLRHPIT-ALLGVVLSAIVVACQPTQP-V--GSQSPTASP--------SPA   42
WP_430678559.1        ------MFRVLKHPIT-VAVAVVLGFLVVACQPNET-A--VFQTATPTP--------SPG   42
MBE9012496.1          ------MFKLLKSPIV-LAITFVFSLLVVSCQPNET-A--VSQTPAAIP--------SPV   42
WP_059000726.1        ------MVRSLKRPIA-FATAFLLGCLMIACQPNET-A--VSQTPSPTP--------SPV   42
WP_482030890.1        ------MNRFLRQPIQFLILTIGLVMLLISCQQE-PQA--QQP-------------SP--   36
WP_445299659.1        ------MKNFLQQPIRFLIAGAAVVLLIVSCQST-PQA--QTSQSTP--QAQ---TSPST   46
WP_442927815.1        ------MKNFLQQPIRFLIAGAAVVLLIVSCQST-PQA--QTSQSTP--QAQ---TSPST   46
WP_445299397.1        ------MKNFLQQPIRFLIAGAAVVLLIISCQST-PQA--QTSQ-----------TSPST   40
WP_445312326.1        ------MKNFLQQPIRFLIAGAAVVLLIISCQST-PQA--QTSQ-----------TSPST   40
HIK16311.1            ------MKQLLHRSTRFLLAGVALLVLLISCATA-PQA--QTP-------------SPT-   37
WP_432812051.1        ------MKNHLTRSIRFLVAGAALLLLLISCQTS-PQA--QIP-------------APT-   37
WP_431655370.1        ------MKKFLQRPIRFLVTGMALVVLLIACQTS-PQA--QSP-------------APTT   38
MGI0483888.1          ---------------------MALVVLLIACQTS-PQA--QSP-------------APTT   23
MBW4417879.1          -------MKRLLKHSLKLLLGLVMVVMFAACQPT-PTA--NSQSP------------APA   38
MBW4540880.1          -------MKRLLKQSLKLLLGVAMVVMFAACQPT-PTA--NSQSP------------APT   38
MBW4644872.1          -----MSIKFLRQQFLLLITGVFLVFVLLACQRA-PIA--ESPTPTATPVTER----TIA   48
PMB52031.1            -----MIIKFLRQQFVLFVTGIFLVSVLLACQGT-PIA--ESPTPTVTPATE---QTLAA   49
WP_190751173.1        -----MLTKFLRQQFVLFIAGIFLVSVLFACQEN-QIT--ESPTATVSPVTQ---QTPLT   49
WP_190549236.1        -----MLTKFLRQQFVLFIAGIFLVSVLFACQEN-QIT--ESPTATVSPVTQ---QTPLT   49
WP_190629649.1        -----MLTKFLRQQFVLFIAGIFLVSVLFACQEN-QIT--ESPTATVSPVTQ---QTPLT   49
MBD2168452.1          -----MTAKFWRQQFVLFITGIFLVSVLFACQGT-PIA--ESPNATTTPVTE---QNLAA   49
MBD2342527.1          -----MIAKFWRQQSALFITGIFLVSVLFACQGT-PIA--ESPNATATPVTE---QNLAA   49
MHN9116841.1          -----MIVKFLRQQIVLLITGFFLVSLLYACQGE-PIA--QSPTPAVTPTATP--AAQTT   50
MFN6514993.1          -----MLIKFLRQQFVLFITGVFLVSALFACQGT-PIA--DSPTPTATPATQ---ETFGT   49
NDJ23487.1            -----MSIKFLRQKLVLFITGAFLVSLLFACQGT-PIV--ESPTPTATPITQ---DTIAT   49
WP_181929818.1        -----MIIKFLRQQVVLFITGAFLVSLLYACQGE-PIA--ESPTPPATPAAQ---DTVVA   49
MBV6623186.1          ----MIVKKTWRQKSILFLTGFFLISLLFACQTA-SIN--ESSTSTATPVTEK----TFS   49
WP_323238745.1        ---MMIIRKFWRQKSILFLTGFFLISLLFACQRV-PIV--ESQTSTATPVAEE----TAT   50
WP_254449390.1        -----MIMKFLQQQVMLFITGVCLVSLLFACQVE-PIT--ELSTSITTPVQDV----VAT   48
MBF2015390.1          -----MLVKFWQQKFGFFLTGFCLIFLLFACQGE-TTT--ESPISTATPATDE----TAT   48
MEB3216192.1          -----MTVKFWRQKFVFFLTGFCLIFLLFACQGE-TIT--QSPSPTATPATDE----TAT   48
WP_193917416.1        -----MTVKIWRQKFVFFLTGFCLIFLLFACQGE-TIT--QSPSPIATPATDE----TAT   48
WP_277875608.1        ------MKSSRWQQFICFCMGIFLISTLYACQPQE-VA----ESSTS-PVATPVQKTAPT   48
MBD2346042.1          -------------------MGIFLISTLYACQPQE-VA----ESSTS-PVATPVQKTAPT   35
WP_010994247.1        ------MKSSLWQRFIGFFMVILVTSMLYACQPQE-AA----NSLPT-NVASPVQTTTPT   48
Nostoc sp. PCC 7120   ------MKSSLWQRFIGFFMVILVTSMLYACQPQE-AA----NSLPT-NVASPVQTTTPT   48
WP_277884298.1        ------MKSSLRQRFIGFFMIILVTSMLYACEPQE-AA----NSLPT-NVATPVQTTAPT   48
WP_414756233.1        ------MKSSLWQRFIGFFMVVLVTSMLYACQPQE-AA----NSLPT-NVASPVQTTAPT   48
WP_011318269.1        ------MKSSLWQRFIGFFMVILVTSMLYACQPQE-AA----NSLPT-NVASPVQTTATA   48
```

```
WP_015215174.1        ------MKTLLLQRVIGFCTGILLVFTLFACQPQQ-VA----ESLET-PIAVPTQTTTLV    48
WP_276752922.1        ------MKSFFLRPIIWFFTGILLVSTLFACQPQQ-VA----ETPTTNTTATPIQTTAPV    49
RUR73923.1            -----MKSFFLRPIIWFFTGILLVSTLFACQPQQ-VA----ETPTTNTTATPIQTTAPV    50
WP_016872332.1        ------MKSFFLRPIIWFFTGILLVSTLFACQPQQ-VA----ETPTTNTTATPIQTTAPV    49
WP_341997750.1        -----MKKSFVLQPIIWFLTGILLVSTLFACQPQQ-VA----ESPT-NTTATPIQTTAPV    49
WP_289789894.1        ------MKSFVLQPIIWFFTGILLVSTLFACQPQQ-VA----ESPT-NTTATPIQTTVPI    48
WP_191756852.1        ------MKSFFWQPVVWFFTVILLVSTLFACQSQQ-VT----ESPSS-TTATPIQTTAPV    48
WP_193198009.1        ------MKSFFWQPVVWFFTVILLVSTLFACQPQQ-VT----ESPST-TPATPVQTTAPV    48
WP_323228814.1        ------MKSYFRQPVILFCVGFFLISTLFACQPQQ-VA----S-PPT-STATPVQTTAPV    47
WP_103138074.1        ------MKKFFKQPVMWFCTTILLVSTLFACQTQQ-AA----ELSET-PLANPVQTSAPV    48
WP_242072232.1        ------MKLFLKQPVIWFCMGMLLISTLFACQPQQ-VA----ESPPI-PTATPVQTSAPA    48
MBD2440923.1          ---------------------MLLISTLFACQPQQ-VA----ESPPI-PTATPVQTSAPA    33
WP_318780343.1        ------MMNFLRQKISYLLMGVMLFMVLGACQP-VPTV--ESQSVTGTPIATSVATQTPA    51
WP_460206421.1        ------MMKFLRQQISYLIIGIALFALLASCQSTTPTA--ESPSPTETPAATSVTTEAQT    52
WP_169219177.1        -------------------MGIALFILLASCQL-TPVA--ESPSPTETPTAIPVVTQTP-    37
WP_332997608.1        ------MRKFLRQQISYLLMGTAVFILLASCQP-TPVA--ESPSPAETSTATPVVTQTP-    50
MBW4498943.1          -------------------MGTALFILLASCQP-TPVA--ESPSPAETATATPVVTQTP-    37
WP_414585734.1        ------MRKFLRQRISYLLMGTALFILLASCQP-TPVA--ESPSPAETPTATPVVTQTP-    50
MCC5641606.1          ------MIGFRRKTIVFFLATIVLAILVIFYEP-VLIA--EAQSSTP----T------AA    41
WP_435020268.1        ------MIKFWRKTIVFFFTTILLAILLISHEP-VPIA--EAQSSTPI--PT------PA    43
WP_292704904.1        ------MLRFWRKKIVFFFTTIVLAILLISHQP-VLIA--EAQSSRPT--PT------PV    43
WP_335036612.1        ------MIGFLRKKIVFFFTIIVLAILLISHQP-VLIA--EAQSSTPT--PT------PA    43
WP_302480924.1        ------MIRFSRKKIVFFFTTIVLAILAISHQP-VLIA--EAQSSTPT--PT------PA    43
WP_311774093.1        ------MIRFLRKKIVFFFTTIVLAILLISHQP-VLIA--EAQSSKPT--PT------SA    43
N. punctiforme 73102------MIRFWRKKIVFFFTTIVLAILLISHQP-VLIA--EAQSSTPT--PT------PA    43
WP_322745234.1        ------MIRFWRKKIVFFFTTIVLAILLISHQP-VLIA--EAQSSTPT--PK------PA    43
WP_306297076.1        ------MIRFLRKKIVFFFTTIVLAILLISHQP-VLIA--EAQSSTPT--PT------PA    43
WP_322746338.1        ------MIRFLRKKIVFFFTTIVLAILLISHQP-VLIA--EAQSSTPT--PT------PA    43
WP_277883559.1        ------MIRFLRKKIVFFFTTIVLAILLISHQP-VLIA--EAQSSTPT--PT------PA    43
WP_392482408.1        ------MIRFLRKKIVFFFTTIVLAILLISHQP-VLIA--EAQSSTPT--PT------PA    43
WP_163931552.1        ------MIRFLRKKIVFFFTTIVLAILLISHQP-VLIA--EAQSSTPT--PT------PA    43
WP_114081005.1        ------MIRFLRKKIVFFFTTIVLAILLISHQP-VLIA--EAQSSTPT--SM------PA    43

E. coli               --------------------MSYTLPSLPYAYDALEPHFDKQTMEIHHTKHHQTYVNNAN    40
G. violaceus 7421     VQPPSVTMPSATQASTTASGYPYKLAPLPYDYSALEPYIDAETMKLHHDKHHQAYVDNLN
     104
WP_015156211.1        QQS-----RPVASPKEELSANPAQLPPLPYEYNALEKAIDAKTMKLHHDEHHATYVEELN
     102
Synechococcus 6312    -------------LAASSPTGVYVLPPLPYAYNALEPYIDAETMEFHHDKHHAAYVNNLN    76
WP_322876523.1        -------------LAASSPTGVYVLPPLPYAYNALEPYIDAETMQFHHDKHHAAYVNNLN    76
WP_277866136.1        --------------AAEPASGSYELPPLPYDYDALEPYIDGETMRFHHDKHHAGYVKKLN    73
WP_011055891.1        -------------VAAGEPSEIYTLPPLPYDYDALEPVIDAETMRLHHDKHHAAYVNTLN    73
WP_297761391.1        -------------VAADEPSEIYTLPPLPYDYDALEPVIDAETMRLHHDKHHAAYVNTLN    73
WP_024125507.1        -------------VAAGEPSEIYTLPPLPYDYEALEPVIDAETMRLHHDKHHAAYVNTLN    73
WP_365970883.1        -------------VAAGEPSEIYTLPPLPYDYEALEPVIDAETMRLHHDKHHAAYVNTLN    73
WP_409873764.1        -------------IAVGEPSEIYTLPPLPYDYDALEPVIDAETMHLHHDKHHAAYVKTLN    73
WP_300872640.1        -------------VDEGEPLEIYKLPPLPYNYDALEPVIDAETMHLHHDKHHGAYVKTLN    73
WP_250682089.1        -------------VAEGEPLEIYTLPPLPYGYDALEPVIDAETMQLHHDKHHAAYVNTLN    73
WP_409869042.1        -------------RAEGELSEIYRLPPLPYAYDALEPVIDAETMHLHHDKHHAAYVKTLN    73
WP_297053013.1        -------------VAEGEPSEIYTLPPLPYGYDALEPVIDAETMQLHHNKHHGAYINTLN    73
WP_297051732.1        -------------VAEGEPSEIYTLPPLPYGYDALEPVIDAETMQLHHNKHHGAYINTLN    73
WP_181494764.1        -------------VAEGDPSDIYTLPPLPYDYDALEPVIDAETMHLHHDKHHAAYVKTLN    73
WP_460193008.1        -------------MAEGEPSEIYTLPPLPYGYDALEPVIDAETMQLHHDKHHAAYVKTLN    73
WP_286627080.1        -------------VAEGEPSEIYTLPPLPYGYDALEPVIDAQTMQLHHDKHHAAYVKTLN    73
WP_310743007.1        -------------VAEGEPSEIYTLPPLPYGYDALEPVIDAETMQLHHDKHHAAYVKTLN    73
WP_206200084.1        -------------VAEGEPSEIYTLPPLPYGYDALEPVIDAETMQLHHDKHHAAYVKTLN    73
WP_310821562.1        -------------VAEGEPSEIYTLPPLPYGYDALEPVIDAETMQLHHDKHHAAYVKTLN    73
WP_310800931.1        -------------VAESEPSEIYTLPPLPYGYDALEPVIDAETMQLHHDKHHAAYVKTLN    73
WP_310454098.1        -------------VAESEPSEIYTLPPLPYGYDALEPVIDAETMQLHHDKHHAAYVKTLN    73
WP_302677659.1        -------------VAEGEPSEIYTLPPLPYGYDALEPVIDAETMQLHHDKHHAAYVKTLN    73
Pseudanabaena 7367    -----TGSLDHLLAQADTPDDPFTLPPLPYDYNALEPHIDVRTMQIHHDRHHAGYVRNLN    90
MCU0526905.1          -------AAARLWDRTAQASETFKLPPLPYAYDALEPHIDAETMQFHHDKHHAAYVKNLN    91
MDJ0592545.1          ------------ALAEDEPQGPFELPPLPYDYNALEPYIDAETMMFHHDKHHAGYTKNLN    75
MBX5479872.1          --------ESVISVSQQTASGPHKLPPLPYEYNALEPYIDTETMQLHHDKHHAAYVNNLN    80
WP_041973980.1        --------AAVISVSQQTASGPHKLPPLPYEYNALEPYIDTETMQLHHDKHHAAYVNNLN    80
WP_428360810.1        ------------TWAAESPKGPFTLPPLPYAYDALEPHIDKATMQFHHDKHHAAYVNNLN    75
WP_233744687.1        ------------TWAAESPKGPFTLPPLPYAYDALEPHIDKATMQFHHDKHHAAYVNNLN    75
WP_445926464.1        ------------TWAAESPKGPFTLPPLPYAYDALEPHIDKATMQFHHDKHHAAYVNNLN    75
WP_015152615.1        ------------WAQTPSASGDFKLPPLPYDYKALEPHIDAQTMRIHHDRHHATYVKNLN    78
WP_190455393.1        ------------ASPTAQTKDTFTLPPLPYDYKALEPHIDAATMQFHHDKHHAAYVKNLN    90
WP_190424133.1        ------------ASPTAQTKDTFTLPPLPYDYKALEPHIDAATMQFHHDKHHAAYVKNLN    90
WP_027843320.1        ------------KVASKPESGVIKLPPLPYAYDALEPHIDAETMRFHHDKHHAGYVKNLN    87
WP_374791410.1        ---------SGSTTNVAQNTGAFELPPLPYAYNALEPYIDARTMEFHHDKHHAAYVTNLN    90
WP_374825979.1        ---------SGSTTNVAQNTGAFELPPLPYAYNALEPYIDARTMEFHHDKHHAAYVTNLN    90
WP_190465982.1        ---------PGSTTNVAQNTGTFELPPLPYAYNALEPYIDARTMEFHHDKHHAAYVKNLN    90
BDA72795.1            -----APSPT---EKSEAPSYAIVLPPLPYGYDALEPHIDAETMRFHHDKHHATYVTNLN    91
GJD16125.1            -----APSPT---EKSEAPSYAIVLPPLPYGYDALEPHIDAETMRFHHDKHHATYVNNLN    91
MEC4881204.1          ----------------PGKTGTIELPPLPYAYDALEPHIDARTMEFHHDKHHAAYVKNLN    81
```

```
WP_190560307.1        ----------------PRTSAIELPPLPYAYDALEPHIDARTMQFHHDKHHAAYVNNLN   74
MFM5980434.1          ----------------PRTSTIELPPLPYAYDALEPHIDARTMQFHHDKHHAAYVNNLN   74
WP_096571931.1        ----------------PRTSTIELPPLPYAYDALEPHIDARTMQFHHDKHHAAYVNNLN   74
WP_190648403.1        ----------------PRTSTIELPPLPYAYDALEPHIDARTMQFHHDKHHAAYVNNLN   74
WP_190347369.1        ----------------PRTSTIELPPLPYAYDALEPHIDARTMQFHHDKHHAAYVNNLN   74
N. punctiforme 73102  -----------LNKTSDKPGAIQLPPLAYAYEALEPHIDAKTMQFHHDKHHATYVKNLN   93
WP_334931430.1        -----------SNTTQNKTGAIQLPPLPYAYEALEPHIDAKTMQFHHDKHHATYVKNLN   93
WP_335056299.1        --------NTIANTTPNTTGAIQLPPLPYAYEALEPHIDAKTMQFHHDKHHATYVKNLN   93
WP_322673311.1        --------NTIANTTPNKPGTIQLPPLPYAYDALEPHIDAKTMQFHHDKHHATYVKNLN   93
WP_322689216.1        --------NTIANTTPNKPGTIQLPPLPYAYEALEPHIDAKTMQFHHDKHHATYVKNLN   93
WP_322701324.1        --------NTIANTTPNKPGTIQLPPLPYAYEALEPHIDAKTMQFHHDKHHATYVKNLN   93
WP_335012934.1        -----------PSTTPNTTGAIQLPPLAYAYEALEPHIDAKTMQFHHDKHHATYVKNLN   89
WP_292868329.1        ---ANTTPNTIAKTTPNTTGAIQLPPLPYAYEALEPHIDAKTMQFHHDKHHATYVKNLN   101
WP_334926809.1        ---ANTTPNTIANTTPNTTGAIQLPPLPYAYEALEPHIDAKTMQFHHDKHHATYVKNLN   101
WP_109007648.1        ------TP--------NTTGAIQLPPLPYAYEALEPHIDAKTMQFHHDKHHATYVKNLN   89
WP_289795776.1        ------TP----NITANTTGAIQLPPLPYAYEALEPHIDAKTMQFHHDKHHATYVKNLN   89
WP_099100092.1        ------TP----NITANTTGAIQLPPLPYAYEALEPHIDAKTMQFHHDKHHATYVKNLN   89
WP_335201774.1        ------TP----NITANTTGAIQLPPLPYAYEALEPHIDAKTMQFHHDKHHATYVKNLN   93
MDZ7959988.1          ----------------TTKGAIALPPLPYAYDALQPHIDAKTMQFHHDKHHAAYVKNLN   86
MBD2208974.1          -----------------TKGAIQLPPLPYAYDALEPHIDAKTMQFHHDKHHATYVKNLN   85
MBD2339284.1          -----------------TKGAIQLPPLPYAYDALEPHIDAKTMQFHHDKHHATYVKNLN   85
MBD2166265.1          -----------------AKGAIQLPPLPYAYDALEPHIDARTMQFHHDKHHAAYVKNLN   85
WP_167724335.1        -----------------AKGAIQLPPLPYAYDALEPHIDAKTMQFHHDKHHATYVKNLN   85
WP_045869977.1        ----------------TTKGAIQLPPLPYAYDALEPHIDAKTMQFHHDKHHAAYVKNLN   86
WP_096582728.1        -----------------TKGAIQLPPLPYAYDALEPHIDAKTMQFHHDKHHATYVKNLN   85
WP_096599871.1        ----------------TTKGAIQLPPLPYAYEALEPHIDAKTMQFHHDKHHATYVKNLN   86
WP_190743315.1        ----------------TTTGAIQLPPLPYAYDALEPHIDARTMQFHHDKHHAAYVKNLN   86
WP_190544005.1        ----------------T-TGAIQLPPLPYAYDALEPHIDAKTMQFHHDKHHAAYVKNLN   85
WP_096643960.1        ----------------T-TGAIQLPPLPYAYDALEPHIDARTMQFHHDKHHAAYVKNLN   85
Cyanothece 7425       VSP-----VPVASPTGELRAYPAELPPLPYDGALETAIDARTMEIHHDRHHAAYVKNLN   99
WP_414578860.1        KSS-----LPVAYPDKELSASPAKLPPLPYDYTALSQAIDAETMKLHHDAHHASYVNNLN   94
WP_323267495.1        KSS-----LPVAYPDKELSASPAKLPPLPYDYTALSQAIDAETMKLHHDAHHASYVNNLN   94
WP_214431594.1        TSL-----PPVASPNGELSAYPAKLPPLPYDYGALEKAIDAETMKLHHDAHHATYVKNLN   100
WP_322662701.1        TAL-----PPVASPDGELSAYPAKLPPLPYDYGALEKAIDAETMKLHHDAHHASYVNNLN   101
RCJ28434.1            TAL-----PPVASPDGELSAYPAKLPPLPYDYGALEKAIDAETMKLHHDAHHASYVNNLN   101
MBE9051467.1          -TF-----PPVASPDGELSAAPAKLPPLPYDYTALGKAIDAETMKLHHDGHHASYVNNLN   97
MBW4558831.1          -TF-----PPVASPDGELSAAPAKLPPLPYDYTALGKAIDAETMKLHHDGHHAGYVNNLN   81
WP_063871086.1        -TF-----PPVASPDGELSASPAKLPLLPYDYAALGKAIDAETMKLHHDGHHASYVNNLN   97
WP_006196478.1        -TF-----PPVASPDGELSASPAKLPLLPYDYAALGKAIDAETMKLHHDGHHASYVNNLN   97
WP_271918904.1        -TF-----PPVASPDGELSASPAKLPLLPYDYAALGKAIDAETMKLHHDGHHASYVNNLN   97
WP_271942563.1        -TF-----PPVASPDGELSASPAKLPLLPYDYAALGKAIDAETMKLHHDGHHASYVNNLN   97
WP_272068383.1        -KF-----PPVASPDGELSASPAQLPLLPYDYAALGKAIDAETMKLHHDGHHASYVNNLN   97
WP_193998321.1        -TF-----PPVASPDGELSANPAKLPALPYDYTALGKAIDAETMKLHHDGHHASYVNNLN   96
WP_089093562.1        -TF-----PPVASPDGELSASPAQLPALPYDYAALGKAIDAETMKLHHDGHHASYVNNLN   97
WP_323215458.1        -TF-----PPVASPDGELSASPAKLPILPYDYAALGKAIDAETMKLHHDGHHASYVNNLN   97
WP_414550770.1        -TF-----PPVASPDGELSASPAKLPLLPYDYSALGKAIDAETMKLHHDGHHASYVNNLN   97
WP_214442418.1        RSR-----PPVVSPQGELSANPAKLPPLPYDYGVLGKAIDAETMRLHHDAHHASYVNNLN   97
MEA5507287.1          GSR-----PPVASAQGELSANPAKLPPLPYDYGVLGKAIDAETMKLHHDAHHASYVKNLN   97
WP_281484005.1        ESR-----PPVASTQNELSANPAKLPPLPYDYGVLGKAIDAETMKLHHDAHHASYVSKLN   99
WP_162421941.1        VTT-----LPVAYPDRELSASPAQLPPLPYAANALNQAIDAQTMQIHHDRHHQTYVDNIN   97
WP_430678559.1        -TV-----RPVAYVDRTLSASPAQLPPLPYPADALSKAIDAKTMQIHHDRHHKSYVDNLN   96
MBE9012496.1          -AA-----RPVAYVDRTLSASPAQLPPLPYPANALSKAIDTRTMQIHHDRHHKSYVDNLN   96
WP_059000726.1        -AD-----RPVAYVDRTLSASPAQLPPLPYPANALSKAIDAETMQLHHDRHHKSYVDNLN   96
WP_482030890.1        --------PVAQPDSELSANPAQLPPLPYEYGALTKAIDAETMKLHHDNHHAAYVENLN   87
WP_445299659.1        TQP-----SAVASPGGELSAYPAQLPPLPYEYGALAKAIDAETMKLHHDRHHASYVENLN   101
WP_442927815.1        TQP-----SAVASPGGELSAYPAQLPPLPYEYGALAKAIDTETMKLHHDRHHASYVENLN   101
WP_445299397.1        TQP-----SAVASPGGELSAYPAQLPPLPYEYGALAKAIDAETMKLHHDRHHASYVENLN   95
WP_445312326.1        TQP-----SAVASRAGELSAYPAQLPPLPYEYGALAKAIDAETMKLHHDRHHASYVDNLN   95
HIK16311.1            AQP-----AAVASPGGELSAYPAQLPPLPYEYGALAKAIDAETMKLHHDRHHASYVNNLN   92
WP_432812051.1        IQP-----TAVASSEGELSANPAQLPPLPYDYGALATAIDAETMKLHHDRHHAAYVENLN   92
WP_431655370.1        AQP-----AAVASPEGELSANPAQLPPLPYEYGALAQAIDAETMKLHHDRHHATYVENLN   93
MGI0483888.1          AQP-----AAVASPEGELSANPAQLPPLPYEYGALAQAIDAETMKLHHDRHHATYVENLN   78
MBW4417879.1          SPD-----VSVAYPARALSASPAQLPPLPYGYDALSKAIDAETMRLHHDQHHAAYVKNLN   93
MBW4540880.1          SPD-----ISVAYPDRALSASPAQLPPLPYGYDALSKAIDVETMRLHHDQHHAAYVKNLN   93
MBW4644872.1          NNQ-----LPVAYPDKELSASPAQLSELPYADNALEKAIDAETMKLHHDRHHAAYVNNLN   103
PMB52031.1            NQF-----PPVAYPERELSASPAELPELPYAYNALEKAIDAETMKLHHDRHHATYVENLN   104
WP_190751173.1        TNS-----PPVAYPDRELTTSPAQLPKLPYAYNALEKAIDAETMKLHHDRHHATYVNNLN   104
WP_190549236.1        TNS-----PPVAYPDRELTATPAQLPKLPYAYNALEKAIDAETMKLHHDRHHATYVNNLN   104
WP_190629649.1        TNS-----PPVAYPDRELTATPAQLPKLPYAYNALEKAIDAETMKLHHDRHHATYVNNLN   104
MBD2168452.1          SNF-----PPVAYPDKELTATPAQLPKLPYAYNALEKAIDAETMKLHHDRHHATYVKNLN   104
MBD2342527.1          SNF-----PPVAYPDKELSATPAQLPKLPYAYNALEKAIDAETMKLHHDRHHATYVENLN   104
MHN9116841.1          DQF-----PPVAYLDRELSTSPAQLPKLPYAYNALEKAIDAETMKLHHDRHHAAYVKNLN   105
MFN6514993.1          AQL-----TPAAYPDRELSASPAQLPKLPYANNALEKAIDAETMKLHHDRHHATYVENLN   104
NDJ23487.1            NQF-----PPVAYPDRELSASPAQLPKLPYAYNALEKAIDAETMKLHHDRHHATYVENLN   104
WP_181929818.1        TQF-----PPVAYPDRELSASPAQLPKLLYAYDALQKAIDAETMKLHHDRHHATYVENLN   104
MBV6623186.1          TQS-----FAVAYPDRELTTSPARLPELPYAANALEKAIDAETMKLHHDRHHAAYVKNLN   104
WP_323238745.1        TQL-----TAVAYPDKELSASPAQLPKLPYAYNALEKAIDAETMKLHHDRHHAAYVKNLN   105
```

```
WP_254449390.1       TQF-----FAVAYPDRELSASPAELPKLPYADNALEKAIDAQTMKLHHDKHHSAYVNKLN    103
MBF2015390.1         TQV-----LPVAYPDRELSASPAELPKLPYAYNALEKAIDAQTMELHHDKHHAAYVNNLN    103
MEB3216192.1         TQV-----LPVAYPDRELSASPAQLPKLPYAYNALEKAIDAQTMELHHDKHHAAYVNNLN    103
WP_193917416.1       TQV-----LPVAYPDRELSASPAQLPKLPYAYNALEKAIDAQTMELHHDKHHAAYVNNLN    103
WP_277875608.1       TDT-----RPIAFLDRQLSTNPAELPPLPYGYNALEKAIDAETMKLHHDRHHAAYVNNLN    103
MBD2346042.1         TDT-----RPIAFLDRQLSTNPAELPPLPYGYNALEKAIDAETMKLHHDRHHAAYVNNLN    90
WP_010994247.1       TDK-----RSIGFIDRQLGTNPAELPPLPYGYDALEKAIDAETMKLHHDKHHAAYVNNLN    103
Nostoc sp. PCC 7120  TDK-----RSIGFIDRQLGTNPAELPPLPYGYDALEKAIDAETMKLHHDKHHAAYVNNLN    103
WP_277884298.1       TDK-----RSIGFLDRQLGTNPAELPPLPYGYDALEKAIDAETMKLHHDKHHAAYVNNLN    103
WP_414756233.1       TDK-----RSIGFLDRPLGTNPAELPPLPYGYDALEKAIDAETMKLHHDKHHAAYVNNLN    103
WP_011318269.1       TDK-----RSIGFLDRPLGTNPAELPPLPYGYDALEKAIDAETMKLHHDKHHAAYVNNLN    103
WP_015215174.1       TDA-----RPIGYLDRQLSANPAELPPLPYADNALEKAIDAETMKLHHDKHHAAYVNNLN    103
WP_276752922.1       TDT-----RPIAFLDRQLSANPAQLPPLPYPYNALEKAIDAETMKLHHDRHHATYVENLN    104
RUR73923.1           TDT-----RPIAFLDRQLSANPAQLPPLPYPYNALEKAIDAETMKLHHDRHHATYVENLN    105
WP_016872332.1       TDT-----RPIAFLDRQLSANPAQLPPLPYPYNALEKAIDAETMKLHHDRHHATYVENLN    104
WP_341997750.1       ADT-----RPIAYLDRQLSANPAQLPPLPYSYNALEKAIDAETMKLHHNRHHATYVENLN    104
WP_289789894.1       TDT-----RPIAFLDRQLSTNPAQLPPLPYPYNALEKAIDAETMKLHHDRHHATYVENLN    103
WP_191756852.1       TNT-----RPVAVLNSQLSANPAQLPPLPYAYNALEKAIDAETMKLHHDKHHAAYVNNLN    103
WP_193198009.1       TNT-----RSVAVLNRQLSANPAQLPPLPYAYNALEKAIDAETMKLHHDKHHAAYVNNLN    103
WP_323228814.1       TNT-----RPVAYVDRQLSTNPAQLPPLPYAYNALEKAIDAETMKLHHDRHHAAYVNNLN    102
WP_103138074.1       VNT-----RPVAFVDRQLSTNPAELPPLPYAYNALEKAIDAETMKLHHDRHHATYVNNLN    103
WP_242072232.1       TNT-----RPVAYLDRQLSTNPAELAPLPYAYNALEKAIDAETMKLHHDKHHAAYVNNLN    103
MBD2440923.1         TNT-----RPVAYLDRQLSTNPAELAPLPYAYNALEKAIDAETMKLHHDKHHAAYVNNLN    88
WP_318780343.1       TSS-----VPVAYPDRELSASPAQLPKLPYAYNALEKAIDAETMKLHHDKHHAAYVNNLN    106
WP_460206421.1       TPF-----SPVAYPDRELGTSPAELPPLPYDYGALEKVIDAETMKLHHDRHHATYVNNLN    107
WP_169219177.1       ATS-----SPVAYVDKELSANPAQLPPLPYDYGALEKAIDAETMKLHHDRHHATYVNNLN    92
WP_332997608.1       ATS-----SPIAYPDKDLSASPAQLPPLPYDYGALGKAIDAETMKLHHDRHHAAYVNNLN    105
MBW4498943.1         ATS-----SPVAYPDKDLSASPAQLPPLPYDYGALGKAIDAETMKLHHDRHHAAYVNNLN    92
WP_414585734.1       ATS-----SPVAYLDKGLSASPAQLPPLPYDYGALGKAIDAETMKLHHDRHHAAYVNNLN    105
MCC5641606.1         KSS-----PPVAYPNRALSAFPAQLPPLPYAYGALGKAIDAETMKLHHDAHHASYVKNLN    96
WP_435020268.1       RSF-----PPVAYPNRALSASPAQLPPLPYAYGALGKAIDAETMKLHHDAHHASYVKNLN    98
WP_292704904.1       RSF-----APVAYPNRALSAFPAQLPPLPYGYGALGKVIDAETMKLHHDAHHASYVKNIN    98
WP_335036612.1       RSF-----PPVAYPNRALSAFPAQLPPLPYAYGALGKAIDAETMKLHHDAHHASYVKNLN    98
WP_302480924.1       RSF-----SPVAYPNRALSAFPAQLPPLPYGYGALGKAIDAETMKLHHDAHHASYVKNLN    98
WP_311774093.1       RSF-----PPVAYPNRALSAFPAQLPPLPYGYGALGKAIDAETMKLHHDAHHASYVKNLN    98
N. punctiforme 73102 SSF-----PPVAYPNRALSAFPAQLPPLPYGYGALGKAIDAETMKLHHDVHHASYVKNLN    98
WP_322745234.1       RSF-----PPVAYPNRALSAFPAQLPPLPYGYGALGKAIDAETMKLHHDAHHASYVKNLN    98
WP_306297076.1       RSF-----PPVAYPNRSLSAFPAQLPPLPYGYGALGKAIDTETMKLHHDAHHASYVKNLN    98
WP_322746338.1       RSF-----PPVAYPNRALSAFPAQLPPLPYGYGALGKAIDAETMKLHHDAHHASYVKNLN    98
WP_277883559.1       RSF-----PPVAYPNRSLSAFPAQLPPLPYGYGALGKAIDAETMKLHHDAHHATYVKNLN    98
WP_392482408.1       RSF-----PPVAYPNRALSAFPAQLPPLPYGYGALGKAIDAETMKLHHDAHHATYVKNLN    98
WP_163931552.1       RSF-----PPVAYPNRSLSAFPAQLPPLPYGYGALGKAIDAETMKLHHDAHHASYVKNLN    98
WP_114081005.1       RSF-----PPVAYPNRSLSAFPAQLPPLPYGYGALGKAIDAETMKLHHDAHHASYVKNLN    98
                                 *  * *    .*    :*   ** :**   **   *    *

E. coli              AALESLPEFANLPVEELITKLDQLPADKKTVLRNNAGGHANHSLFWKGLK--KGTTLQGD    98
G. violaceus 7421    KALEKYPDLQKKSPEQLLRDLKQVPEDTRAAVRGNGGGHVNHTMFWEIMKPKGGGEPAGP    164
WP_015156211.1       SALKKYPDLRNRSVETMLRDLDGIPEAIRTTVQNNGGGHLNHTIFWQIMSPKGGGQPTGQ    162
Synechococcus 6312   AALQKYPQWSGLRIEELLEKLDQLPNDIRQTVRNNGGGHANHSLFWESMSPAAGGKPRGK    136
WP_322876523.1       AALQKYPQWSGLRIEELLEKLDQLPNDIRQTVRNNGGGHANHSLFWESMGPAAGGEPTGN    136
WP_277866136.1       RALSEYRQWQNLAIEDLMAKISYLPSSIRQTVRNNGGGHANHSLFWQSMSPQGGGEPSQE    133
WP_011055891.1       KALTPYRQWHGLAIETLLRHLHQLPTAIQSTVRNHGGGHANHSLFWQSMAPNGGGEPAGD    133
WP_297761391.1       KALTPYRQWHGLAIEELLHHLHQLPTAIQQTVRNHGGGHANHSLFWESMAPNGGGEPTGD    133
WP_024125507.1       KALTPYRQWHGLAIEELLPHLHQLPTAIQETVRNHGGGHANHSLFWQSMAPNGGGEPTGD    133
WP_365970883.1       KALTPYRQWHGLAIEELLPHLHQLPTAIQETVRNHGGGHANHSLFWQSMAPNGGGEPTGD    133
WP_409873764.1       KALTPYRQWHSLAIEDLLRHLRQLPTAIQQTVRNHGGGHANHSLFWQSMTPKGGGEPTGD    133
WP_300872640.1       KALAPYRQWHGLPIEDLLRHIRQLPTAIQQTVRNHGGGHANHSLFWQSMAPNGGGEPTGD    133
WP_250682089.1       KALAPYRQWHGLPIEELLRHIRQLPTAIQQTVRNHGGGHANHSLFWQSMSPDGGGEPTGE    133
WP_409869042.1       KALTPYRQWHGLAIEDLLGHIRQLPTAIQQTVRNHGGGHANHSLFWQSMAPNGGGEPTGE    133
WP_297053013.1       KALTPYRQWHGLPIEDLLRHIHQLPTAIQQTVRNHGGGHANHSLFWQSMAPNGGGEPMGE    133
WP_297051732.1       KALTPYRQWHGLPIEDLLRHIHQLPTAIQKTVRNHGGGHANHSLFWQSMAPNGGGEPMGE    133
WP_181494764.1       KALTPYRQWQGLPIEDLLRHIRQLPTAIQQTVRNHGGGHANHSLFWQSMAPNGGGEPTGE    133
WP_460193008.1       KALTPYRQWQGLPIEDLLRHIRQLPTAIQQTVRNHGGGHANHSLFWQSMAPNGGGQPTGD    133
WP_286627080.1       KALTPYRQWQGLPIEDLLRHIRQLPTAIQQTVRNHGGGHANHSLFWQSMAPNGGGEPTGE    133
WP_310743007.1       KALTPYRQWQGLPIEDLLCHIRQLPTAIQQTVRNHGGGHANHSLFWQSMAPNGGGEPTGE    133
WP_206200084.1       KALTPYRQWQGLPIEDLLCHIRQLPTAIQQTVRNHGGGHANHSLFWQSMAPNGGGEPTGE    133
WP_310821562.1       KALTPYRQWQGLPIEDLLRHIRQLPTAIQQTVRNHGGGHANHSLFWQSMAPNGGGEPTGE    133
WP_310800931.1       KALTPYRQWQGLPIEDLLRHIRQLPTAIQQTVRNHGGGHANHSLFWQSMAPNGGGEPTEE    133
WP_310454098.1       KALTPYRQWQGLPIEDLLRHIRQLPTAIQQTVRNHGGGHANHSLFWQSMAPNGGGEPTGE    133
WP_302677659.1       KALTPYRQWQGLPIEDLLRHIRQLPTAIQQTVRNHGGGHANHSLFWQSMAPNGGGEPTGE    133
Pseudanabaena 7367   KAIATYPDLAGMSAEDMLRDLTQVPEPIRTTVRNNAGGHVNHSMFWEIMSPNGGGTPTGA    150
MCU0526905.1         AAIDQYPDLKLMTAEEMLRQIDTIPESIRTTVRNNGGGHVNHSMFWEIMSPTGGGSPSGA    151
MDJ0592545.1         KAVSKYPELASKSAEDILREIDTVPEDVAQTIRNNGGGYVNHQMFWEIMSPNGGGEPTGG    135
MBX5479872.1         AAIAKHPELSNRSAEDLIRNLQSVPADIREQVRNNGGGHVNHTMFWQIMKPNGGGEPTGP    140
WP_041973980.1       AAIAKHPELSNRSAEDLIRNLQSVPADIREQVRNNGGGHVNHTMFWQIMKPNGGGEPTGP    140
WP_428360810.1       AAVAKYPQLQTRSVESLIQNLNSLPEDIRTTVRNNGGGHLNHTMFWEIMGPKAGGAPTGA    135
WP_233744687.1       AAVAKYPQLQTRSVESLIQNLNSLPEDIRTTVRNNGGGHLNHTMFWEIMGPQAGGAPTGA    135
```

WP_445926464.1 AAVAKYPQLQTRSVESLIQNLNSLPEDIRITVRNNGGGHLNHTMFWEIMGPKAGGPPVGA 135
WP_015152615.1 AAIAKYPKLKGQTAEQLIQKLDSLPQDIRMTIRNNGGGHVNHSMFWEIMSPKGGGEPTGE 138
WP_190455393.1 EAVNKYPQLQGKSAEELISNLNSIPEDIRTTVRNNGGGHVNHTMFWEIMAPNAGGAPTGK 150
WP_190424133.1 EAVNKYPQLQGKSAEELISNLNSIPEDIRTTVRNNGGGHVNHTMFWEIMAPNAGGAPTGK 150
WP_027843320.1 AALKKHPELKDKSIEDLLSDLESVPSDIRRTVRNNGGGHVNHSMFWEIMGPNAGGEPTGA 147
WP_374791410.1 AAIAKYPQLKGKTVEELLQNLNTVPEDIRTTVRNNGGGHLNHSMFWRIMKPNGGGEPTGA 150
WP_374825979.1 GAIAKYPQLKGKTVEELLQNLNTVPEDIRTTVRNNGGGHLNHSMFWRIMKPNGGGEPTGA 150
WP_190465982.1 DAIAKYPELKGKTVEELLQNLNTVPEDIRTTIRNNGGGHLNHSMFWRIMKPNGGGEPTGA 150
BDA72795.1 AALDKHPQLKGKTVEQMLQNLDSVPTDIRNVVRNNGGGHVNHKMFWEIMKPKGGGEPTGA 151
GJD16125.1 AALDKHPQLKGKTVEQMLQNLEGIPADIRNVVRNNGGGHVNHKMFWEIMKPKGGGEPTGA 151
MEC4881204.1 AALDKYPELKSKSVEELLRDLNSVPEDIRRTVRNNGGGHVNHSMFWRIMTPKGGGEPQGP 141
WP_190560307.1 KALDKYPELKNRTVEELLQNLDNVPADIRTTVRNNGGGHVNHSMFWKIMKPNGGGEPTGE 134
MFM5980434.1 KALDKYPELKNKTVEQLLQNLDNVPADIRTTVRNNGGGHVNHSMFWKIMKPNGGGEPTGE 134
WP_096571931.1 KALDKYPELKNKTVEQLLQNLDNVPADIRTTVRNNGGGHVNHSMFWKIMKPNGGGEPTGE 134
WP_190648403.1 KALDKYPELKNKTVEQLLQNLDNVPADIRTTVRNNGGGHVNHSMFWKIMKPNGGGEPTGE 134
WP_190347369.1 KALDKYPELKNKTVEQLLQNLDNVPADIRTTVRNNGGGHVNHSMFWKIMKPNGGGEPTGE 134
***N. punctiforme* 73102** AALDKHPELKGKTIEELLQKLDSVPQDIRRTVRNNGGGHVNHSMFWEIMKPKGGGEPTGN 153
WP_334931430.1 AALDKHPELKGKTIEELLQKLDSVPQDIRKTVRNNGGGHVNHSMFWEIMKPKGGGEPTGN 153
WP_335056299.1 AALEKHPELKGKTIEELLQKIDNVPQDIRRTVRNNGGGHVNHSMFWQIMKPKGGGEPTGN 153
WP_322673311.1 AALEKHPELKGKTVEELLQKLDSVPQDIRKTVRNNGGGHVNHSMFWQIMKPKGGGEPTGN 153
WP_322689216.1 AALEKHPELKGKTVEELLQKLDSVPQDIRKTVRNNGGGHVNHSMFWQIMKPKGGGEPTGN 153
WP_322701324.1 AALEKHPELKGKTVEELLQKLDSVPQDVRKTVRNNGGGHVNHSMFWQIMKPKGGGEPTGN 153
WP_335012934.1 AALDKHPELKGKTIEELLRKLDNVPQDIRRVVRNNGGGHVNHSMFWEIMKPKGGGEPTGN 149
WP_292868329.1 AALAKHPELKGKTVEELLQKLDNVPQDIRKTVRNNGGGHVNHSMFWEIMKPKGGGEPTGN 161
WP_334926809.1 AALAKHPELKGKTVEELLQKLDNVPQDIRKTVRNNGGGHVNHSMFWEIMKPKGGGEPTGN 161
WP_109007648.1 AALAKHPELKGKSVEELLRKLDNVPQDIRNTVRNNGGGHVNHSMFWQIMKPKGGGEPTGN 149
WP_289795776.1 AALEKHPELKGKTVEELLQKLDKVPQDIRRTVRNNGGGHVNHSMFWQIMKPKGGGEPTGN 149
WP_099100092.1 AALEKHPELKGKTVEELLQKLDSVPQDIRRTVRNNGGGHVNHSMFWQIMKPKGGGEPTGN 149
WP_335201774.1 AALEKHPELKGKTVEELLQKLDNVPQDIRRTVRNNGGGHVNHSMFWQIMKPKGGGEPTGN 153
MDZ7959988.1 AALDKHPQLKGKTVEEILRKLDTVPADIRKTVRNNGGGHVNHSMFWQIMKPQGGGEPTGA 146
MBD2208974.1 AALEKHPQLKGKTVEELLRQLDKVPADIRKTLRNNGGGHVNHSMFWQIMKPKGGGEPTGA 145
MBD2339284.1 AALEKHPQLKGKTVEELLRQLDKVPADIRKTLRNNGGGHVNHSMFWQIMKPKGGGEPTGA 145
MBD2166265.1 AALEKHPQLKGRTVEELLRQLDKVPADIRKTVRNNGGGHVNHSMFWQIMKPKGGGEPTGA 145
WP_167724335.1 AALEKHPQLKGRTVEELLRQLDKVPADIRKTLRNNGGGHVNHSMFWQIMKPKGGGEPTGA 145
WP_045869977.1 AALEKHPQLKGRTVEELLRQLDKVPADIRKTVRNNGGGHVNHSMFWQIMKPKGGGEPTGA 146
WP_096582728.1 AALEKHPQLKGRTVEELLRQLDKVPADIRKTVRNNGGGHINHSMFWQIMKPKGGGEPTGA 145
WP_096599871.1 AALEKHPELKSRSVEELLRKLDKVPEDIRKIVRNNGGGHVNHSMFWQIMKPQGGGEPTGA 146
WP_190743315.1 AALEKYPQLKSRSVEDLLRQLDRVPEDIRKTVRNNGGGHVNHSMFWQIMKPQGGGEPTGA 146
WP_190544005.1 AALEKYPKLKGKTVEDLLRQLDRVPEDIRKTVRNNGGGHVNHSMFWQIMKPQGGGEPTGA 145
WP_096643960.1 AALEKYPKLKGRTVEDLLRQLDRVPEDIRKTVRNNGGGHVNHSMFWQIMKPQGGGEPTGA 145
***Cyanothece* 7425** AALQQDTNLQNQSVEALLRDLNSVPEAIRERVRNNGGGHLNHTIFWQIMGPDGGGEPTGD 159
WP_414578860.1 DALKKYPNLQKRSVEALLRDLDSVPEDIRTKVRNNGGGHLNHTIFWQIMSPDGGGQPTGK 154
WP_323267495.1 DALKKYPNLQKRSVEALLRDLDSVPEDIRTKVRNNGGGHLNHTIFWQIMSPDGGGQPTGK 154
WP_214431594.1 DALTKYPDLQKNSVEALLKDLNKVPEDIRTKVRNNAGGHLNHTIFWQIMSPQGGGEPTGA 160
WP_322662701.1 NALKRYPDLQKNSVEALLKDLNKVPEDIRTKVRNNGGGHLNHTIFWQIMSPQGGGEPTGA 161
RCJ28434.1 DALKRYPDLQKNSVEALLKDLNKVPEDIRTKVRNNGGGHLNHTIFWQIMSPQGGGEPTGA 161
MBE9051467.1 DALKQESGLQKSSVEALLRDLNSVPENIRTKVRNNGGGHLNHTIFWQIMSPEGGGEPTGE 157
MBW4558831.1 DALKQEPDLQKSSVEALLRDLNSVPENIRTKVRNNGGGHLNHTIFWQIMSPEGGGEPTGE 141
WP_063871086.1 DALKQAPDLQKNSVEALLRDLNNVPENIRTKVRNNGGGHLNHTIFWQIMSPEGGGEPIDE 157
WP_006196478.1 DALKQAPDLQKNSVEALLRDLNNVPENIRTKVRNNGGGHLNHTIFWQIMSPEGGGEPIEE 157
WP_271918904.1 DALKQAPDLQKNSVEALLRDLNNVPENIRTKVRNNGGGHLNHTIFWQIMSPEGGGEPIDE 157
WP_271942563.1 DALKQAPDLQKNSVEALLRDLNNVPENIRTKVRNNGGGHLNHTIFWQIMSPEGGGEPIDE 157
WP_272068383.1 DALKQEPDLQKSSVEALLRNLNNIPENIRTKVRNNGGGHLNHTIFWQIMSPDGGGEPTGE 157
WP_193998321.1 DALKKEPDLQKSSVEALLRDLNNVPESIRTDVRNNGGGHLNHTIYWQIMSPEGGGEPTEE 156
WP_089093562.1 DALKEEPELQKSSVEALLRDLNKVPENIRTKVRNNAGGHLNHTIFWQIMSPDGGGEPTDE 157
WP_323215458.1 DALKQEPDLQESSVEALLRDLNKVPENIRTKVRNNGGGHLNHTIFWQIMSPDGGGEPTGE 157
WP_414550770.1 DALKQEPELQKSSVEALLRDLNKVPENIRTKVRNNAGGHLNHTIFWQIMSPDGGGEPTDE 157
WP_214442418.1 DVLKNYPNLQKRSIEALLRDLNSVPADIRTKVRDNGGGHLNHTMFWQIMSPKGGGEPTGA 157
MEA5507287.1 DALKNYPNLRKRSVEALLKDLNSVPEDIRTKVRNNGGGHLNHTMFWQIMSPQGGGEPTGA 157
WP_281484005.1 DALKNYPNLRKKSVEALLKDLNSVPEDIRTKVRNNGGGHVNHTMFWQIMSPQGGGEPKGA 159
WP_162421941.1 DALKDQPNVQKLSVEAMLRDLNQVPEAIRQTVRNNAGGHLNHTIFWQIMAPNGGGQPTGA 157
WP_430678559.1 SALKDQPNLQKLSVEAMLRDLNAVPENLRQRVRNNAGGHLNHTIFWQIMSPNGGGQPTGA 156
MBE9012496.1 SALKDQPNLQKLSIEAMLLNLNAVPENIRQRVRNNAGGHLNHTIFWQIMSPTGGGQPTGA 156
WP_059000726.1 SALKDQPNLQKLSIEAMLRDLNAVPENIRQKVRNNAGGHLNHTIFWQIMSPNGGGQPTGA 156
WP_482030890.1 NALKQQSDLRSRSIEAMLRDLNSVPEEIRTTVRNNGGGHLNHTLFWRLMSPDGGGQPTGE 147
WP_445299659.1 EALKQQSNLQNRSVEALLRDLNSVPEAIRTTVQNNGGGHLNHTIFWQIMSPNGGGQPTGA 161
WP_442927815.1 EALKQQSNLQNRSVEALLRDLNSVPEAIRTTIRNNGGGHLNHTIFWQIMSPNGGGQPTGA 161
WP_445299397.1 EALKQQSNLQNRSVEALLRDLNSVPEAIRTTVQNNGGGHLNHTIFWQIMSPNGGGQPTGA 155
WP_445312326.1 EALKQQSNLQNRSVEALLRDLNSVPEAIRTTVQNNGGGHLNHTIFWQIMSPNGGGQPTGA 155
HIK16311.1 EALKPQSNLQNRSVEALLRDLNSLPEDIRTTVRNNGGGHLNHTIFWQLMSPNGGGQPTGE 152
WP_432812051.1 AALTQQPNLQNRSVEALLRDLNSLPEAIRTTVQNNAGGHLNHTIFWQIMSPNGGGQPTGE 152
WP_431655370.1 EALKEQSDLRNRSVEALLRDLNSVPEGIRTTVQNNAGGHLNHTIFWQIMSPNGGGQPTGE 153
MGI0483888.1 EALKEQSDLRNRSVEALLRDLNSVPEGIRTTVQNNAGGHLNHTIFWQIMSPNGGGQPTGE 138
MBW4417879.1 EALKDQPQVQNRSVEALLRDLNSVPESIRMKVRNNAGGHLNHTIFWQIMSPDGGGQPTGA 153
MBW4540880.1 EALKDQPQVQNRSVEALLKDLNSVPEPIRMKVRNNAGGHLNHTIFWQIMSPNGGGQPTGA 153
MBW4644872.1 EALQKYPELKNRSVEAMLRDLNSIPEDIRQTVRNNGGGHLNHTIFWQLMSPDGGGQPTGE 163
PMB52031.1 EALQKYPELQNRSVEAMLRDLNSLPEDIRETVRNNGGGHLNHTIFWQIMSPNGGGEPTGE 164

```
WP_190751173.1        EALQKYPELKNRSVEAMLRDLNNIPEDIRDTVRNNGGGHLNHTIFWQIMSPNGGGQPTGE    164
WP_190549236.1        EALQKHPELKNRSVEAMLRDLNNIPEDIRETVRNNGGGHLNHTIFWQIMSPNGGGQPTGE    164
WP_190629649.1        EALQKYPELKNRSVEAMLRDLNNIPEDIRDTVRNNGGGHLNHTIFWQIMSPNGGGQPTGE    164
MBD2168452.1          EALQKHPELKNRSVEAMLRDLNSIPEDIRETVRNNGGGHLNHTIFWQIMSPNGGGQPTGE    164
MBD2342527.1          EALQKHPELKNRSVEAMLRDLNSIPEDIRETVRNNGGGHLNHTIFWQIMSPNGGGQPTGE    164
MHN9116841.1          EALQKHPELKNRSVEAILRDLNSIPEDIRETVRNNGGGHLNHTIFWQIMSPNGGGQPTGE    165
MFN6514993.1          EALQKHSELKNKSVEAMLRDLNSIPEDIRETVRNNGGGHLNHTIFWQIMSPTGGGQPTGE    164
NDJ23487.1            EALQKHSELKNRSVEAMLGDLNSIPEDIRETVRNNGGGHLNHTIFWQIMSPNGGGQPTGE    164
WP_181929818.1        EALQKHSELKNRSVEAMLRDLNSIPEDIRETVRNNGGGHLNHTIFWQLMSPNGGGQPTGE    164
MBV6623186.1          AALQKYPELQNKSVEAMLRDLNSVPEDIRETVRNNGGGHLNHTIFWQIMSPNGGGEPTGE    164
WP_323238745.1        AALQKYPELQNKSVEAMLRDLNSIPEDIRETVRNNGGGHLNHTILWQIMSPNGGGEPTGD    165
WP_254449390.1        EALQKHPELKSRSVEAILRDLNSVPEDIRQTVRNNGGGHLNHTIFWQIMSPNGGGEPTGK    163
MBF2015390.1          TALQKHPELKSRSVEVMLRDLDSIPEDIRETVRNNGGGHLNHTIFWQIMSPNGGGEPTGQ    163
MEB3216192.1          EALQKHPELKNSSVESMLRDLDNIPEDIRETVRNNAGGHLNHTIFWQIMSPDGGGEPTGQ    163
WP_193917416.1        EALQKHPELKNRSVEAMLRDLDSIPEDIRATVRNNGGGHLNHTIFWQIMSPNGGGEPTGQ    163
WP_277875608.1        NALKKHTELQNRSVEALLRDLNSVPEDIRTTVRNNGGGHLNHTIFWQIMSPEGGGQPTGE    163
MBD2346042.1          NALKKHTELQNRSVEALLRDLNSVPEDIRTTVRNNGGGHLNHTIFWQIMSPEGGGQPTGE    150
WP_010994247.1        NALKKHPELQNSSVEALLRDLNSVPEDIRTTVRNNGGGHLNHTIFWQIMSPDGGGQPTGD    163
Nostoc sp. PCC 7120   NALKKHPELQNSSVEALLRDLNSVPEDIRTTVRNNGGGHLNHTIFWQIMSPDGGGQPTGD    163
WP_277884298.1        NALKKHPELQNSSVEALLRDLNSVPEDIRNTVRNNGGGHLNHTIFWQIMSPDGGGEPTGD    163
WP_414756233.1        NALKKHPELQNSSVEALLRDLNSVPEDIRNTVRNNGGGHLNHTIFWQIMSPNGGGQPTGD    163
WP_011318269.1        NALKKHPELQNSSVEALLRDLNSVPEDIRNTVRNNGGGHLNHTIFWQIMSPNGGGQPTGD    163
WP_015215174.1        NALKTQPNLQNQSVEALLRDLNNVPENIRGTVRNNGGGHLNHTIFWQIMSPQGGGQPTGE    163
WP_276752922.1        NALNQYPQLQDRSVEALLRDLNSVPEDIRTTVRNNGGGHLNHTIFWQIMSPQGGGEPTGE    164
RUR73923.1            NALNQYPQLQDRSVEALLRDLNSVPEDIRTTVRNNGGGHLNHTIFWQIMSPQGGVEPTGE    165
WP_016872332.1        NALNQYPQLQDRSVEALLRDLNSVPEDIRTTVRNNGGGHLNHTIFWQIMSPQGGVEPTGE    164
WP_341997750.1        EALKQYPQLQDRSVEALLRDLNGVPEDIRTAVRNNGGGHLNHTIFWQIMSSQGGGEPTGE    164
WP_289789894.1        EALKKYPQLQDRSVEALLRDLNSVPEDIRTAVRNNGGGHLNHTIFWQIMSPQGGGQPTGE    163
WP_191756852.1        DALKKYPQLQNSSVESLLRDLNSVPEDIRTKVRNNGGGHLNHTIFWQIMSPQGGGQPTGD    163
WP_193198009.1        NALKKYPQLQNSSVESLLRDLNSVPEDIRTTVRNNGGGHLNHTIFWQIMSPQGGGQPTGD    163
WP_323228814.1        DALKKYPQLQNSSVETLLRDLNSIPEEIRTTVRNNGGGHLNHTIFWQIMSPQGGGQPTGE    162
WP_103138074.1        NALKKYPQLQNSSVEALLQDLNSIPEDIRTTVRNNGGGHLNHTIFWQIMSPDGGGEPTGE    163
WP_242072232.1        NALKKYRQLQKSSVEALLLDLNSIPEDIRTTVRNNGGGHLNHTIFWQIMSPKGGGEPTGE    163
MBD2440923.1          NALKKYRQLQKSSVEALLLDLNSIPEDIRTTVRNNGGGHLNHTIFWQIMSPKGGGEPTGE    148
WP_318780343.1        NALKKYSDLQNRSVEAILRDLESVPEDIRTTVRNNGGGHLNHTIFWQIMSPNGGGQPTGE    166
WP_460206421.1        DALKKHPELQNRSVESLLRDLDNVPEDIRATVRNNGGGHLNHTIFWQIMSPQGGGQPTGV    167
WP_169219177.1        DALTKHPEQQNRSVEALLGDLDSVPEDIRTTVRNNAGGHLNHTVFWQIMSPQGGGEPTGQ    152
WP_332997608.1        EALKEHPELQNRSVEALLRDLNSVPEDIRTTVRNNGGGHLNHTIFWQIMSPQGGGQPTGT    165
MBW4498943.1          EALKEHPDLQNRSIEALLRDLNSVPEDIRTTVRNNGGGHLNHTIFWQIMSPQGGGQPTGT    152
WP_414585734.1        EALKEHPDLQNRSVEALLRDLDSVPEDIRTTVRNNGGGHLNHTIFWQIMSPQGGGQPTGA    165
MCC5641606.1          DALKQYPDLQKRSVEALLVNLNSVPEDIRTKVRNNGGGHLNHTIFWQLMSPEGGGKPSGA    156
WP_435020268.1        DALKQYPDLQKRSVEALLVDLNSVPEDIRTKVRNNGGGHLNHTIFWQLMSPEGGGQPSGA    158
WP_292704904.1        DALKQYPDLQKRSVEALLVDLNSVPEDIRAKVRNNGGGHLNHTIFWQLMSPDGGGQPTGA    158
WP_335036612.1        DALKQYPDLQKRSVEALLVDLNSVPEAIRTKVRNNGGGHLNHTIFWQLMSPDSGGQPTGA    158
WP_302480924.1        DALKQYPDLQKRSVEGLLVDLNSVPEDIRTKVRNNGGGHLNHTIFWQLMSPDSGGQPTGA    158
WP_311774093.1        DALKQYPDLQKRSVEALLVDLNSVPEDIRTKVRNNGGGHLNHTIFWQLMSPDSGGQPTGA    158
N. punctiforme 73102  DALKQYPDLQKRSVEALLVDLKSVPEDIRTKVRNNGGGHLNHTIFWQLMSPDSGGQPTGA    158
WP_322745234.1        DALKQYPDLQKRSVEALLVDLNSVPEDIRTKVRNNGGGHLNHTIFWQLMSPDSGGQPTGA    158
WP_306297076.1        DALKQYPDLQKRSVEALLVDLNSVPEDIRTKVRNNGGGHLNHTIFWQLMSPDSGGQPTGA    158
WP_322746338.1        DALKQYPDLQKRSVEALLVDLKSVPEDIRTKVRNNGGGHLNHTIFWQLMSPDSGGQPTGA    158
WP_277883559.1        DALKQYPDLQKRSVEALLVDLNSVPEDIRTKVRNNGGGHLNHTIFWQLMSPDSGGQPTGA    158
WP_392482408.1        DALKQYPDLQKRSVEALLVDLNSVPEDIRTKVRNNGGGHLNHTIFWQLMSPDSGGQPTGA    158
WP_163931552.1        DALKQYPDLQKRSVEALLVDLNSVPEDIRTKVRNNGGGHLNHTIFWQLMSPDSGGQPTGA    158
WP_114081005.1        DALKQYPDLQKRSVEALLVDLNSVPEDIRTKVRNNGGGHLNHTIFWQLMSPDSGGQPTGA    158
                       .:           * ::  .:  :*       ::.:.**: **  : *.  :     *

E. coli               LKAAIERDFGSVDNFKAEFEKAAASRFGSGWAWLVLK-GDKLAVVSTANQDSPLMGEAIS    157
G. violaceus 7421     IAAAIRTNFGSFDAFKTQFNEAGTKRFGSGWVWLVSNPGGKLEIVSTANQDSPIAD----    220
WP_015156211.1        IAQEINKTFGSFEKFREQFNAAGGDRFGSGWVWLVRNAKGQLEITSMPNQDNPIMK----    218
Synechococcus 6312    LATAMQARFGSFEDFQAQFNSAGLKQFGSGWVWLGLGQDGTLQVFTTPNQDSPLSQ----    192
WP_322876523.1        LATAIQARFGSFGDFQAQFNSAGLKQFGSGWVWLGLGQDGTLQVFTTPNQDSPLSQ----    192
WP_277866136.1        LGGAIAETFSSFQDFQGQFNGAGLGQFGSGWVWLVVTPAKQLAILTTPNQDSPLMD----    189
WP_011055891.1        LAIAIENTFGSFPEFQAQFQQAGLRHFGSGWVWLVLTPQGNLTITTTLNQDSPLMQ----    189
WP_297761391.1        LAIAIEDTFGSFPEFQTQFQQAGLQHFGSGWVWLVLTPQGNLTITTTLNQDSPLMQ----    189
WP_024125507.1        LAIAIEDTFGSFPEFQTQFQQAGLQHFGSGWVWLVLTPQGNLTITTTLNQDSPLMQ----    189
WP_365970883.1        LAIAIEDTFGSFPEFQTQFQQAGLQHFGSGWVWLVLTPQGNLTITTTLNQDSPLMQ----    189
WP_409873764.1        LATAIVATFGSFSEFQTQFQQAGLKHFGSGWVWLVLTPQGHLEITTTLNQDSPLIQ----    189
WP_300872640.1        LATAIAATFGSFSEFQAQFQQAGLKHFGSGWVWLVLTPQGNLTITTTLNQDSPLMQ----    189
WP_250682089.1        LATAIAATFGSFSEFQAQFQQAGLKHFGSGWVWLVLTPQGNLAITTTLNQDSPLMQ----    189
WP_409869042.1        LATAIAATFGSFAEFQTQFQQAGLKHFGSGWVWLVLTPQGTLAITTTLNQDSPLME----    189
WP_297053013.1        LATAIAATFGSFSEFQTQFQQAGLKHFGSGWVWLVLTPQGNLAITTTLNQDSPLMQ----    189
WP_297051732.1        LATAIAATFGSFSEFQTQFQQAGLKHFGSGWVWLVLTPQGNLAITTTLNQDSPLMQ----    189
WP_181494764.1        LATAIAATFGSFAEFQTQFQQAGLKHFGSGWVWLVLTPQGNLAITTTLNQDSPLME----    189
WP_460193008.1        LATAIATTFGSFSEFQTQFQQAGLKHFGSGWVWLVLTPQGNLAITTTLNQDSPLMQ----    189
WP_286627080.1        LATAIAATFGSFSAFQTQFQQAGLKHFGSGWVWLVLTPQGNLAITTTLNQDSPLMQ----    189
WP_310743007.1        LATAIATTFGSFSAFQTQFQQAGLKHFGSGWVWLVLTPQGNLALTTTLNQDSPLMQ----    189
WP_206200084.1        LATAIATTFGSFSAFQTQFQQAGLKHFGSGWVWLVLTPQGNLAITTTLNQDSPLMQ----    189
```

| | | |
|---|---|---|
| WP_310821562.1 | LATAIATTFGSFSAFQTQFQQAGLKHFGSGWVWLVLTPQGNLALTTTLNQDSPLMQ---- | 189 |
| WP_310800931.1 | LATAIATTFGSFSAFQTQFQQAGLKHFGSGWVWLVLTPQGNLAITTTLNQDSPLMQ---- | 189 |
| WP_310454098.1 | LATAIATTFGSFSAFQTQFQQAGLKHFGSGWVWLVLTPQGNLAITTTLNQDSPLMQ---- | 189 |
| WP_302677659.1 | LATAIATTFGSFSAFQTQFQQAGLKHFGSGWVWLVLTPQGNLAITTTLNQDSPLMQ---- | 189 |
| ***Pseudanabaena* 7367** | IAAMIAATFGNFENLQTAFNQAGASRFGSGWAWLVLDKQGNLKVTSTANQDSPLLE---- | 206 |
| MCU0526905.1 | LATAIRQAFGSFEAFQTTFNTAGAKQFGSGWAWLVMNPTGDLEVISTPNQDSPLMQ---- | 207 |
| MDJ0592545.1 | LAEAITAKFGSFEAFKTQFNDAGSKQFGSGWAWLVLDSNGELQVMGTPNQDSPLME---- | 191 |
| MBX5479872.1 | IAEAIKRTFGSFAEFKKQFNDAGMKRFGSGWVWLVRDRAGRLQIISTPNQDSPLTQ---- | 196 |
| WP_041973980.1 | IAEAIKRTFGSFAEFKKQFNDAGMKRFGSGWVWLVRDRAGRLQIISTPNQDSPLTQ---- | 196 |
| WP_428360810.1 | IATAINESFGSFEEFKKQFNDAGAKRFGSGWAWLTIDRTGKLQVSSTANQDSPLME---- | 191 |
| WP_233744687.1 | IATAINESFGSFEEFKKQFNDAGAKRFGSGWAWLTIDRMGKLQVSSTANQDSPLME---- | 191 |
| WP_445926464.1 | IATAINESFGSFEEFKKQFNDAGAKRFGSGWAWLTIDRAGKLQVSSTANQDSPLME---- | 191 |
| WP_015152615.1 | IASVIKKNFGSFANFQQQFNQAGEKQFGSGWAWLVRTPDGKFQVTSTANQDSPLME---- | 194 |
| WP_190455393.1 | IADAINQTFGSFDAFKKQFNEAGSKRFGSGWAWLARGKDGKLQVVSTANQDSPLME---- | 206 |
| WP_190424133.1 | IADAINQTFGSFDAFKKQFNEAGSKRFGSGWAWLVRGKDGKLQVVSTANQDSPLME---- | 206 |
| WP_027843320.1 | IAEAIKKEFGSFEKFQGEFNEAGKSRFGSGWAWLVLGEDQKLEVMTTANQDSPFSE---- | 203 |
| WP_374791410.1 | IADAIKTNFGSFAAFKQQFNDAGAKRFGSGWAWLVRTKDGKLQVTSTANQDSPLMD---- | 206 |
| WP_374825979.1 | IADAIKTNFGSFAAFKQQFNDAGAKRFGSGWAWLVRTKDGKLQVTSTANQDSPLMD---- | 206 |
| WP_190465982.1 | IADAIKTNFGSFAAFKQQFNDAGAKRFGSGWAWLVRTKDGKLQVTSTTNQDSPLMD---- | 206 |
| BDA72795.1 | LAAAIAENFGSFADFKKKFNTEGGKRFGSGWVWLVRNKDGKLEVVSTANQDSPISA---- | 207 |
| GJD16125.1 | LAAAIAENFGSFADFKKKFNTEGGKRFGSGWVWLVRNKDGKLEVVSTANQDSPISA---- | 207 |
| MEC4881204.1 | IASAISKNFGSFAAFKKQFNEAGTSRFGSGWVWLVLNQDDKLEVMTTANQDSPLSE---- | 197 |
| WP_190560307.1 | IATAIKDNFGSFADFKKQFNEAGGSRFGSGWVWLVRNKNGKLEITTTANQDSPLME---- | 190 |
| MFM5980434.1 | IATAINDNFGSFADFKKQFNEAGGGRFGSGWVWLVRNKNGKLEITTTANQDSPLME---- | 190 |
| WP_096571931.1 | IATAINDNFGSFADFKKQFNEAGAGRFGSGWVWLVRNKNGKLEITTTANQDSPLME---- | 190 |
| WP_190648403.1 | IATAINDNFGNFADFKKQFNEAGAGRFGSGWVWLVRNKNGKLEITTTANQDSPLME---- | 190 |
| WP_190347369.1 | IATAINDNFGNFADFKKQFNEAGAGRFGSGWVWLVRNKNGKLEITTTANQDSPLME---- | 190 |
| ***N. punctiforme* 73102** | IASAINQSFGSFAAFKKQFNEAGASRFGSGWVWLVRNKGGKLEVTTTANQDSPLST---- | 209 |
| WP_334931430.1 | IASAINQSFGSFAAFKKQFNEAGAGRFGSGWVWLVRNKGGKLEVTTTANQDSPLSA---- | 209 |
| WP_335056299.1 | IASAINQSFGSFAAFKKQFNEAGAGRFGSGWVWLVRSKGGKLEVTTTANQDTPLSA---- | 209 |
| WP_322673311.1 | IASAINQNFGSFAAFKKQFNEAGAGRFGSGWVWLVRNKGGKLEVTTTANQDTPLSA---- | 209 |
| WP_322689216.1 | IASAINQNFGSFAAFKKQFNEAGAGRFGSGWVWLVRNKGGKLEVTTTANQDTPLSA---- | 209 |
| WP_322701324.1 | IASAINQNFGSFAAFKKQFNEAGAGRFGSGWVWLVRNKGGKLEVTTTANQDTPLSA---- | 209 |
| WP_335012934.1 | IASAINQSFGSFAAFKKQFNEAGVSRFGSGWVWLVRNKDGKLEVTTTANQDSPLSS---- | 205 |
| WP_292868329.1 | IASAINQNFGSFAAFKKQFNEAGAGRFGSGWVWLVRNKGGKLEVATTANQDSPLSA---- | 217 |
| WP_334926809.1 | IASGINQNFGSFAAFKKQFNEAGAGRFGSGWVWLVRNKDGKLEVTTTGNQDSPLSA---- | 217 |
| WP_109007648.1 | IASAINQNFGSFAAFKKQFNEAGAGRFGSGWVWLVHNKGGKLEVTTTANQDTPLSA---- | 205 |
| WP_289795776.1 | IASAINQSFGSFAAFKKQFNEAGAGRFGSGWVWLVRNKDGKLEVTTTANQDTPLSA---- | 205 |
| WP_099100092.1 | IASAINQSFGSFAAFKKQFNEAGAGRFGSGWVWLVRNKDGKLEVTTTANQDTPLSA---- | 205 |
| WP_335201774.1 | IASAINQNFGSFAAFKKQFNEAGAGRFGSGWVWLVRNKGGKLEVTTTANQDTPLSA---- | 209 |
| MDZ7959988.1 | IASAINENFGNFAAFKKQFNEAGAGQFGSGWVWLVRNKNGKLEVTTTTNQDTPLSQ---- | 202 |
| MBD2208974.1 | IATAINENFGNFAAFKKQFNEAGAGQFGSGWVWLVRNKNGKLEVTTTANQDTPLSQ---- | 201 |
| MBD2339284.1 | IATAINENFGNFAAFKKQFNEAGAGQFGSGWVWLVRNKNGKLEVTTTANQDTPLSQ---- | 201 |
| MBD2166265.1 | IATAINENFGNFAAFKKQFNEAGAGQFGSGWVWLVRNKDGKLEVTTTANQDTPLSQ---- | 201 |
| WP_167724335.1 | IATAINENFGNFAAFKKQFNEAGAGQFGSGWVWLVRNKNGKLEVTTTANQDTPLSQ---- | 201 |
| WP_045869977.1 | IATAINQNFGNFAAFKKQFNEAGAGQFGSGWVWLVRNKNGKLEVTTTANQDTPLSQ---- | 202 |
| WP_096582728.1 | IATAINQNFGNFAAFKKQFNEAGAGQFGSGWVWLVRNKNGKLEVTTTANQDTPFSQ---- | 201 |
| WP_096599871.1 | IATAIKENFGNFADFKKQFNAAGAAQFGSGWVWLVRNRNGKLEVTTTANQDTPLSQ---- | 202 |
| WP_190743315.1 | IASAIKENFGSFADFKKQFNAAGAAQFGSGWVWLVRNKTGKLEVTTTANQDTPLSQ---- | 202 |
| WP_190544005.1 | IASAIKENFGSFADFKKQFNAAGAAQFGSGWVWLVRNKAGKLEVTTTANQDTPLSQ---- | 201 |
| WP_096643960.1 | IASAIKENFGSFADFKKQFNAAGAAQFGSGWVWLVRNKAGKLEVTTTANQDTPLSQ---- | 201 |
| ***Cyanothece* 7425** | LAQEINQTFGSFEELRKQFNQAGGDRFGSGWVWLVRNPQGQLQIVTTANQDNPITD---- | 215 |
| WP_414578860.1 | LAEEINQTFGSFDAFKTQFNAAGAGRFGSGWVWLVRNPQGQLQIMTTANQDSPIME---- | 210 |
| WP_323267495.1 | LAEEINQTFGSFDAFKTQFNAAGASRFGSGWVWLVRNPQGQLQIMTTANQDSPIME---- | 210 |
| WP_214431594.1 | IAQEINQTFGSFDAFKKQFNTAGGDRFGSGWVWLVRNSQGKLQIMNTPNQDNPITE---- | 216 |
| WP_322662701.1 | IAQEINQTFGSFDAFKKQFNAAGGDRFGSGWVWLVRNPQGKLQIVSTANQDNPITE---- | 217 |
| RCJ28434.1 | IAQEINQTFGSFDAFKKQFNVAGGDRFGSGWVWLVRNPQGKLQIVSTANQDNPITE---- | 217 |
| MBE9051467.1 | IAQEINRTFGGFDEFKKQFNQAGGDRFGSGWVWLVRNPAGQLQIVTTANQDNPIMD---- | 213 |
| MBW4558831.1 | IAQEINRTFGSFDEFKKQFNQAGGDRFGSGWVWLVRNPAGQLQIVTTANQDNPIMD---- | 197 |
| WP_063871086.1 | IAAEINQTFGSFEEFKKQFNAAGGDRFGSGWVWLVRNPAGELQIVTTANQDNPIMD---- | 213 |
| WP_006196478.1 | IAAEINQTFGSFEEFKKQFNAAGGDRFGSGWVWLVRNPAGELQIVTTANQDNPIMD---- | 213 |
| WP_271918904.1 | IAAEINQTFGSFEEFKKQFNAAGGDRFGSGWVWLVRNPAGELQIVTTANQDNPIMD---- | 213 |
| WP_271942563.1 | IAAEINQTFGSFEEFKKQFNAAGGDRFGSGWVWLVRNPAGELQIVTTANQDNPIMD---- | 213 |
| WP_272068383.1 | IAAEINQTFGSFDQFKQQFNAAGGDRFGSGWVWLVRNPAGELQIVTTANQDNPIMD---- | 213 |
| WP_193998321.1 | IAAEINQTFGSFDQFKEQFNQAGGDRFGSGWVWLVRNPAGELQILTTANQDNPIMD---- | 212 |
| WP_089093562.1 | IAAEINQTFGSFDQFKEQFNAAGGDRFGSGWVWLVRNPAGELQIITTANQDNPVMD---- | 213 |
| WP_323215458.1 | IAAEINQTFGSFDEFKKQFNQAGGDRFGSGWVWLVRNPAGELQIVTTANQDNPIMD---- | 213 |
| WP_414550770.1 | IAAEINQTFGSFDQFKQQFNAAGGDRFGSGWVWLVRNPAGQLQIVSTANQDNPIMD---- | 213 |
| WP_214442418.1 | IAKEINQTFGSFDAFKKQFNAAGSDRFGSGWVWLVRNPQGKLQIVTTANQDNPIID---- | 213 |
| MEA5507287.1 | IAQEINQTFGSFDAFKKQFNAAGGDRFGSGWVWLVRNPQGKLQIVTTANQDSPITD---- | 213 |
| WP_281484005.1 | IAKEINQTFGSFDDFKKQFNAAGGDRFGSGWVWLVRNPQGKLQITTTANQDSPITD---- | 215 |
| WP_162421941.1 | IAQEINQTFGSFEEFRKQFNAAGSDRFGSGWVWLVRNPQGQLQIVSTPNQDNPISD---- | 213 |
| WP_430678559.1 | IAQSINQTFGNFENFRKQFNQAGGDRFGSGWVWLVRNPQGQLQIVSTPNQDNPISE---- | 212 |
| MBE9012496.1 | IAQAINQTFGSFENFRKQFNQAGSDRFGSGWVWLVRNPQGQLQIVSTANQDNPISE---- | 212 |
| WP_059000726.1 | IAQVINQTFGNFESFRKQFNQAGSDRFGSGWVWLVRNPQGQLQIVSTANQDNPISE---- | 212 |
| WP_482030890.1 | IAQAINQSFGSFDSFKKQFNEAGGDQFGSGWVWLVRNSQGQLQIVSTPNQDNPIMD---- | 203 |
| WP_445299659.1 | VAEAINQTFGSFENFRKQFNEAGGDRFGSGWVWLVRNSQGQLQITTTPNQDNPITE---- | 217 |

```
WP_442927815.1       VAEAINQTFGSFENFRKQFNEAGGGRFGSGWVWLVRNSQGQLQITTTPNQDNPITE----   217
WP_445299397.1       VAEAINQTFGSFENFRKQFNEAGGGRFGSGWVWLVRNSQGQLQITTTPNQDNPITE----   211
WP_445312326.1       VAEAINQTFGSFENFRKQFNEAGGGRFGSGWVWLVRNSQGQLQITTTPNQDNPITE----   211
HIK16311.1           VAQAITQTFGSFDNFKKEFNEAGGDRFGSGWVWLVRNGQGQLQIVSTPNQDNPIME----   208
WP_432812051.1       IAQAINQSFGSFDNFRKQFNQAGGDRFGSGWVWLVRNPQGQLQITTTPNQDNPITT----   208
WP_431655370.1       IAQAINQNFGSFDNFRKQFNEAGGDRFGSGWVWLVRNAQGQLQITTTANQDSPIME----   209
MGI0483888.1         IAQAINQNFGSFDNFRKQFNEAGGDRFGSGWVWLVRNAQGQLQITTTANQDSPIME----   194
MBW4417879.1         IAQEINQTFGSFENFQKQFNVAGGDRFGSGWVWLVRNGQGQLQITTTPNQDNPISE----   209
MBW4540880.1         IAQEINQTFGSFENFQKQFNAAGGDRFGSGWVWLVRNGQGQLQITTTPNQDNPISE----   209
MBW4644872.1         IAQQINQTFGSFDAFRKQFNEAGGDRFGSGWVWLVRNGQNQLQIVTTPNQDNPITD----   219
PMB52031.1           IAQQINQTFGSFDAFRKQFNQEGEDRFGSGWVWLVRNAQNQLQIVNTPNQDSPIME----   220
WP_190751173.1       IAQQINQTFGSFDAFRKQFNEAGGDRFGSGWVWLVRNAQNQLQIVTTANQDSPIMD----   220
WP_190549236.1       IAQQINQTFGSFDAFRKQFNEAGGDRFGSGWVWLVRNAQNQLQIVTTANQDSPIMD----   220
WP_190629649.1       IAQQINQTFGSFDAFRKQFNQAGGDRFGSGWVWLVRNAQNQLQIVTTANQDSPIMD----   220
MBD2168452.1         IAQQINQTFGSFDAFRKQFNEAAGDRFGSGWVWLVRNAQNQLQIVTTPNQDSPIMN----   220
MBD2342527.1         IAQQINQTFGSFDAFRKQFNEAAGDRFGSGWVWLVRNAQNQLQIVTTPNQDSPIMN----   220
MHN9116841.1         IAQQINQTFGSFDAFKKQFNEVGGDRFGSGWVWLVRNGQNQLQIVTTPNQDSPITD----   221
MFN6514993.1         IAQQINQTFGSFDAFKKQFNEAGGDRFGSGWVWLVRNAQNQLQIVNTANQDSPIME----   220
NDJ23487.1           IAQQINQTFGSFDAFKKQFNEAGGDRFGSGWVWLVRNAQNQLQIVTTPNQDNPITD----   220
WP_181929818.1       IAQQINQTFGSFDAFRKQFNEAGEGRFGSGWVWLVRNAQNQLQIVTTPNQDNPITE----   220
MBV6623186.1         IAQQINQTFGSFDAFKKQFNQAGGDRFGSGWVWLVRNPQNQLQIVTTPNQDSPIMD----   220
WP_323238745.1       IAQQINQTFGSFDAFKKQFNEEGGNRFGSGWVWLVRNPQNQLQIVTTPNQDSPIAD----   221
WP_254449390.1       IAQEINQTFGSFDAFKKQFNEAGGNRFGSGWVWLVRNPQNQLQIVTTPNQDSPIMD----   219
MBF2015390.1         IAQQINQTFGSFDNFKKQFNEAGGNRFGSGWVWLVRNAQNQLQIVNTPNQDSPILE----   219
MEB3216192.1         IAQQINQTFGSFDNFKKQFNQAGGDRFGSGWVWLVRNAQNQLQIVNTPNQDSPIID----   219
WP_193917416.1       IAQQINQTFGSFDNFKKQFNEAGGDRFGSGWVWLVRNAQNQLQIVNTPNQDSPILD----   219
WP_277875608.1       IAQEINQTFGSFEEFRKQFNQAGGDRFGSGWVWLVRNPQGQLQIVSTPNQDSPIME----   219
MBD2346042.1         IAQEINQTFGSFEEFRKQFNQAGGDRFGSGWVWLVRNPQGQLQIVSTPNQDSPIME----   206
WP_010994247.1       IAQEINQTFGSFEEFKKQFNQAGGDRFGSGWVWLVRNPQGQLQIVSTPNQDSPIME----   219
Nostoc sp. PCC 7120  IAQEINQTFGSFEEFKKQFNQAGGDRFGSGWVWLVRNPQGQLQIVSTPNQDSPIME----   219
WP_277884298.1       IAQEINQTFGNFEEFKKQFNKAGGDRFGSGWVWLVRNPQGQLQIVSTPNQDSPITE----   219
WP_414756233.1       IAQEINQTFGNFEEFKKQFNKAGGDRFGSGWVWLVRNPQGQLQIVSTPNQDSPITE----   219
WP_011318269.1       IAQEINQTFGNFEEFKKQFNKAGGDRFGSGWVWLVRNPQGQLQIVSTPNQDSPIAE----   219
WP_015215174.1       IAQEINQTFGSFEEFRNRFNQAASDRFGSGWVWLVRNPQGQLQITTTPNQDNPITE----   219
WP_276752922.1       IAQEINQSFGSFEEFRKQFNEAGGDRFGSGWVWLVRNPQGQLQITSTPNQDSPITE----   220
RUR73923.1           IAQEINQTFGSFEEFRKQFNEAGGDRFGSGWVWLVRNPQGQLQITSTPNQDSPITE----   221
WP_016872332.1       IAQEINQTFGSFEEFRKQFNEAGGDRFGSGWVWLVRNPQGQLQITSTPNQDSPITE----   220
WP_341997750.1       IAQEINQTFGSFEEFRKQFNEAGGDRFGSGWVWLVRNPQGQLQITSTPNQDSPITE----   220
WP_289789894.1       IAQEINQTFGSFEEFRNQFNEAGGDRFGSGWVWLVRNPQGQLQITSTPNQDSPITE----   219
WP_191756852.1       IAQEINQSFGSFDAFKKQFNQAGGDRFGSGWVWLVRNPQGQLQITSTPNQDSPITE----   219
WP_193198009.1       IAQEINQSFGSFDAFKKQFNQAGGDRFGSGWVWLVRNPQGQLQITSTPNQDSPITE----   219
WP_323228814.1       IAQEINQNFGSFDAFKKQFNEAGGDRFGSGWVWLVRNPQGQLQITSTPNQDSPITE----   218
WP_103138074.1       IAQEINQSFGSFDAFRKQFNEAGGDRFGSGWVWLVRNAQGQLQITSTLNQDNPISD----   219
WP_242072232.1       IAQEINQTFGSFEAFRKQFNAAGGDRFGSGWVWLVRNAQGQLQITSTPNQDNPISE----   219
MBD2440923.1         IAQEINQTFGSFEAFRKQFNAAGGDRFGSGWVWLVRNAQGQLQITSTPNQDNPISE----   204
WP_318780343.1       IAEEINKTFGNFEGFQKQFNAAGSDRFGSGWVWLVRNPQGQLQITSTPNQDSPIME----   222
WP_460206421.1       IATEINQTFGSFDAFKKQFNEAGAKQFGSGWVWLVRNPQGQLQIVSTPNQDNPIME----   223
WP_169219177.1       IAQEINQTFGSFDAFRKQFNEAGGDRFGSGWVWLVRNPQGQLQITTTPNQDSPIME----   208
WP_332997608.1       IATEINQTFGSFDAFKKQFDEAGGDRFGSGWVWLVRSPQGQLQITTTPNQDSPIME----   221
MBW4498943.1         IATEINQTFGSFDAFRKQFNEAGGDRFGSGWVWLVRNPQGQLQITTTPNQDSPIME----   208
WP_414585734.1       IATEINQTFGSFDAFKKQFDEAGGDRFGSGWVWLVRNPQGQLQITTTPNQDSPIME----   221
MCC5641606.1         IAQEINQTFGSFDTFKKQFNAAGAARFGSGWVWLVRNPQNQLQIVTTANQDNPIMD----   212
WP_435020268.1       IAQEINKTFGSFDAFKKQFNAAGAARFGSGWVWLVRNPQNQLQIVTTANQDNPIME----   214
WP_292704904.1       IAQEINQTFGSFDAFKKQFNAAGAARFGSGWVWLVRNPQNQLQIVTTANQDNPIMD----   214
WP_335036612.1       IAQEINQTFGSFDAFKKQFNAAGAARFGSGWVWLVRNPQNQLQIVTTANQDNPIMD----   214
WP_302480924.1       IAQEINQTFGSFDAFKKQFNAAGAARFGSGWVWLVRNPQNQLQIVTTANQDNPIMD----   214
WP_311774093.1       IAQEINQTFGSFDAFKKQFNAAGAARFGSGWVWLVRNPQNQLQIVTTANQDNPIMD----   214
N. punctiforme 73102 IAQEINQTFGSFDAFKKQFNAAGAARFGSGWVWLVRNPQNQLQIVTTANQDNPIMD----   214
WP_322745234.1       IAQEINQTFGSFDAFKKQFNAAGAARFGSGWVWLVRNPQNQLQIVTTANQDNPIMD----   214
WP_306297076.1       IAQEINQTFGSFDAFKKQFNAAGAARFGSGWVWLVRNPQNQLQIVTTANQDNPIMD----   214
WP_322746338.1       IAQEINQTFGSFDAFKKQFNAAGAARFGSGWVWLVRNPQNQLQIVTTANQDNPIMD----   214
WP_277883559.1       IAQEINQTFGSFDAFKKQFNAAGAARFGSGWVWLVRNPQNKLQIVTTANQDNPIMD----   214
WP_392482408.1       IAQEINQTFGSFDAFKKQFNAAGAARFGSGWVWLVRNPQNQLQIVTTANQDNPIMD----   214
WP_163931552.1       IAQEINQTFGSFDAFKKQFNAAGAARFGSGWVWLVRNPQNQLQIVTTANQDNPIMD----   214
WP_114081005.1       IAQEINQTFGSFDAFKKQFNAAGAARFGSGWVWLVRNPQNQLQIVTTANQDNPIMD----   214
                     :  :   *...  ::  *:  .  :*****.**       : :    ***.*.

E. coli              GASGFPIMGLDVWEHAYYLKFQNRRPDYIKEFWNVVNWDEAAARFAAKKIIICRLLQ      214
G. violaceus 7421     --GKYPIMGNDVWEHAYYLKYQNRRADYLSAWWNTLNWDEINRRFQKASTGL-----      270
WP_015156211.1       --DRYPIMGNDVWEHAYYLRYQNRRAEYLNNWWNVVNWEEINRRWQASSQQQ-----      268
Synechococcus 6312   --GITPLLGNDVWEHAYYLSYRNRRDQYLQAWWNVVNWDKIGERYEQATG-------      240
WP_322876523.1       --GITPLLGNDVWEHAYYLSYRNRRDQYLQAWWNVVNWDKIGERYEQATG-------      240
WP_277866136.1       --GAIPLLGNDLWEHAYYLTYRNRRDDYLQAWWNVVNWPIVEERYRRAIA-------      237
WP_011055891.1       --GQVPILGNDLWEHAYYLTYRNRRDQYLEAWWQVVNWPWVSDRYAQMRELLA----      240
WP_297761391.1       --GQVPILGNDLWEHAYYLTYRNRRDQYLEAWWQVVNWPWVSDRYAQMRALLA----      240
WP_024125507.1       --GQVPLLGNDLWEHAYYLTYRNRRDQYLEAWWQVVNWPWVSDRYAQMRDLLA----      240
WP_365970883.1       --GQVPLLGNDLWEHAYYLTYRNRRDQYLEAWWQVVNWPWVSDRYAQMRDLLA----      240
```

```
WP_409873764.1          --GQLPILGNDLWEHAYYLTYRNRRDQYLEAWWQVVNWPWLSDRYAQMRALLA----      240
WP_300872640.1          --GHVPILGNDLWEHAYYLTYRNRRDQYLEAWWQVVNWPWLSDRYAQMRALLA----      240
WP_250682089.1          --GQVPILGNDLWEHAYYLTYRNRRDQYLEAWWQVVNWPYLSDRYAQMRALLA----      240
WP_409869042.1          --GQVPILGNDLWEHAYYLTYRNRRDQYLEAWWQVVNWPWLSDRYVQMRALLA----      240
WP_297053013.1          --GQVPILGNDLWEHAYYLTYRNRRDQYLEAWWQVVNWPWLSDRYAQMRALLA----      240
WP_297051732.1          --GQVPILGNDLWEHAYYLTYRNRRDQYLEAWWQVVNWPWLSDRYAQMRALLA----      240
WP_181494764.1          --GQVPILGNDLWEHAYYLTYRNRRDQYLEAWWQVVNWPWLSDRYAQMRALLA----      240
WP_460193008.1          --GQVPILGNDLWEHAYYLTYRNRRDQYLEAWWQVVNWPWLSDRYAQTRALLA----      240
WP_286627080.1          --GQVPILGNDLWEHAYYLTYRNRRDQYLEAWWQVVNWPWLSDRYAQMRALLT----      240
WP_310743007.1          --GQVPILGNDLWEHAYYLTYRNRRDQYLEAWWQVVNWPWLSDRYAQMRALLT----      240
WP_206200084.1          --GQVPILGNDLWEHAYYLTYRNRRDQYLEAWWQVVNWPWLSDRYAQMRALLT----      240
WP_310821562.1          --GQVPILGNDLWEHAYYLTYRNRRDQYLEAWWQVVNWPWLSDRYAQMRALLT----      240
WP_310800931.1          --GQVPILGNDLWEHAYYLTYRNRRDQYLEAWWQVVNWPWLSDRYAQMRALLT----      240
WP_310454098.1          --GQVPILGNDLWEHAYYLTYRNRRDQYLEAWWQVVNWPWLSDRYAQMRALLT----      240
WP_302677659.1          --GQVPILGNDLWEHAYYLTYRNRRDQYLEAWWQVVNWPWLSDRYAQMRALLT----      240
Pseudanabaena 7367      --GLFPIMGNDVWEHAYYLNYQNRRGDYLKAWWNLVNWEEVNNRFLNAMV-------      254
MCU0526905.1            --GYYPIMGNDVWEHAYYLRYRNKRDEYLKAWWNVLNWEAIAQRYEAAVQV------      256
MDJ0592545.1            --GMYPIMGNDVWEHAYYLKYRNKRGEYLENWWNVVNWQEVTQRYEQAIA-------      239
MBX5479872.1            --GLYPIMGNDVWEHAYYLKYRNRRNEYLEAWWNVVNWDEINRRFETATSQRTTRRT      251
WP_041973980.1          --GLYPIMGNDVWEHAYYLKYQNRRNEYLEAWWNVVNWDEINRRFETATSQRTTRRT      251
WP_428360810.1          --GRYPILGNDVWEHAYYLKYQNRRADYLNAWWNVVNWEAVNKRFSQVRA-------      239
WP_233744687.1          --GRYPILGNDVWEHAYYLKYQNRRADYLNAWWNVVNWEAVNQRLRQAKVSIG----      242
WP_445926464.1          --GRYPILGNDVWEHAYYLKYQNRRTDYLNAWWNVVNWEAVNQRLRQAKASIG----      242
WP_015152615.1          --GNYPIMGNDVWEHAYYLKYRNRRAEYLKAWWNVVNWQEINQRLALAAKAA-----      244
WP_190455393.1          --GMYPIMGNDVWEHAYYLKYQNKRADYLNAWWNVVNWNEVNKRFERASA-------      254
WP_190424133.1          --GMYPIMGNDVWEHAYYLKYQNKRADYLNAWWNVVNWNEVNKRFERASA-------      254
WP_027843320.1          --GKYPIMGNDVWEHAYYIKYRNRRAEYLKAWWNVLNWDEINKRFAKAVA-------      251
WP_374791410.1          --GAYPIMGNDVWEHAYYLKYQNRRAEYLDAWWNVLNWDEINKRLQQANA-------      254
WP_374825979.1          --GAYPIMGNDVWEHAYYLKYQNRRAEYLDAWWNVLNWDEINKRLQQANA-------      254
WP_190465982.1          --GAYPIMGNDVWEHAYYLKYQNRRAEYLDAWWNVLNWDEINKRLQQANG-------      254
BDA72795.1              --GKYPIMGNDVWEHAYYLKYQNQRAKYLEAWWNVLNWDEINKRFEQASKVA-----      257
GJD16125.1              --GKYPIMGNDVWEHAYYLKYQNKRADYLEAWWNVLNWDEINKRFEQANKVA-----      257
MEC4881204.1            --GQYPIMGNDVWEHAYYLKYQNRRADYLDAWWNVLNWEEINRRLVEGGRE------      246
WP_190560307.1          --GNYPIFGNDVWEHAYYLKYQNRRPDYLEAWWNVVNWDEINQRFADASKIG-----      240
MFM5980434.1            --GKYPIFGNDVWEHAYYLKYQNRRPEYLEAWWNVVNWDEINQRFADASKIE-----      240
WP_096571931.1          --GKYPIFGNDVWEHAYYLKYQNRRPEYLDAWWNVVNWDEINQRFADASKIE-----      240
WP_190648403.1          --GKYPIFGNDVWEHAYYLKYQNRRPEYLEAWWNVVNWDEINQRFADASKIE-----      240
WP_190347369.1          --GKYPIFGNDVWEHAYYLKYQNRRPEYLEAWWNVVNWDEINQRFADASKIE-----      240
N. punctiforme 73102--GKYPILGNDVWEHAYYLNYQNRRADYLDAWWNVVNWDEINKRFAAANKFA-----      259
WP_334931430.1          --GKYPILGNDVWEHAYYLNYQNRRADYLDAWWNVVNWDEINKRFAAASKFA-----      259
WP_335056299.1          --GKYPILGNDVWEHAYYLNYQNRRADYLDAWWNVVNWDEINKRFAAASKFA-----      259
WP_322673311.1          --GKYPILGNDVWEHAYYLNYQNRRADYLDAWWNVVNWDEINKRFAAASKFA-----      259
WP_322689216.1          --GKYPILGNDVWEHAYYLNYQNRRADYLDAWWNVVNWDEINKRFAAASKFA-----      259
WP_322701324.1          --GKYPILGNDVWEHAYYLNYQNRRADYLDAWWNVVNWDEINKRFAAASKFA-----      259
WP_335012934.1          --GKYPILGNDVWEHAYYLNYQNRRADYLDAWWNVVNWDEINKRFAAASKFA-----      255
WP_292868329.1          --GKYPILGNDVWEHAYYLNYQNRRADYLDAWWNVVNWDEINKRFAAASKFA-----      267
WP_334926809.1          --GKYPILGNDVWEHAYYLNYQNRRADYLDAWWNVVNWDEINKRFAAASKFA-----      267
WP_109007648.1          --GKYPILGNDVWEHAYYLNYQNRRADYLDAWWNVVNWDEINKRFAAATKFA-----      255
WP_289795776.1          --GKYPILGNDVWEHAYYLNYQNRRADYLDAWWNVVNWDEINKRFAAASKFA-----      255
WP_099100092.1          --GKYPILGNDVWEHAYYLNYQNRRADYLDAWWNVVNWDEINKRFAAASKFA-----      255
WP_335201774.1          --GKYPILGNDVWEHAYYLNYQNRRADYLDAWWNVVNWDEINKRFAAASKLA-----      259
MDZ7959988.1            --GKYPILGNDVWEHAYYLNYQNRRADYLEAWWNVVNWDEINKRFAAATKFA-----      252
MBD2208974.1            --GKYPILGNDVWEHAYYLNYQNRRADYLEAWWNVVNWDEINKRFATASKFA-----      251
MBD2339284.1            --GKYPILGNDVWEHAYYLNYQNRRADYLEAWWNVVNWDEINKRFAAASKFA-----      251
MBD2166265.1            --GKYPILGNDVWEHAYYLNYQNRRADYLEAWWNVVNWDEINKRFAAASKFA-----      251
WP_167724335.1          --GKYPILGNDVWEHAYYLNYQNRRADYLEAWWNVVNWDEINKRFAAASKFA-----      251
WP_045869977.1          --GKYPILGNDVWEHAYYLNYQNRRADYLEAWWNVVNWDEINQRFAAASKFA-----      252
WP_096582728.1          --GKYPILGNDVWEHAYYLNYQNRRADYLEAWWNVVNWDEINQRFATASKFA-----      251
WP_096599871.1          --GLYPILGNDVWEHAYYLNYQNRRADYLEAWWNVVNWDEINKRYAVVSKSA-----      252
WP_190743315.1          --GLYPILGNDVWEHAYYLNYQNRRADYLEAWWNVLNWDEINKRFADASKLA-----      252
WP_190544005.1          --GLYPILGNDVWEHAYYLNYQNRRADYLEAWWNVLNWDEINQRFAAASKSA-----      251
WP_096643960.1          --GLYPILGNDVWEHAYYLNYQNRRADYLEAWWNVLNWDEINQRFAAASKSA-----      251
Cyanothece 7425         --SNYPIMGNDVWEHAYYLRYQNQRAEYLNNWWNVVNWGEIDRRAQGSQLMLRSN--      268
WP_414578860.1          --GLYPIMGNDVWEHAYYLRYRNRRPEYLNNWWNVVNWTEIDKRAAIR---------      256
WP_323267495.1          --GLYPIMGNDVWEHAYYLRYRNRRPEYLNNWWNVVNWAEIDKRAAIR---------      256
WP_214431594.1          --GLYPILGNDVWEHAYYLRYRNRRPEYLNNWWNVVNWSEINKRNQISSAK------      265
WP_322662701.1          --GLYPILGNDVWEHAYYLRYRNRRPEYLNNWWNVVNWSEINRRNQISSTK------      266
RCJ28434.1              --GLYPILGNDVWEHAYYLRYRNRRPEYLNNWWNVVNWSEINRRNQISSTK------      266
MBE9051467.1            --GAYPIMGNDVWEHAYYLRYRNRRAEYLNNWWNVVNWPEINRRSQVSLQLAS----      264
MBW4558831.1            --GAYPIMGNDVWEHAYYLRYRNRRGEYLNNWWNVVNWPEINRRSQVSLQLAS----      248
WP_063871086.1          --GAYPIMGNDVWEHAYYLRYRNRRAEYLKNWWNVVNWPEVNKRSLSGE--------      260
WP_006196478.1          --GAYPIMGNDVWEHAYYLRYRNRRAEYLKNWWNVVNWPEVNKRSLVNLQKGT----      264
WP_271918904.1          --GAYPIMGNDVWEHAYYLRYRNRRAEYLNNWWNVVNWPEINKRSLVSLQ-------      261
WP_271942563.1          --GAYPIMGNDVWEHAYYLRYRNRRAEYLNNWWNVVNWPEINKRSLVNLQKGT----      264
WP_272068383.1          --GAYPIMGNDVWEHAYYLRYRNRRAEYLTNWWNVVNWPEINRRSLVSLQSAT----      264
WP_193998321.1          --GAYPIMGNDVWEHAYYLRYRNRRAEYLNNWWNVVNWPEINKRSLVSLQSGS----      263
WP_089093562.1          --GAYPIMGNDVWEHAYYLRYRNRRGEYLTNWWNVVNWPEINKRSLASLQSGR----      264
```

```
WP_323215458.1       --GAYPIMGNDVWEHAYYLRYRNRRAEYLNNWWNVVNWPEINKRSLVSLQSGS----     264
WP_414550770.1       --GAYPIMGNDVWEHAYYLRYRNRRAEYLNNWWNVVNWPEINKRSLISLQ-------     261
WP_214442418.1       --GVYPIMGNDVWEHAYYLRYRNRRSEYLNNWWDVVNWQEINKRSQLSLQQPR----     264
MEA5507287.1         --GAYPIMGNDVWEHAYYLRYRNRRGEYLNNWWNVVNWPEINKRSQLSSQQPR----     264
WP_281484005.1       --GAYPIMGNDVWEHAYYLRYRNRRGEYLDNWWNVVNWPEIDKRSQLSLQQRG----     266
WP_162421941.1       --GSYPIMGNDVWEHAYYLRYQNQRAQYLNNWWNVVNWDEVNRRTQASQPR------     262
WP_430678559.1       --GSYPIMGNDVWEHAYYLRYQNRRAEYLNNWWNVVNWDEINRRFQAAQQGGA----     263
MBE9012496.1         --GSYPIMGNDVWEHAYYLRYQNRRADYLNNWWNVVNWGEINRRYQASQQR------     261
WP_059000726.1       --GSYPIMGNDVWEHAYYLRYRNRRADYLNNWWNVVNWSEINRRLQASQQSS-----     262
WP_482030890.1       --GNYPIMGNDVWEHAYYLRYQNRRAEYLSNWWNVVNWTEVNRRFQLASQQGS----     254
WP_445299659.1       --GLYPILGNDVWEHAYYLRYQNRRAEYLTNWWNVVNWEEVNRRFQLASQP------     266
WP_442927815.1       --GLYPILGNDVWEHAYYLRYQNRRAEYLTNWWNVVNWEEVNRRFQLASQP------     266
WP_445299397.1       --GLYPILGNDVWEHAYYLRYQNRRAEYLTNWWDVVNWEEVNRRFQLASQP------     260
WP_445312326.1       --GLYPILGNDVWEHAYYLRYQNRRAEYLTNWWNVVNWEEVNRRFQLASQP------     260
HIK16311.1           --GNFPIMGNDVWEHAYYLRYQNRRAEYLNNWWNVVNWQEVNRRFQLAAQQ------     257
WP_432812051.1       --GNFPLLGNDVWEHAYYLRYQNRRAEYLSNWWNVVNWQEVNRRWQRASQR------     257
WP_431655370.1       --GNFPIMGNDVWEHAYYLRYQNRRAEYLNNWWNVVNWQEVNRRWQLASQQ------     258
MGI0483888.1         --GNFPIMGNDVWEHAYYLRYQNRRAEYLNNWWNVVNWQEVNRRWQLASQQ------     243
MBW4417879.1         --GNFPIMGNDVWEHAYYLKYRNRRAEYLSNWWNVVNWAEVDRRAKVRLGTASS---     261
MBW4540880.1         --GNFPIMGNDVWEHAYYLKYRNRRAEYLNNWWNVVNWAEVDRRAKVRLGTANS---     261
MBW4644872.1         --GLYPIMGNDVWEHAYYLRYQNRRAEYLNNWWNVVNWPEINRRLQAASRQQS----     270
PMB52031.1           --GLYPIMGNDVWEHAYYLRYQNRRAEYLNNWWNVVNWSEINRRVQAVSQQQG----     271
WP_190751173.1       --GLYPIMGNDVWEHAYYLRYQNRRPEYLNNWWNVVNWTEINRRSQAGLQQKV----     271
WP_190549236.1       --GLYPIMGNDVWEHAYYLRYQNRRPEYLNNWWNVVNWTEINRRSQAGLQQKV----     271
WP_190629649.1       --GLYPIMGNDVWEHAYYLRYQNRRPEYLNNWWNVVNWTEINRRSQAGLQQKV----     271
MBD2168452.1         --GLYPIMGNDVWEHAYYLRYQNRRPEYLNNWWNVVNWSEINRRSQAALQQKA----     271
MBD2342527.1         --GLYPIMGNDVWEHAYYLRYQNRRPEYLNNWWNVVNWSEINRRSQAALQQKT----     271
MHN9116841.1         --GLYPIMGNDVWEHAYYLRYQNRRPEYLNNWWNVVNWSEINRRSQVKSQQG-----     271
MFN6514993.1         --GLYPIMGNDVWEHAYYIRYQNRRAEYLNNWWNVVNWSQINRRVQAVSQQKN----     271
NDJ23487.1           --GLYPIMGNDVWEHAYYLRYQNRRAEYLDNWWNVVNWSEINRRMQVKSQQS-----     270
WP_181929818.1       --GLYPIMGNDIWEHAYYLRYQNRRAEYLNNWWNVVNWSEINTRAQVKSQQS-----     270
MBV6623186.1         --SLYPIMGNDVWEHAYYLRYQNQRAEYLNNWWNVVNWSEINRRIQAVSQQDDT---     272
WP_323238745.1       --GLYPIMGNDVWEHAYYLRYQNRRAEYLNNWWNVVNWSEINRRLQSVSQPA-----     271
WP_254449390.1       --GLYPIMGNDVWEHAYYLRYQNRRAEYLNNWWNVVNWTEINRRVQAVSSQ------     268
MBF2015390.1         --GLYPIMGNDVWEHAYYLRYQNRRAEYLNNWWNVVNWSEINRRAQVVSQ-------     267
MEB3216192.1         --GLYPIMGNDVWEHAYYLRYQNRRAEYLENWWNVVNWSEINRRAQVVSQQQS----     270
WP_193917416.1       --GLYPIMGNDVWEHAYYLRYQNRRAEYLDNWWNVVNWSEINRRAQVVSQQG-----     269
WP_277875608.1       --GSYPIMGNDVWEHAYYLRYQNRRADYLNNWWDVVNWTEINRRAQASQQNNS----     270
MBD2346042.1         --GSYPIMGNDVWEHAYYLRYQNRRADYLNNWWDVVNWTEINRRAQASQQNNS----     257
WP_010994247.1       --GSYPIMGNDVWEHAYYLRYQNRRPEYLNNWWNVVNWSEINRRTQASRQSNS----     270
Nostoc sp. PCC 7120 --GSYPIMGNDVWEHAYYLRYQNRRPEYLNNWWNVVNWSEINRRTQASRQSNS----     270
WP_277884298.1       --GSYPIMGNDVWEHAYYLRYQNRRPEYLNNWWNVVNWTEINKRAQASRQSNS----     270
WP_414756233.1       --GSYPIMGNDVWEHAYYLRYQNRRPEYLNNWWNVVNWSEINKRAQASRQSNS----     270
WP_011318269.1       --GSYPIMGNDVWEHAYYLRYQNRRPEYLNNWWNVVNWSEINKRAQASRQSNS----     270
WP_015215174.1       --GSYPIMGNDVWEHAYYLRYQNRRADYLNNWWNVVNWTEINKRAQASRQQA-----     269
WP_276752922.1       --GSYPIMGNDVWEHAYYLRYQNRRAEYLNNWWNVVNWQEINRRAQASRQQS-----     270
RUR73923.1           --GSYPIMGNDVWEHAYYLRYQNRRAEYLNNWWNVVNWQEINRRAQASRQQS-----     271
WP_016872332.1       --GSYPIMGNDVWEHAYYLRYQNRRAEYLNNWWNVVNWQEINRRAQASRQQS-----     270
WP_341997750.1       --GSYPIMGNDVWEHAYYLRYQNRRAEYLNNWWNVVNWQEINRRAQASRQQS-----     270
WP_289789894.1       --GSYPIMGNDVWEHAYYLRYQNRRAEYLNNWWNVVNWQEINRRAQASRQQS-----     269
WP_191756852.1       --GLFPIMGNDVWEHAYYLRYQNRRADYLNNWWNVVNWTEINRRAEASRQKT-----     269
WP_193198009.1       --GLFPIMGNDVWEHAYYLRYQNRRADYLNNWWNVVNWPEINRRAEASRQNT-----     269
WP_323228814.1       --GLYPIMGNDVWEHAYYLRYQNRRADYLNNWWNVVNWSEVNRRSQASRQQT-----     268
WP_103138074.1       --GAYPIMGNDVWEHAYYLRYQNRRADYLNNWWNVVNWTEVNRRAQASRQNS-----     269
WP_242072232.1       --GSYPIMGNDVWEHAYYLRYQNRRADYLNNWWNVVNWTEINRRAQASRQKT-----     269
MBD2440923.1         --GSYPIMGNDVWEHAYYLRYQNRRADYLNNWWNVVNWTEINRRAQASRQKT-----     254
WP_318780343.1       --GLYPIMGNDVWEHAYYLRYQNRRADYLNNWWNVVNWSEINRRAQASLKQPA----     273
WP_460206421.1       --DSYPIMGNDVWEHAYYLRYRNRRAEYLDNWWNVVNWSEVNRRAQVASQKQA----     274
WP_169219177.1       --GLYPIMGNDVWEHAYYLRYQNQRAKYLENWWNVVNWSEINRRAQVSSQRQS----     259
WP_332997608.1       --GLYPIMGNDVWEHAYYLRYQNRRAEYLENWWNVVNWSEVNKRAQVSSQRQS----     272
MBW4498943.1         --GLYPIMGNDVWEHAYYLRYQNRRAEYLENWWNVVNWSEVNRRAQVSSQRQS----     259
WP_414585734.1       --GLYPIMGNDVWEHAYYLRYQNRRAEYLENWWNVVNWSEVNRRAQVSSQRQS----     272
MCC5641606.1         --GSYPIMGNDLWEHAYYLKYRNRRGEYLDNWWNVVNWPQINRRSQLSSQQPR----     263
WP_435020268.1       --DSYPIMGNDLWEHAYYLKYRNRRAEYLNNWWNVVNWPQINRRYLSSSTERSVN--     267
WP_292704904.1       --GFYPIMGNDLWEHAYYLKYRNRRAEYLNNWWNVVNWPQINRRSQLSPKQPR----     265
WP_335036612.1       --GFYPIMGNDLWEHAYYLKYRNRRAEYLNNWWNVVNWPQINRRYLR----------     259
WP_302480924.1       --GFYPIMGNDLWEHAYYLKYRNRRAEYLNNWWNVVNWPQINRRSQLSSQQPR----     265
WP_311774093.1       --GFYPIMGNDLWEHAYYLKYRNRRAEYLNNWWNVVNWPQINRRSQLSSQQPR----     265
N. punctiforme 73102--GFYPIMGNDLWEHAYYLKYRNRRAEYLNNWWNVVNWPQINRRSQLSSQQPR----     265
WP_322745234.1       --GFYPIMGNDLWEHAYYLKYRNRRAEYLNNWWNVVNWPQINRRSQLSSQQPR----     265
WP_306297076.1       --GFYPIMGNDLWEHAYYLKYRNRRAEYLNNWWNVVNWPQINRRWQLSSQQPR----     265
WP_322746338.1       --GFYPIMGNDLWEHAYYLKYRNRRAEYLNNWWNVVNWPQINRRSQLSSQQPR----     265
WP_277883559.1       --GFYPIMGNDLWEHAYYLKYRNRRAEYLNNWWNVVNWPQINRRSQLSSQQPR----     265
WP_392482408.1       --GFYPIMGNDLWEHAYYLKYRNRRAEYLNNWWNVVNWPQINRRSQLSSQQPR----     265
WP_163931552.1       --GFYPIMGNDLWEHAYYLKYRNRRAEYLNNWWNVVNWPQINRRSQLSSQQPR----     265
WP_114081005.1       --GFYPIMGNDLWEHAYYLKYRNRRAEYLNNWWNVVNWPQINRRSQLSSQQPR----     265
                       .  *::* *:******: ::*:* .*:  :*: :**     *
```

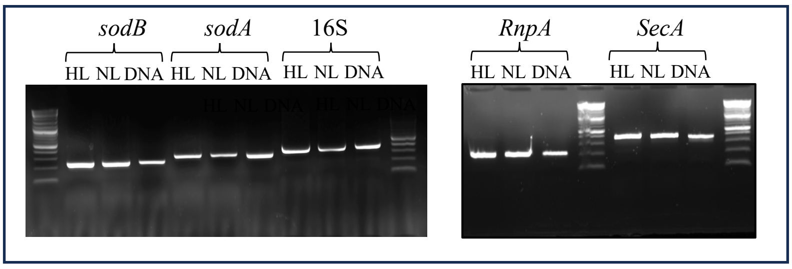

**Fig. S5:** Semi-quantitative RT-PCR showing the expression under different light intensities of both *sodA* and *sodB*. *rnpA*, *secA* and 16S were used as housekeeping controls.

| Experiment | Protein | Coverage | TotalPeptides | UniquePeptides | RazorPeptides | TotalSpectralCo... | UniqueSpectral... | RazorSpectralC... |
|---|---|---|---|---|---|---|---|---|
| Total | Cyan7425_3014\|ID:3828554\| | 14,55 | 3 | 3 | 3 | 4 | 4 | |
| Membrane | Cyan7425_3014\|ID:3828554\| | 14,55 | 3 | 3 | 3 | 4 | 4 | |

TOTAL

Cyan7425_3014|ID:3828554|

M N S S R S I S H Y L R Q L V V L L L T F V L A G L V I A C Q P T P T T
Q A P S P T P T V S P V P V A S P T G E L R A Y P A E L P P L P Y D Y G
A L E T A I D A R T M E I H H D R H H A A Y V K N L N A A L Q Q D T N L
Q N Q S V E A L L R D L N S V P E A I R E R V R N N G G G H L N H T I F
W Q I M G P D G G G E P T G D L A Q E I N Q T F G S F E E L R K Q F N Q
A G G D R F G S G W V W L V R N P Q G Q L Q I V T T A N Q D N P I T D S
N Y P I M G N D V W E H A Y Y L R Y Q N Q R A E Y L N N W W N V V N W G
E I D R R A Q G S Q L M L R S N

MEMBRANE

Cyan7425_3014|ID:3828554|

M N S S R S I S H Y L R Q L V V L L L T F V L A G L V I A C Q P T P T T
Q A P S P T P T V S P V P V A S P T G E L R A Y P A E L P P L P Y D Y G
A L E T A I D A R T M E I H H D R H H A A Y V K N L N A A L Q Q D T N L
Q N Q S V E A L L R D L N S V P E A I R E R V R N N G G G H L N H T I F
W Q I M G P D G G G E P T G D L A Q E I N Q T F G S F E E L R K Q F N Q
A G G D R F G S G W V W L V R N P Q G Q L Q I V T T A N Q D N P I T D S
N Y P I M G N D V W E H A Y Y L R Y Q N Q R A E Y L N N W W N V V N W G
E I D R R A Q G S Q L M L R S N

| Experiment | Sequence | Charge | Retenti... | Observ... | Observ... | Calcula... |
|---|---|---|---|---|---|---|
| Total | NH2-HHAAYVK-COOH | 2 | 193,811 | 824,424 | 413,219 | 824,429 |
| Total | NH2-NLNAALQQDTNLQNQSVEALLR-COOH | 2 | 2 515,679 | 2 452,253 | 1 227,134 | 2 452,267 |
| Total | NH2-NLNAALQQDTNLQNQSVEALLR-COOH | 2 | 2 690,334 | 2 454,226 | 1 228,12 | 2 452,267 |
| Total | NH2-FGSGWVWLVR-COOH | 2 | 2 733,997 | 1 205,631 | 603,823 | 1 205,634 |
| Membrane | NH2-HHAAYVK-COOH | 2 | 159,508 | 824,427 | 413,221 | 824,429 |
| Membrane | NH2-NLNAALQQDTNLQNQSVEALLR-COOH | 2 | 2 525 | 2 452,26 | 1 227,138 | 2 452,267 |
| Membrane | NH2-NLNAALQQDTNLQNQSVEALLR-COOH | 2 | 2 526,898 | 2 452,26 | 1 227,137 | 2 452,267 |
| Membrane | NH2-FGSGWVWLVR-COOH | 2 | 2 737,62 | 1 205,632 | 603,823 | 1 205,634 |

| Experiment | DeltaM... | Exp... | Hyper... | Next... | Prob... | Protein... | Protein... | Protein |
|---|---|---|---|---|---|---|---|---|
| Total | -0,003 | 0,005 | 15,833 | 0 | 0.9957 | 90 | 96 | Cyan7425_3014\|ID:382855 |
| Total | -0,012 | 0 | 33,403 | 9,416 | 1.0 | 97 | 118 | Cyan7425_3014\|ID:382855 |
| Total | 1,961 | 0,146 | 12,844 | 8,33 | 0.9191 | 97 | 118 | Cyan7425_3014\|ID:382855 |
| Total | -0 | 0,001 | 20,322 | 0 | 1.0 | 186 | 195 | Cyan7425_3014\|ID:382855 |
| Membrane | -0,002 | 0,001 | 17,117 | 0 | 0.9993 | 90 | 96 | Cyan7425_3014\|ID:382855 |
| Membrane | -0,006 | 0 | 36,575 | 9,51 | 1.0 | 97 | 118 | Cyan7425_3014\|ID:382855 |
| Membrane | -0,006 | 0 | 29,258 | 7,924 | 1.0 | 97 | 118 | Cyan7425_3014\|ID:382855 |
| Membrane | 0 | 0 | 22,574 | 0 | 1.0 | 186 | 195 | Cyan7425_3014\|ID:382855 |

**Fig. S6**: Mass spectrometry identification of the MnSodA (Cyan7425_3014) in total protein fraction and in isolated membrane fraction of *Cyanothece* PCC 7425.

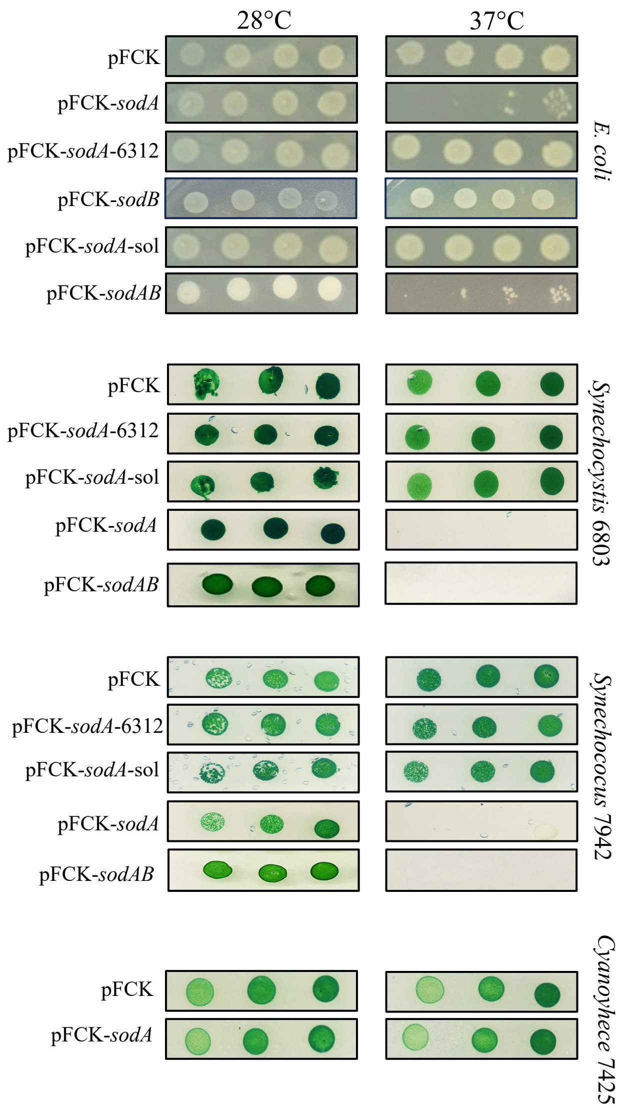

**Fig. S7**: Effect of expression of the MnSodA or its N-ter from *Cyanthece* PCC 7425 in *E. coli* and model cyanobacteria.

Alignement SodB/ SodAB

```
SodB        -----------------------------------------------------------M      1
SodAB       MNSSRSISHYLRQLVVLLLTFVLAGLVIACQPTPTTQAPSPTPTVSPVPVASPTGELRAY      60

SodB        AFEQPPLPYDPGALEPYGMSARTFEFHYGKHHAAYVTNLNNLTKDTELADKSLEEVIKVT      61
SodAB       AFEQPPLPYDPGALEPYGMSARTFEFHYGKHHAAYVTNLNNLTKDTELADKSLEEVIKVT      120
            ************************************************************

SodB        FKDPGKVGVFNNAAQVWNHTFFWNSLKPGGGGTPSGDLAAKIDSAFGSFDKFKEEFKTAA      121
SodAB       FKDPGKVGVFNNAAQVWNHTFFWNSLKPGGGGTPSGDLAAKIDSAFGSFDKFKEEFKTAA      180
            ************************************************************

SodB        ATQFGSGWAWLVEDGGSLKVIKTPNAENPLAYGQKSLLTLDVWEHAYYIDYQNRRPDFIQ      181
SodAB       ATQFGSGWAWLVEDGGSLKVIKTPNAENPLAYGQKSLLTLDVWEHAYYIDYQNRRPDFIQ      240
            ************************************************************

SodB        NFLDQLVNWDFVAQNLAA*    199
SodAB       NFLDQLVNWDFVAQNLAA*    258
            ******************
```

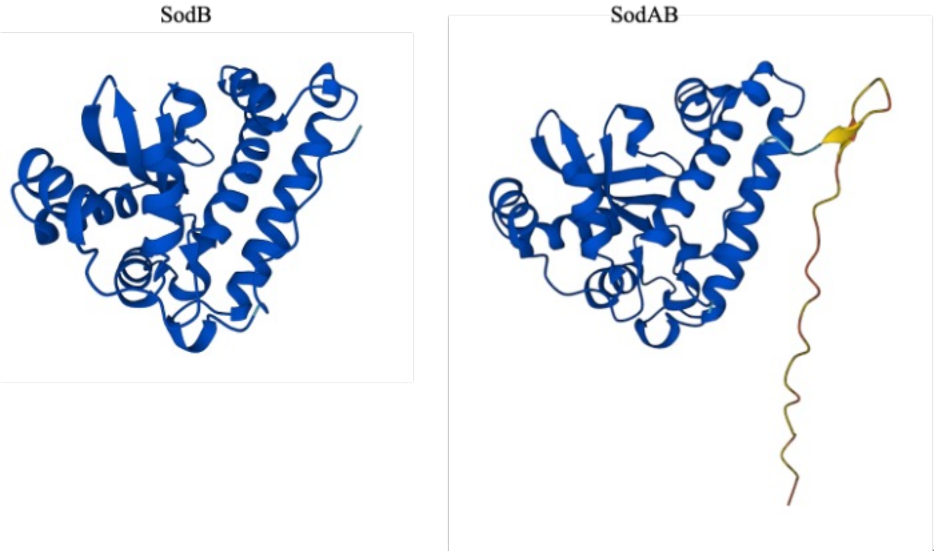

**Fig. S8**: Sequence alignment and structural prediction of the fused SodAB protein. SodAB was generated by fusion of the N-terminal region of MnSodA with the FeSodB

**MnSOD Diversifies, while FeSOD Dominates Superoxide Defense in Cyanobacteria**

Monis Athar KHAN, Anne DURAND, Anne-Soisig STEUNOU, Solène LELEUX, Corinne CASSIER-CHAUVAT, Franck CHAUVAT, Soufian OUCHANE*

### Supplementary material
### (Tables)

**Table S1:** Strains and plasmids used in this study

| Strains | Relevant features | References |
|---|---|---|
| ***E. coli*** | | |
| NEB10-beta | Δ(*ara-leu*) 769 7ara D139 *fhuA* Δ*lac*X74 *galK16 galE15 e14*-ϕ80d*lacZ*ΔM15 *recA1 relA1 endA1 nupG rpsL* ($Str^R$) rph *spoT1* Δ(*mrr-hsdRMS-mcrBC*) | New England Biolabs |
| NEB10-beta/pSL2680 | NEB10-beta strain containing the pSL2680 plasmid | [1] |
| NEB10-beta/pSL-ΔsodA | NEB10-beta strain containing the pSL-ΔsodA plasmid | This study |
| NEB10-beta/pSL-ΔsodB | NEB10-beta strain containing the pSL-ΔsodB plasmid | [1] |
| NEB10-beta/pFCK | NEB10-beta strain containing the pFCK plasmid | This study |
| NEB10-beta/pFCK-*sodA*-6312 | NEB10-beta strain containing the pFCK-*sodA*-6312plasmid | This study |
| NEB10-beta/pFCK-*sodA* | NEB10-beta strain containing the pFCK-*sodA* plasmid | This study |
| NEB10-beta/pFCK-*sodA*-sol | NEB10-beta strain containing the pFCK-*sodA*-sol plasmid | This study |
| NEB10-beta/pFCK-*sodB* | NEB10-beta strain containing the pFCK-*sodB* plasmid | This study |
| NEB10-beta/pFCK-*sodAB* | NEB10-beta strain containing the pFCK-*sodAB* plasmid | This study |
| AM5501 | Donor strain containing both pRL443 and pAM5505 plasmids for conjugation | [2] |
| CM404 | Donor strain containing the self-transmissible plasmid pRK2013 for conjugative transfer of RSF1010 plasmids | [3] |
| **Cyanobacteria** | | |
| *Cyanothece* PCC 7425 | | Pasteur institute (Paris, France) |
| *Cyanothece* PCC 7425-ΔsodA | | This study |
| *Cyanothece* PCC 7425-ΔsodB | | [1] |
| *Synechocystis* sp. PCC 6803 | | Pasteur institute (Paris, France) |
| *Synechococcus* PCC 7942 | | Pasteur institute (Paris, France) |
| *Synechococcus* PCC 6312 | | Pasteur institute (Paris, France) |

| **Plasmids** | | |
|---|---|---|
| pC | RSF1010 derivative ($SpSm^R$, $Cm^R$) harboring the λ phage promoter allowing strong constitutive expression. | [4] |
| pFCK | RSF1010 derivative ($Km^R$, $Cm^R$) harboring the λ phage promoter allowing strong temperature regulated expression. | [5] |
| pSL2680 | RSF1010 derivative ($Km^R$) harboring the cas12a gene that is expressed from a lac promoter and the endogenous CRISPR array from *Francisella novicida* from a J23119 promoter. | [6] |
| pSL-ΔsodA | pSL2680 vector ($Km^R$, $SpSm^R$) with crRNA targeting *sodA,* and $SpSm^R$ cassette flanking homology platforms (~600 bp) for deletion of *sodA* gene | This study |
| pSL-ΔsodB | pSL2680 vector ($Km^R$, $SpSm^R$) with crRNA targeting *sodB,* and $Sm^R$/$Sp^R$ cassette flanking homology platforms (~500 bp) for deletion of *sodB* gene | [1] |

| | | |
|---|---|---|
| pFCK-*sodA* | Expression vector pFCK plasmid harboring the *Cyanothece* 7425 *sodA* gene cloned at the NdeI/ EcoRI site. | This study |
| pFCK-*sodA-6312* | Expression vector pFCK plasmid harboring *Synechococcus* PCC 6312 *sodA* gene cloned at the NdeI/ EcoRI site. | This study |
| pFCK-*sodA-sol* | Expression vector pFCK plasmid harboring *Cyanothece* 7425 *sodA* truncated gene cloned at the NdeI/ EcoRI site. | This study |
| pFCK-*sodB* | Expression vector pFCK plasmid harboring the *Cyanothece* 7425 *sodB* gene cloned at the NdeI/ EcoRI site. | This study |
| pFCK-*sodAB* | Expression vector pFCK plasmid harboring the chimeric *sodAB* gene cloned at the NdeI/ EcoRI site. | This study |

1. Khan MA, Durand A, Skouri-Panet F, Benzerara K, Cassier-Chauvat C, Chauvat F, et al. Targeted genome editing of the non-model cyanobacterium Cyanothece PCC 7425 via CRISPR/Cas12a. Appl Microbiol Biotechnol. 2026 [cited 27 Sept 2026]. doi:10.1007/s00253-026-13959-y

2. Modification of RSF1010-Based Broad-Host-Range Plasmids for Improved Conjugation and Cyanobacterial Bioprospecting: iScience. [cited 1 Oct 2026]. Available: https://www-cell-com.insb.bib.cnrs.fr/iscience/fulltext/S2589-0042(19)30339-6

3. Marraccini P, Bulteau S, Cassier-Chauvat C, Mermet-Bouvier P, Chauvat F. A conjugative plasmid vector for promoter analysis in several cyanobacteria of the genera Synechococcus and Synechocystis. Plant Mol Biol. 1993;23: 905–909. doi:10.1007/BF00021546

4. Chenebault C, Diaz-Santos E, Kammerscheit X, Görgen S, Ilioaia C, Streckaite S, et al. A Genetic Toolbox for the New Model Cyanobacterium Cyanothece PCC 7425: A Case Study for the Photosynthetic Production of Limonene. Front Microbiol. 2020;11: 586601. doi:10.3389/fmicb.2020.586601

5. Marceau F, Lamothe-Sibold M, Farci S, Ouchane S, Cassier-Chauvat C, Chauvat F. First Characterization of a Cyanobacterial Xi-Class Glutathione S-Transferase in Synechocystis PCC 6803. Antioxidants. 2024;13: 1577. doi:10.3390/antiox13121577

6. Ungerer J, Pakrasi HB. Cpf1 Is A Versatile Tool for CRISPR Genome Editing Across Diverse Species of Cyanobacteria. Sci Rep. 2016;6: 39681. doi:10.1038/srep39681

**Table S2 :** primers used in this study

| **Primer** | **5'-3' Sequence** |
|---|---|
| cr-sodA_Fw | agatAATAGTGTGCCAGAAGCCATTC |
| cr-sodA_Rv | agacGAATGGCTTCTGGCACACTATT |
| F1-sodA_Fw | CGGTAGTTGGTACCGTCGACGCCCAACTGCTGCATCTGGA |
| F1-sodA_Rv | ACAGGCTTATGAATAACATGGTTAAATTTAATTAATTTCTACCCCGG |
| F2-sodA_Fw | CCATGTTATTCATAAGCCTGTTCGGTTCGTAAGC |
| F2-sodA_Rv | CCCTACCCCTTTAATGCGGATGTTGCGATTACTTCG |
| F3-sodA_Fw | GCATTAAAGGGGTAGGGTGACCGT |
| F3-sodA_Rv | CGGATTACAGATCCTCTAGAGTCGACTCGATCAAACCCTGCAGCTG |
| sodA-Fw | CTGTCTGCATCTGGTCAGGA |
| sodA-Rv | GCAAGAACTCAGCCAGATGA |
| sodB-Fw | CCGTAATTGTGCAGGTCGTC |
| sodB-Rv | CTGCAAATCAACGATCGCCT |
| sodA-WT_NdeI Fw | TACTAAGGAGGTCATATGAATAGTTCTCGCAGC |
| sodA-EcoRI_ Rv | TTTCATTGCCATACGGAATTCAAGGCACAGGT |
| sodA-sol_NdeI Fw | TACTAAGGAGGTCATATGTACCCCGCTGAGTTAC |
| sodB-NdeI Fw | TACTAAGGAGGTCATATGGCCTTTGAACAGCCTC |
| sodB-EcoRI_ Rv | TCATTGCCATACGGAATTCAGGCCATCTTTGGTC |
| sodA6312- Fw | AAGGAGGTCAATGACAGACAAATCCTGTAGTCGCC |
| sodA6312- Rv | GTCTTTCATTGCCATACGGTTATCCCGTAGCTTGCTCA |
| sodAN- Fw | AAGGAGGTCATATGATGAATAGTTCTCGCAGCATCAG |
| sodAN- Rv | TCAAAGGCGTAGGCTCTCAGTTCACCTGTTGG |
| sodBC Fw | AGCCTACGCCTTTGAACAGCCTCCCT |
| sodBC Rv | CATTGCCATACGGAATTCTCAAGCCGCCAA |

**Table S3 :** Strain used for the phylogenetic analysis of cyanobacterial MnSODs in **Fig. 4** and the predicted localization of MnSODs according to SignalP 6.0.

| Strain | Accession | Classification (SignalP 6.0) |
|---|---|---|
| Nostocales cyanobacterium LEGE 11386 | MBE9051467.1 | Lipoprotein |
| Trichormus sp. ATA11-4-KO1 | MBW4558831.1 | Lipoprotein |
| Nodularia sp. UHCC 0506 | WP_323215458.1 | Lipoprotein |
| Myxacorys almedinensis | WP_162421941.1 | Lipoprotein |
| Halotia wernerae UHCC 0503 | MEA5507287.1 | Lipoprotein |
| Microcoleus sp. SVA1B4 | WP_445299659.1 | Lipoprotein |
| Dendronalium sp. ChiSLP03b | WP_322662701.1 | Lipoprotein |
| Nostoc minutum NIES-26 | RCJ28434.1 | Lipoprotein |
| Pantanalinema rosaneae | WP_431655370.1 | Lipoprotein |
| Pantanalinema sp. GBBB05 | WP_432812051.1 | Lipoprotein |
| Microcoleus sp. A003_D6 | WP_442927815.1 | Lipoprotein |
| Microcoleus sp. B4-C3 | WP_445299397.1 | Lipoprotein |
| Anabaena subtropica | WP_277875608.1 | Lipoprotein |
| Neowestiellopsis sp. NC-DW | MHN9116841.1 | Lipoprotein |
| Pantanalinema rosaneae CENA516 | MGI0483888.1 | Lipoprotein |
| Microcoleus vaginatus | WP_445312326.1 | Lipoprotein |
| Nostoc sp. B(2019) | NDJ23487.1 | Lipoprotein |
| Nodularia spumigena | WP_271918904.1 | Lipoprotein |
| Rivularia sp. MS3 | MBV6623186.1 | Lipoprotein |
| Nodularia spumigena | WP_271942563.1 | Lipoprotein |
| Nodularia sphaerocarpa | WP_272068383.1 | Lipoprotein |
| Dendronalium phyllosphericum | WP_214431594.1 | Lipoprotein |
| Pseudanabaenaceae cyanobacterium LEGE 13415 | MBE9012496.1 | Lipoprotein |
| Rivularia sp. UHCC 0363 | WP_323238745.1 | Lipoprotein |
| Kalymmatonema hyalinum WJT4-NPBG1 | MBW4498943.1 | Lipoprotein |
| Nodularia spumigena | WP_063871086.1 | Lipoprotein |
| Scytonema sp. PCC 10023 | WP_414585734.1 | Lipoprotein |
| Leptolyngbya sp. NIES-2104 | WP_059000726.1 | Lipoprotein |
| Scytonema sp. PRP1 | WP_332997608.1 | Lipoprotein |
| Nodularia spumigena | WP_006196478.1 | Lipoprotein |
| Fischerella thermalis CCMEE 5201 | PMB52031.1 | Lipoprotein |
| Halotia branconii | WP_281484005.1 | Lipoprotein |
| Trichormus variabilis NIES-23 | WP_010994247.1 | Lipoprotein |
| Anabaena subtropica FACHB-260 | MBD2346042.1 | Lipoprotein |
| Myxacorys californica WJT36-NPBG1 | MBW4417879.1 | Lipoprotein |
| Atlanticothrix silvestris | WP_214442418.1 | Lipoprotein |
| Kovacikia sp. CALU 1929 | WP_482030890.1 | Lipoprotein |
| Nostocales cyanobacterium 94392 | MEB3216192.1 | Lipoprotein |
| Nostoc parmelioides | WP_277884298.1 | Lipoprotein |
| Leptolyngbyaceae cyanobacterium M33_DOE_097 | HIK16311.1 | Lipoprotein |
| Chlorogloeopsis sp. ULAP02 | WP_341997750.1 | Lipoprotein |
| Goleter apudmare HA4340-LM2 | MBW4644872.1 | Lipoprotein |
| Calothrix parietina FACHB-288 | WP_190549236.1 | Lipoprotein |
| Nodularia sp. LEGE 06071 | WP_193998321.1 | Lipoprotein |
| Anabaena sp. CCY 9910 | WP_414756233.1 | Lipoprotein |
| Calothrix sp. FACHB-168 | WP_190629649.1 | Lipoprotein |
| Leptolyngbya sp. AN10 | WP_430678559.1 | Lipoprotein |
| Plectonema radiosum | WP_193917416.1 | Lipoprotein |
| Atlanticothrix silvestris | WP_318780343.1 | Lipoprotein |
| Nostoc sp. UHCC 0252 | WP_323228814.1 | Lipoprotein |
| Komarekiella delphini-convector | WP_191756852.1 | Lipoprotein |
| Nodularia sp. NIES-3585 | WP_089093562.1 | Lipoprotein |
| Calothrix membranacea FACHB-236 | MBD2168452.1 | Lipoprotein |
| Nostoc sp. CreGUA01 | MFN6514993.1 | Lipoprotein |
| Chlorogloeopsis sp. ULAP01 | WP_289789894.1 | Lipoprotein |
| Nostoc sp. UIC 10630 | WP_163931552.1 | Membrane |
| Myxacorys chilensis ATA2-1-KO14 | MBW4540880.1 | Lipoprotein |
| Nostoc sp. ATCC 53789 | WP_114081005.1 | Membrane |
| Chlorogloeopsis fritschii | WP_276752922.1 | Lipoprotein |
| Tolypothrix sp. FACHB-123 | WP_190751173.1 | Lipoprotein |
| Nostoc sp. LEGE 12447 | WP_322745234.1 | Membrane |
| Nostoc sp. NMS1 | WP_292704904.1 | Membrane |
| Nostoc sp. | WP_335036612.1 | Membrane |
| Nostoc sp. C110 | WP_392482408.1 | Membrane |
| Nostoc sp. MG11 | WP_193198009.1 | Lipoprotein |
| Nostoc punctiforme PCC 73102 | WP_012408308.1 | Membrane |
| Nostoc sp. FACHB-133 | WP_277883559.1 | Membrane |
| Calothrix sp. FACHB-156 | MBD2342527.1 | Lipoprotein |
| Nostoc sp. LEGE 12450 | WP_322746338.1 | Membrane |
| Chlorogloeopsis fritschii PCC 6912 | RUR73923.1 | Lipoprotein |
| Nostoc sp. TCL240-02 | WP_302480924.1 | Membrane |
| Chlorogloeopsis fritschii | WP_016872332.1 | Lipoprotein |
| Nostoc sp. UCD120 | WP_311774093.1 | Membrane |
| Brasilonema sp. UFV-L1 | WP_169219177.1 | Lipoprotein |
| Nostoc sp. PCC 7120 = FACHB-418 | RUR83944.1 | Lipoprotein |
| Nostoc sp. FACHB-110 | WP_242072232.1 | Lipoprotein |
| Nostocaceae | WP_011318269.1 | Lipoprotein |
| Anabaena sp. CCY 0017 | WP_414550770.1 | Lipoprotein |
| Nostoc sp. C057 | WP_306297076.1 | Membrane |
| Nostoc edaphicum | WP_181929818.1 | Lipoprotein |
| Scytonema sp. NUACC21 | WP_460206421.1 | Lipoprotein |
| Rivularia sp. T60_A2020_040 | MBF2015390.1 | Lipoprotein |
| Nostoc sp. CENA543 | WP_103138074.1 | Lipoprotein |
| Anabaena sp. CCY 9402-a | WP_414578860.1 | Lipoprotein |

| Strain | Accession | Classification (SignalP 6.0) |
|---|---|---|
| Anabaena sp. UHCC 0399 | WP_323267495.1 | Lipoprotein |
| Anabaena sp. UHCC 0253 | WP_254449390.1 | Lipoprotein |
| Anabaena cylindrica PCC 7122 | WP_015215174.1 | Lipoprotein |
| Nostoc sp. CALU 1950 | WP_435020268.1 | Membrane |
| Nostoc sp. CHAB 5824 | MCC5641606.1 | Membrane |
| Nostoc sp. FACHB-110 | MBD2440923.1 | Lipoprotein |
| Nostoc punctiforme PCC 73102 | ACC83785.1 | Lipoprotein |
| Cyanothece sp. PCC 7425 | WP_041237046.1 | Lipoprotein |
| Gloeobacter violaceus PCC 7421 | BAC88623.1 | Other |
| Pseudanabaena sp. PCC 7367 | WP_015163874.1 | Soluble |
| Escherichia coli | WP_248807151.1 | Soluble |
| Synechococcus sp. PCC 6312 | WP_172636034.1 | Soluble |
| Pseudocalidococcus azoricus | WP_322876523.1 | Soluble |
| Thermosynechococcus sp. HN-54 | WP_250682089.1 | Soluble |
| Mastigocoleus testarum | WP_027843320.1 | Lipoprotein |
| Thermosynechococcus sp. | WP_297761391.1 | Soluble |
| Pleurocapsa sp. MO_226.B13 | MDJ0592545.1 | Soluble |
| Candidatus Synechococcus calcipolaris | WP_277866136.1 | Soluble |
| Calothrix parietina FACHB-288 | WP_190544005.1 | Lipoprotein |
| Calothrix membranacea FACHB-236 | MBD2166265.1 | Soluble |
| Calothrix sp. FACHB-168 | WP_096643960.1 | Lipoprotein |
| Tolypothrix sp. FACHB-123 | WP_190743315.1 | Lipoprotein |
| Thermosynechococcus sp. TG252 | WP_310454098.1 | Soluble |
| Pyrinomonas methylaliphatogenes | MBX5479872.1 | Soluble |
| Thermosynechococcus sp. NK55a | WP_024125507.1 | Soluble |
| Thermosynechococcus sp. FA-CM-4201 | WP_460193008.1 | Soluble |
| Thermosynechococcus vestitus | WP_011055891.1 | Soluble |
| Nostoc sp. LPT | WP_292868329.1 | Lipoprotein |
| Thermosynechococcus sp. | WP_409873764.1 | Soluble |
| Nostoc sp. | WP_334926809.1 | Lipoprotein |
| Elainella sp. Prado103 | MCU0526905.1 | Soluble |
| Thermosynechococcus sp. M55_K2018_012 | WP_297053013.1 | Soluble |
| Tolypothrix sp. PCC 7601 | WP_045869977.1 | Lipoprotein |
| Thermosynechococcus sp. OHK43 | WP_365970883.1 | Soluble |
| Aerosakkonema funiforme | WP_374791410.1 | Lipoprotein |
| Aerosakkonema funiforme FACHB-1375 | WP_190465982.1 | Lipoprotein |
| Tolypothrix tenuis | WP_096582728.1 | Lipoprotein |
| Thermosynechococcus sp. B1 | WP_286627080.1 | Soluble |
| Thermosynechococcus sp. PP42 | WP_302677659.1 | Soluble |
| Thermosynechococcus sichuanensis | WP_181494764.1 | Soluble |
| Pyrinomonas methylaliphatogenes | WP_041973980.1 | Soluble |
| Thermosynechococcus sp. M98_K2018_005 | WP_297051732.1 | Soluble |
| Nostoc linckia FACHB-104 | MBD2208974.1 | Lipoprotein |
| Calothrix sp. FACHB-156 | MBD2339284.1 | Lipoprotein |
| Scytonema sp. PMC 1070.18 | MEC4881204.1 | Lipoprotein |
| Tolypothrix sp. PCC 7910 | WP_167724335.1 | Lipoprotein |
| Thermosynechococcus sp. QS41 | WP_310800931.1 | Soluble |
| Thermosynechococcus sp. GLH187 | WP_310821562.1 | Soluble |
| Calothrix sp. NIES-2100 | WP_096599871.1 | Lipoprotein |
| Nostoc sp. | WP_335056299.1 | Lipoprotein |
| Thermosynechococcus sp. HY213 | WP_206200084.1 | Soluble |
| Aerosakkonema sp. BLCC-F183 | WP_374825979.1 | Lipoprotein |
| Sphaerospermopsis kisseleviana | WP_096571931.1 | Lipoprotein |
| Leptodesmis sichuanensis | WP_428360810.1 | Soluble |
| Thermosynechococcus sp. PKX95 | WP_310743007.1 | Soluble |
| Sphaerospermopsis sp. FACHB-1094 | WP_190648403.1 | Lipoprotein |
| Aulosira sp. DedQUE10 | MDZ7959988.1 | Lipoprotein |
| Nostoc commune | WP_109007648.1 | Lipoprotein |
| Nostoc sp. GT001 | WP_289795776.1 | Lipoprotein |
| Anabaena sphaerica | WP_190560307.1 | Lipoprotein |
| Leptodesmis sichuanensis | WP_233744687.1 | Soluble |
| Nostoc sp. DedQUE05 | WP_322673311.1 | Lipoprotein |
| Trichocoleus sp. FACHB-69 | WP_190455393.1 | Soluble |
| Nostoc edaphicum CCNP1411 | QMS89058.1 | Lipoprotein |
| Nostoc sp. | WP_334931430.1 | Lipoprotein |
| Thermosynechococcus sp. Uc | WP_300872640.1 | Soluble |
| Sphaerospermopsis kisseleviana | MFM5980434.1 | Lipoprotein |
| Nostoc sp. 'Peltigera malacea cyanobiont' DB3992 | WP_099100092.1 | Lipoprotein |
| Thermosynechococcus sp. | WP_409869042.1 | Soluble |
| Leptodesmis sp. | WP_445926464.1 | Soluble |
| Calothrix sp. PCC 7716 | BDA72795.1 | Lipoprotein |
| Nostoc sp. DedSLP04 | WP_322689216.1 | Lipoprotein |
| Sphaerospermopsis sp. FACHB-1194 | WP_190347369.1 | Lipoprotein |
| Nostoc sp. | WP_335012934.1 | Lipoprotein |
| Nostoc sp. DedQUE08 | WP_322701324.1 | Lipoprotein |
| Nostoc sp. | WP_335201774.1 | Lipoprotein |
| Rivularia sp. IAM M-261 | GJD16125.1 | Lipoprotein |
| Trichocoleus sp. FACHB-832 | WP_190424133.1 | Soluble |
| Chroococcidiopsis thermalis PCC 7203 | WP_015152615.1 | Soluble |
| Chroococcidiopsis thermalis PCC 7203 | WP_015156211.1 | Other |

**Table S4 :**

Doubling time of the WT and the Δsod mutants grown under low, standard or high light. Results represent mean ± stdev from 3 independent experiments.

| **Strain** | **Doubling time (h)** | **Standard Deviation** |
|---|---|---|
| Low-Light (600 lux) | | |
| **WT** | 32 | 0.6 |
| ***ΔsodA*** | 34 | 0.1 |
| ***ΔsodB*** | 42 | 0.6 |
| Standard-Light (2500 lux) | | |
| **WT** | 26 | 0.1 |
| ***ΔsodA*** | 28 | 0.3 |
| ***ΔsodB*** | 29 | 0.7 |
| High-Light (7000 lux) | | |
| **WT** | 21 | 0.1 |
| ***ΔsodA*** | 25 | 2.1 |
| ***ΔsodB*** | 31 | 1.1 |
